# De novo homology assessment from landmark data: A workflow to identify and track segmented structures in plant time series images

**DOI:** 10.1101/2021.02.21.432162

**Authors:** John G. Hodge, Qing Li, Andrew N. Doust

## Abstract

Assessing the phenotypes underlying plant growth and development is integral to exploring the development, genetics, and evolution of morphology and plays an essential role in agronomic and basic research studies. Although various automated or semi-automated phenomic approaches have recently been developed, tools assessing differential growth of plant organs remains a key topic of interest, but one which is often difficult to analyze due to the requirements of segmenting and annotating specific structures or positions in the plant body in time-series data. To address this gap, we have developed a generalized workflow linking our previously published function, *acute*, with a companion function, *homology*, in the PlantCV environment. The *homology* function uses a generalized strategy of dimensionality reduction via *starscape* followed by hierarchical clustering through *constella* to identify ‘constellations’ of segments in eigenspace that represent the same landmark in consecutive images of a time-series. We devised a quality control function, *constellaQC*, that can test the accuracy of the clustering approach, and we use it to show that the approach accurately clustered the pseudo-landmarks derived from *acute*, although with several sources of error. We discuss the reasons for and consequences of these errors in automated workflows, and suggest how to develop these functions so that they can easily be repurposed for other phenomics datasets that may vary in dimensional complexity.

## Introduction

Phenomics methodologies are revolutionizing studies of plant morphology, as the use of high-throughput digital imaging techniques provides a level of objective trait measurement that can far surpass the available time and accuracy of manual observers. Both holistic and component approaches (Das Choudhury et al. 2018) have been proposed and used for specific studies, with holistic methods treating the whole plant as a single object, such as the use of convex hulls of 2D images to approximate biomass in grasses and rosette size in Arabidopsis (Fahlgren, et al. 2015, Dobrescu et al. 2017, Green et al. 2012, Minervini et al. 2017), or component, measuring individual organs (McCormick et al. 2016, Miao et al. 2020, Pound et al. 2017). A somewhat different approach, that of persistent homology, has been used to score both size and shape attributes at different scales within the plant (Li, et al. 2016, Li, et al. 2018). Measurements can be made on individual images or on time series, but a particular issue for most phenotyping methods for time-series data is how to track organs through time, i.e. how to identify the same organ or other landmark through consecutive images of a time-series. Accurately specifying the same segmented structure, such as the tip of a leaf, through multiple images is essential for understanding the behavior of that structure and presents many challenges. In most applications, these challenges have been overcome through manual identification, curation, and annotation, which presents a significant time burden (e.g. McCormick et al. 2016). Other approaches have relied on the rapidly developing methodologies of machine learning (e.g. Pound et al. 2017), but these require large training sets and manual validation for their success. We are interested to find solutions that bridge the divide between full manual annotation and machine learning approaches, and here we present a workflow which automates much of the annotation of structures through time and creates data sets that can be easily curated for further analysis.

A general purpose, fast, and tunable method of capturing landmarks and tracking them through development that could be applied to multiple plant imaging needs would be a valuable starting point. We have previously published a landmarking method called *acute* in the PlantCV package (Gehan et al. 2017) and here present a detailed explanation and test of that method, along with an extension, *homology*, that automates linkage of homology groups of landmarks that represent the same organ through multiple time-series images. We investigate the accuracy and biases of the automated homology assessment on a set of trait data generated for a subset of an existing recombinant inbred line (RIL) mapping population of a cross between *Setaria italica* (foxtail millet) and its wild progenitor *S. viridis* (green foxtail), selecting exemplar individuals that are morphotypic extremes of the population (Mauro-Herrera and Doust 2016, Feldman et al. 2017).

## Materials and Methods

### Plant Growth and Image Collection

Germplasm from the parental lines A10.1 (here referred to as A10) and B100 were used alongside 3 RILs, RIL39, RIL110, and RIL159 (Mauro-Herrera et al. 2013). RILs were selected to represent genotypes with distinct growth forms, based on morphometric studies of previous growth trial data (Mauro-Herrera et al. 2016). Plants were grown in 3” pots in Metromix 360 media (Sun Gro Horticulture, Agawam, MA) in Percival E36L growth chambers (Percival, Perry, Iowa) using a program set to maintain 12hr light/12hr dark cycles at 28°C and 22°C temperatures respectively, with humidity kept at approximately 50-60%. Plants were watered 2-3 times a week as needed, with imaging being performed daily until either heading or 30 days of growth occurred. A raspberry pi 3 with a 3-megapixel camera was used for plant imaging, with the camera mounted 1m away from a stationary point where plants were positioned against a high contrast white background. The raspistill function was used with the following settings sharpness=80, quality=100, timeout=12000 resulting in the generation of 1944 x 2592 jpeg output images. Plants were repositioned at the same angle each successive day unless torsion of the main culm was noted, in which case the fixed angle for imaging was readjusted so that the primary axes leaves were orthogonal to the camera. Prior to analysis, data images were thresholded using the *a* and *b* color channels in PlantCV, and subsequently merged into a joined binary mask. In some cases, manual adjustments were made to binary masks in GIMP following thresholding due to the belt-like shape of grass leaves which can allow them to become disjointed masks at regions where their blades are near parallel with the camera. Contour arrays were then generated using the OpenCV function *findContours* in order to capture the shape of the plant. Following these adjustments binary masks were then analyzed using our pseudo-landmarking analysis workflow, *acute*, followed by grouping of homologous pseudo-landmarks (p-lms) using the *homology* functions comprised of *space, starscape*, and *constella*. Assessment of the accuracy of the *acute* and *homology* pipelines was assessed through manual measurement and annotation of the time-series images.

### Overview of pseudo-landmark (p-lm) analysis workflow

The semi-automated landmark analysis workflow requires multiple steps, as outlined in Figure 1, and detailed in the jupyter files in Appendix 1. A preliminary step is to mask images in order to threshold the plant shape out of the background and generate a binary mask. This mask is subsequently used to generate p-lms with the PlantCV function *acute*. With time series data of sufficient resolution, p-lms can be assigned identities utilizing the *constella* pipeline, which clusters these morphological features into de novo homology groups. After homology group assignment, manual curation can be used to assign identities to features, which gives them biological meaning to enable their use in downstream comparative analyses.

**Figure 1.**
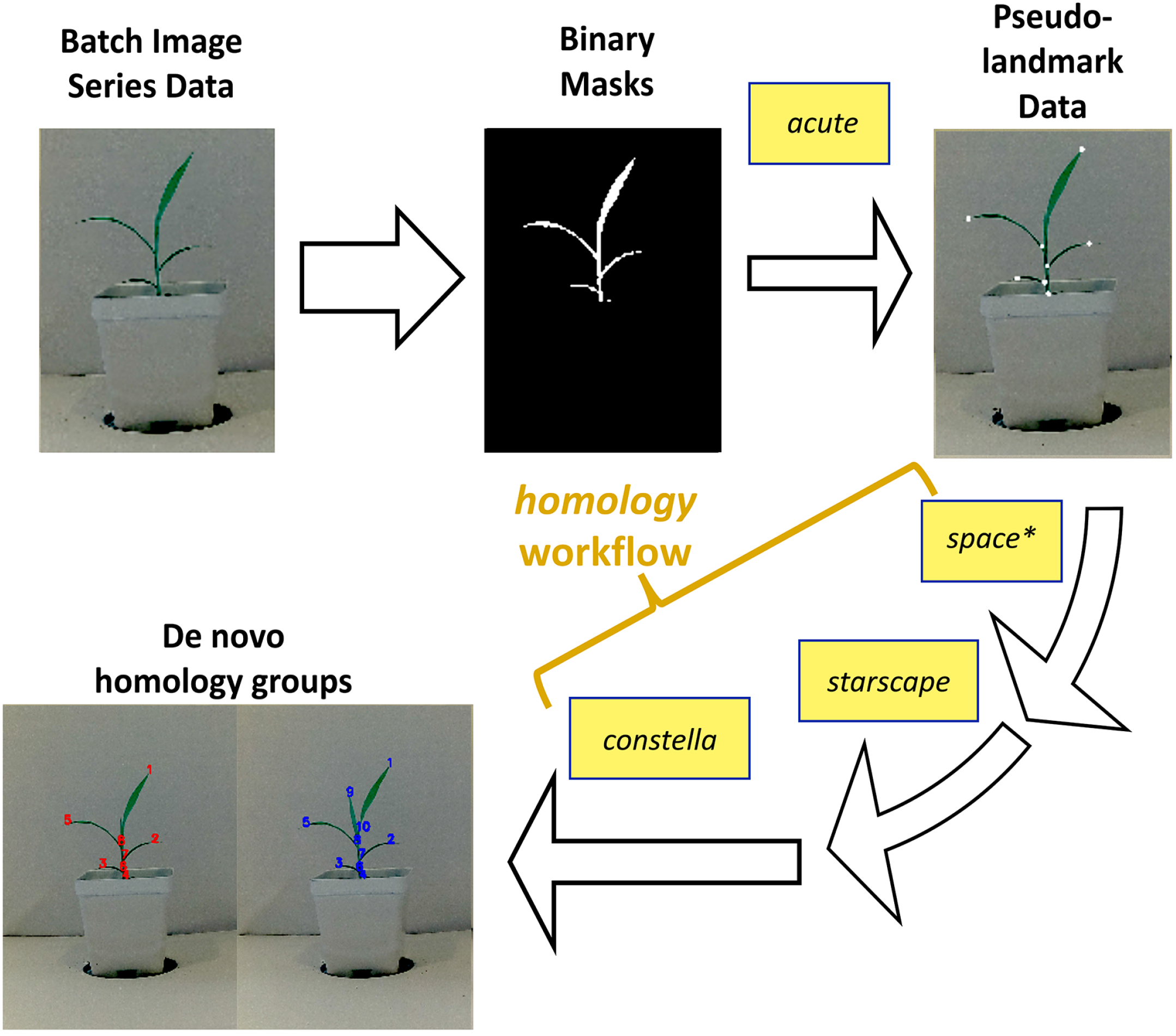
General outline of the *acute* pseudo-landmark analysis workflow from batch image data and the resulting grouping of de novo identities for time series data provided by the *homology* workflow. *Homology* itself operates using predominantly 2 functions *starscape* and *constella* with an optional function *space* (asterisk) that can be used prior to these other functions.

### I. Acute

*Acute* is a function written in python, built within the PlantCV environment, that is used for morphometric analysis of image data. This is done by isolating p-lms along the contour of a plant by identifying acute regions within a contour with an approach akin to chain coding. As previously described, OpenCV contour arrays serve as a template for generating these p-lm calls. The various input and output parameters of *acute* and an overview for its optimization steps are discussed below. *Acute* requires 5 parameters in order to generate p-lms for an image that are denoted within the function as *obj, mask, win, thresh*, and *debug*. Of these parameters, two are the direct outputs of OpenCV itself, *obj*, which is an OpenCV contour array generated from the binary image matrix, *mask*. The parameter *win* fine-tunes the generation of angle scores along the length of the contour array, *obj*, and the parameter *thresh* identifies acute-angled p-lm regions based on the angle scores along this contour. The importance of proper *win* and *thresh* specification for p-lm designation and factors affecting their behavior will be discussed below. Finally, the *debug* parameter designates whether verbose outputs should be produced during troubleshooting of the development of new p-lm workflows.

### Properties of the *obj* and *mask* inputs

The parameters *obj* and *mask* are both objects derived from native OpenCV functions. The object, *mask*, is the first to be generated and contains a binary mask derived from the OpenCV *threshold* function that captures the shape of the plant within an image. Although not necessary for generating p-lms per se, the inclusion of *mask* generates a helpful layer of metadata used in parsing downstream p-lms by generating convexity-concavity values based on mask pixel intensity. The two-dimensional array *obj* is an OpenCV contour array generated by *findContours* when *mask* is used as an input. This contour array is the primary input upon which *acute* generates p-lms based on the other specifying parameters *win* and *thresh*.

### Properties of the *win* parameter

The *win* parameter plays an integral role in the designation of p-lms as it provides a sliding window size by which angles can be computed along the contour. This is accomplished by using win to select 3 points from which an angle can be calculated. The selection of these three points begins with the specification of a vertex position, *k*, for which the angle score will be generated. From this middle point a conditional loop then begins which follows the general logic of the pseudo-code below in which the reverse coordinate, *A*, is obtained by migrating backwards along the contour until a vertex is found that is the largest distance from *k* while still being less than *win*. Forward coordinate, B, conversely is obtain by migrating forward along the contour while adopting a similar strategy. The final two points are ultimately selected by coordinates whose absolute distance from *k* is less than or equal to *win*. After points *k, A* and *B* are determined an angle is computed using the following law of cosines equations in order to define an output angle score given *k*, defined as *a_k_*. This is first done by specifying 3 distance values *P_Ak_*, *P_Bk_*, and *P_AB_*:

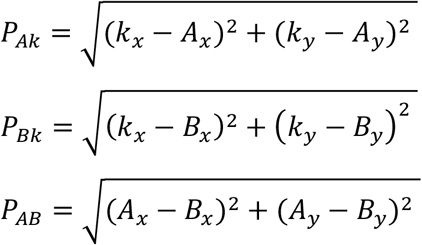

From these distances the arccosine of their dot product can then be used to solve for the angle score, *a_k_*, as:

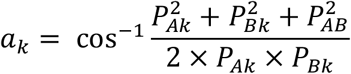

The resulting *a_k_* scores can subsequently be stored for the entire contour as a vector, *chain*, which can then be used to optimize and select an informative *win* value as this must often be empirically determined for each application. These steps will be described in detail in the output parameters section below.

### Properties of the *thresh* parameter

The *thresh* parameter is used to interpret the output that the chain parameter generated by *obj* and *win*. This is done by identifying islands of consecutively acute coordinates on contours which are merged together as representative of a single structure. The resulting vertex positions obtained from the thresholding chain can then be used to designate p-lms within this contiguous island (Fig. 2). Following thresholding, the island vertices are then reduced down to a small subset of points based on specific criteria. First, is an implicit assumption that only a single p-lm is required for islands of conterminous index positions along a contour (Fig. 2). When multiple points are present that are immediately neighboring one another, the best approximation of a landmark position, *LA*, is selected based on being the maximum distance between the starting site, SC, of the island and the terminal site, *T_s_*, of the island, based on the following formula:

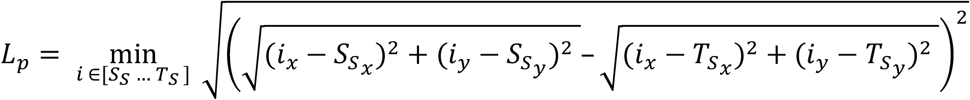

**Figure 2.**
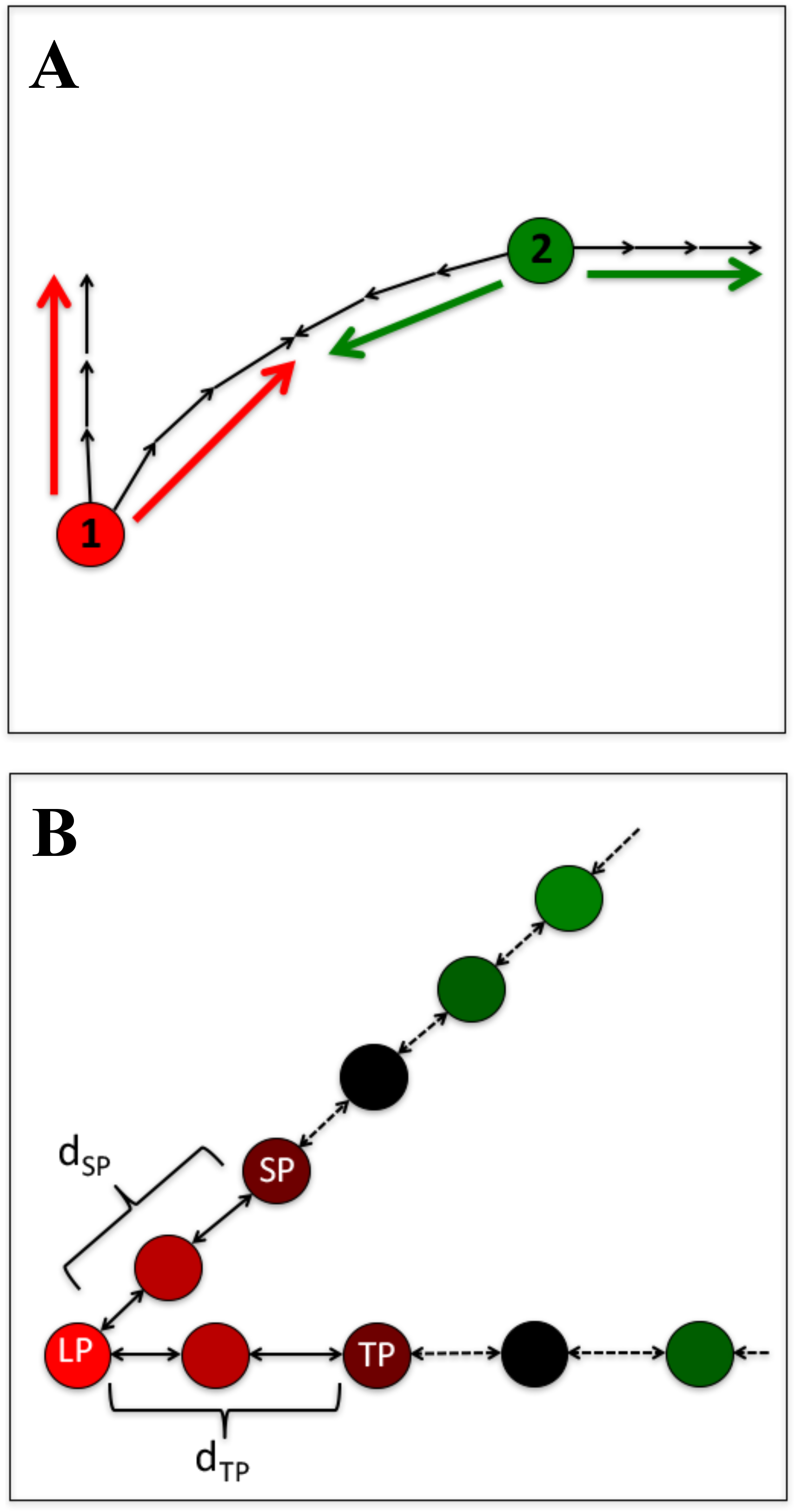
Demonstration of *acute* pseudo-landmark (p-lm) assignments. (A) Illustration demonstrating the methodology of assigning angle scores for vertices along a contour by measuring the angles at each vertex given a sliding window (3 steps in this case) which can produce an acute score for vertex 1, which differs from the obtuse score which would be derived for vertex 2. (B) Following the generation of angle scores across a contour, acute islands can be identified that represent a conterminous series of acute vertices, from which a starting site (SS) and termination site (TS) can be specified along its edges, and a pseudo-landmark which is equidistant between them (LP) can be defined.

On the occasions that there are either isolated or pairs of conterminous points the solitary position or the more acute point of the pair or solitary point is stored as a p-lm respectively. When beginning an analysis, an initial *thresh* value of 80-90 is generally a good default value for retrieving acute regions.

### Output parameters

There are several different output parameters returned from *acute* when it is run in debugging mode. These include *homolog_pts*, *start_pts*, *stop_pts*, *pt_vals*, *chain*, and *max_dist*. When *acute* is run with debugging mode off only *homolog_pts* will be returned in order to simplify outputs. The first three outputs are all abridged opencv contour arrays made up of x-y pixel coordinates with *homolog_pt*s containing the p-lm coordinates for the image, and *start_pt*s and *end_pts* consisting of the reverse and forward positions initially used in the angle score calculation of the p-lm. When these three pixels are used in conjunction they can infer the direction of the acute island which is used in downstream methods to link landmarks between samples or time points. The *pt_vals* output provides an average pixel intensity value within each acute island along the contour in order to delineate concave and convex areas as a “convexity-concavity” or “CC” score that varies between 0 and 255. Pixel colors in using a binary mask can be viewed as defining variables where 0 (i.e. black background) defines ligules or leaf axils and 255 (i.e. white object) defines leaf tips. The CC score can then be used as a ratio which will operate within the confines of this range as serve as a reasonable bimodal predictor about the direction each p-lm is facing along the contour. The *chain* output is an array containing the angle scores generated using the *win* input parameter and is used in the optimization and selection of an effective sliding window size. Lastly verbose directs *acute* to print verbose outputs dynamically while running for use in troubleshooting and diagnosing behaviors while running this function.

### *Acute* workflow and optimization

Given that the *acute* function is versatile and can be used on a wide range of image data sets, optimization steps must be taken in order to successfully generate informative p-lms for downstream analyses. The most critical step is the selection of an appropriate *win* value as this determines the radial window size needed to generate angle scores. Moreover, the manner in which angle scores are computed with *win* may vary between imaging systems, as the parameter is sensitive to changes in resolution. A good best practice is to find the shortest distance between two landmark regions of interest in an image dataset and use half of this pixel distance as the initial *win* value. For example, within our growth experiment dataset the half of the average linear distance between the first leaf tip and the ligule of the first leaf of a representative set of images is used as a *win* value. Selection of a *thresh* value is simpler, but is equally as important, because it is used as a threshold to define acute p-lm regions within a contour. Initial *thresh* values around 80 to 90 are often appropriate, although more stringent values can be used if deemed appropriate. When selecting *thresh* it can sometimes be useful to look at the chain waveform output graphs in which the angle score of each contour point within an image is plotted based on its position within the contour (Fig. 3). When viewing these graphs, acute regions can be recognized as sharp valleys within the line plot and *thresh* values can be selected that capture the bases of these valleys while reducing noise.

**Figure 3.**
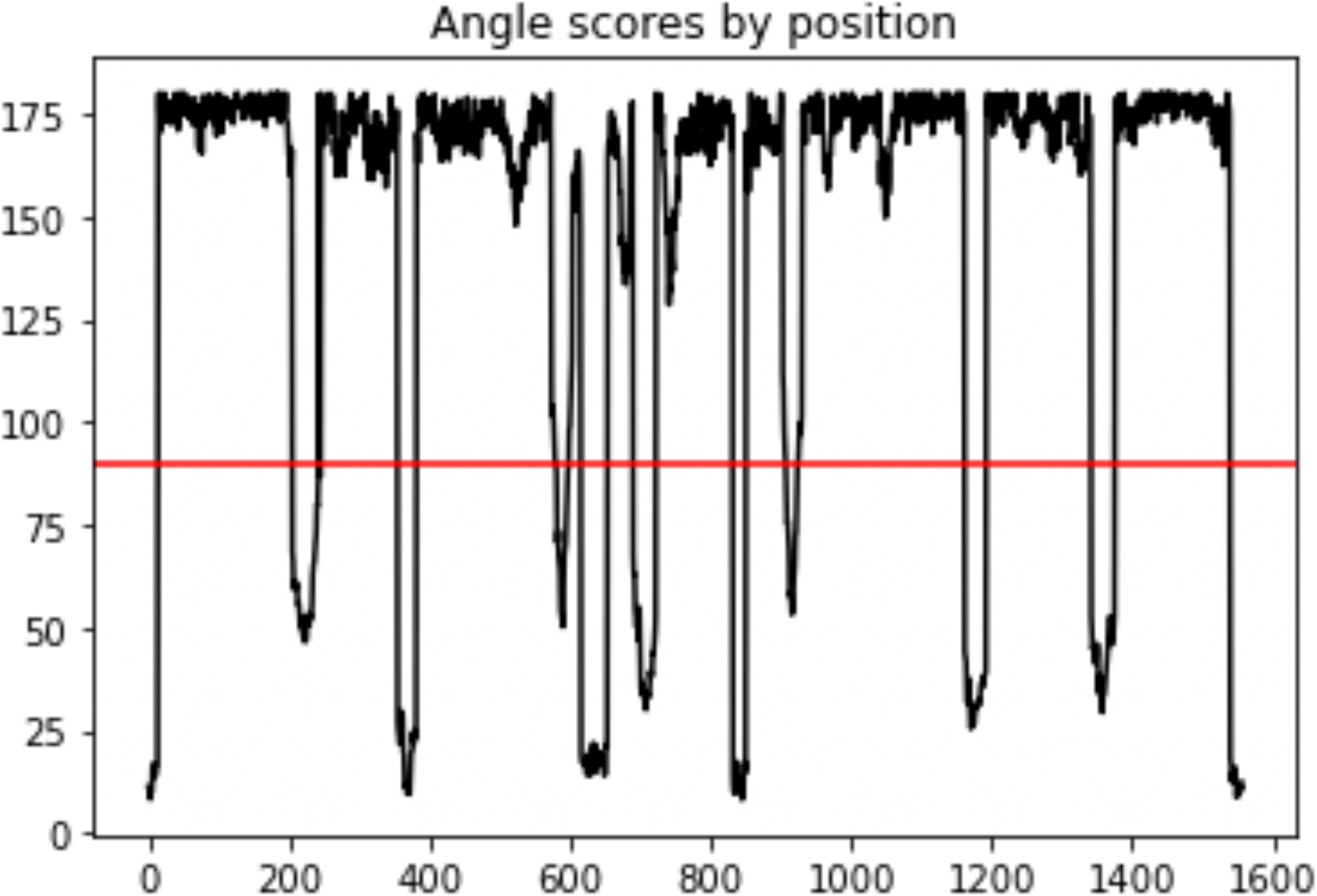
Chain output demonstrating *acute* angle scores of all vertices along a contour. The acute islands extracted for generating p-lm assignments are the regions of this contour which fall below the threshold of 90° (red line).

### Assessment of accuracy of the *acute* function

Following accurate annotation, it is possible to assess how well measurements made from the curated dataset correspond to other methods of measurement. We took a simple approach of comparing the measurements of height derived from the difference between the topmost ligule p-lm and the base of the plant in the last image of each time-series to measurements from the same images using the distance function in ImageJ.

### II. The *homology* workflow: grouping through de novo landmark linkage

Landmark linkage through our de novo homology grouping pipeline is specifically designed for tracking orthologous p-lms through time course data, enabling quantitative measurement and providing high resolution tracking of the ontogeny of growth of specific organs. This method was designed in such a way as to mimic object permanence, the idea that there is an understanding that develops in early childhood that a recently viewed object still exists, even when it is out of sight (Piaget 1954). By analogy, the homology workflow judges p-lms in consecutive images that remain physically close in space to most likely represent the same structure at two consecutive time-points. *Homology* operates through the use of three functions that operate sequentially to, first, generate metadata from *acute* outputs for use in homology group estimation with *space*, second, utilize a principal component analysis to extract maximally informative axes for estimating relationships between p-lms in the multivariate metadata space through the use of *starscape*, and third, utilize hierarchical clustering to estimate distances between points, from which homology groups are assigned using an iterative nearest neighbor joining strategy with *constella*.

### The *space* function

Given the lack of precedents for designing a naive homology grouping approach, we wanted to know whether developing tools which expanded the meta-information used in clustering homologous structures would prove useful. The space function was originally designed with this issue in mind by serving to provide some additional dimensions derived from the base *acute* outputs that might improve grouping accuracy. For each pair of neighboring image frames in a time series the *space* function operates by taking a series of informative attributes initially generated by *acute* and expanding them into a series of additional positional or directional attributes that maximize the differences between p-lms. Positional attributes first begin by generating a bounding box around the p-lms for the image pair to identify corners as well as a ‘center of mass’ centroid in which the mean p-lm x and y coordinates are used to define a mid-point. From each of these five points a Euclidean distance is calculated for the p-lm to describe its position in space relative to other p-lms being considered. Directional attributes operate by calculating degree angles to describe the orientation of p-lms within this space. To define the orientation of each p-lm as the direction it is facing on a 2-dimensional surface the 3 coordinates describing each p-lm (i.e. the landmark itself as well as the starting position, *S_p_*, and termination position, *T_p_*, which were generated during p-lm assignment) are used by first estimating a mid-point between the bounding starting and termination coordinates and driving a line towards the p-lm coordinate to generate a slope. The slope, *m*, resulting from the line generated can then be converted to an angle in degrees through the use of the following formula:

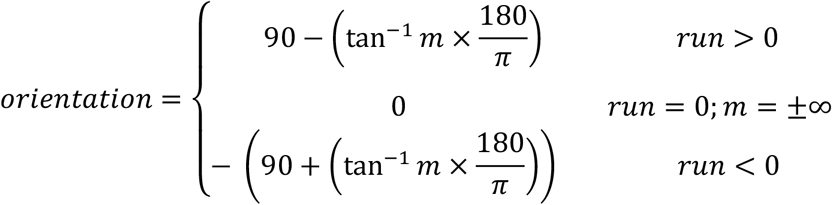

Angles are subtracted against the quadrant weight of 90 so that leaf orientation describes deviance from a vertical plane, where negative numbers describe increasing left skew and increasing positive numbers describe increasing right skew.

### The *starscape* function

The combined p-lms from two sequential images along with the meta-variables generated from are subjected to PCA analysis, and eigenvalues larger than 1 are extracted and stored in an output dataframe for use in homology grouping. In addition to these principal component values for p-lms, the eigenvalues used to generate the scree plot output (Fig. 4A) and the loadings of the informative components onto the original dimensions are also provided as outputs. A scatterplot of the first three principal component axes with p-lms of the two neighboring frames color coded to allow for easy cross comparison is also generated (Fig. 4B).

**Figure 4.**
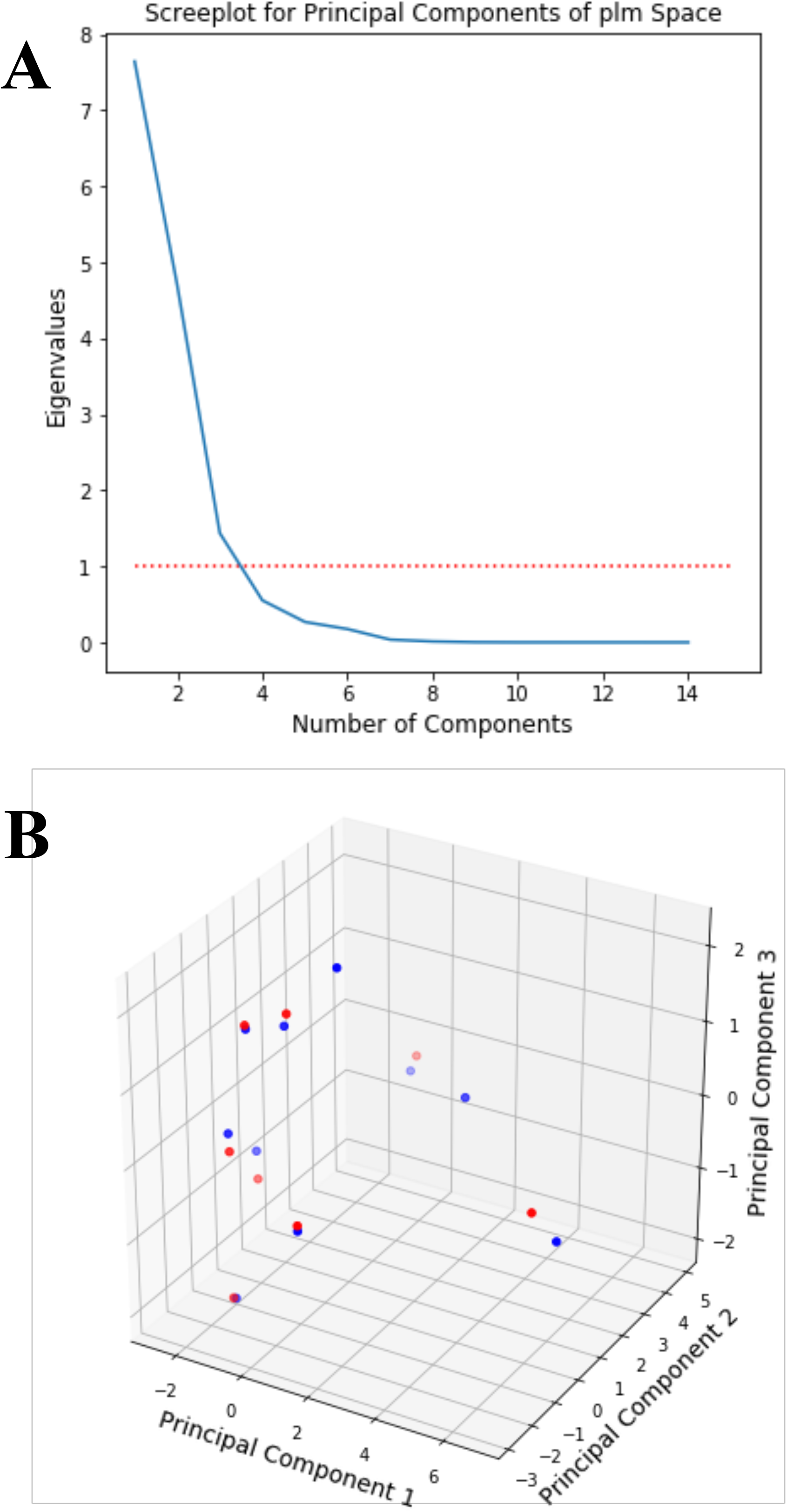
Graphical outputs resulting from *starscape* function. (A) Scree plot of eigenvalues for principal components. (B) Plot of first three components with neighboring images color coded in red and blue respectively.

### The *constella* function

Following the generation of the informative principal component axes, hierarchical clustering is used to identify close neighbors, which are inferred to be the same organ in the different images, and so are classified into homology groups within the constellation of the multivariate ‘starscape’ (Fig. 5), while also having the flexibility to identify novel features as they appear. This step is conducted by *constella* using the original *space* multivariate dataframe and the dataframe of informative PC axes generated by *starscape*, in addition to a group iterator seed number which will serially assign unique serial numbers to homology groups from this integer onward.

**Figure 5.**
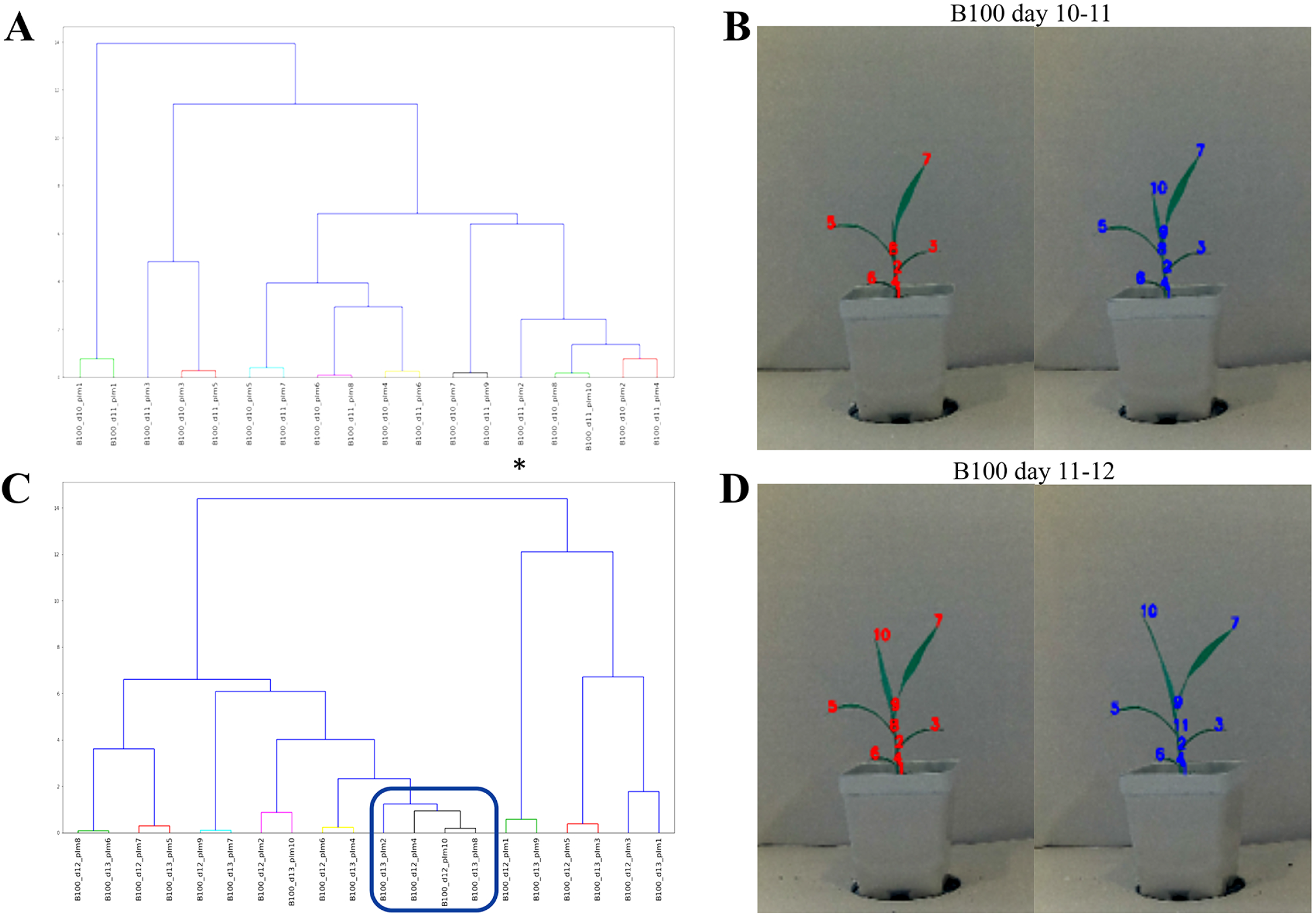
Graphical outputs resulting from *constella* function (A & C) as well as these resulting homology group ID numbers superimposed onto original images (B & D). Note that the majority of p-lms form duets or pairs of points with high fidelity, however rogue points (A-asterisk) occasionally arise when new structures are formed. In addition, quartets (C-box) may sometimes occur when structures are still migrating and have a closer relationship with their nearest duet pair than they do with each other.

Closest neighbors within the *starscape* point cloud are defined through a nested neighbor joining like strategy in order to associate p-lms which are likely homologous based on their close proximity, and thus can be labeled the same structure in sequential images. This agglomerative strategy expands to capture nearest neighbor pairs as clusters and link them under a shared group identity number. Homology group pairs are of two distinct types, ‘duets’ and ‘quartets’. Duets, a single pair of p-lms in neighboring time points that form a cluster of two, are the simplest and most common type of cluster to generate assignments, and describe many structures that are relatively stationary after they have finished developing (Figs. 5A, 5B). Quartets, by contrast, are when two p-lms form a grade around a duet pair resulting in a comb of four p-lms in a cluster (Figs. 5C, 5D-box). This grade is often the result of capturing a structure that is either rapidly developing or is shifting its position relative to other structures within the body plan. This inflates the branch length of these newer p-lms, driving them out of a close association with their true neighbor pair (Fig. 5D group 10). So long as the initial duet that a quartet pair has formed around is annotated properly (the agglomerative strategy of walking back into the depth of the dendrogram is designed to promote this) quartet pairs can generally also be recovered with little difficulty. Following the identification of all clear pairs of points, assignments of what are deemed ‘rogue’ points can then begin (Fig. 5A-asterisk). As their name suggests these points are solitary and lack any discernible neighbors to pair with. This could be the result of two different phenomena, either these points could essentially be chaff resulting from parallax or other confounding events which produce acute regions and thus p-lms spontaneously for a transient period in time, or these points can also be the initial recording of a new structure as it appears in a plant’s ontogeny. In time-series datasets, the latter can often readily be distinguished from the former because linkage groups of p-lms begin to form from seeded rogue points over the course of development.

### III. *constellaQC* for landmark linkage error assessment

Following linkage assignments it is important to validate the overall accuracy for de novo calling. The first step is to manually curate a reduced list of samples in order to rename and correct for erroneously linked samples. This sample dataset can then act as a standard by which known features of interest to an investigator, such as leaf tips, can be compared to the predicted values through the use of *constellaQC*, in order to assess the accuracy of the de novo linkage method. Calculation of landmark linkage accuracy is conducted by assessing two different forms of error, splitting errors, in which features of interest are fractionated across multiple homology groups, and clumping errors in which a single homology group is mistakenly assigned to more than one discrete feature of interest. The means by which these two errors can be assessed is done by forming a pairwise matrix of known homology features compared against predicted homology groups (Fig. 6). In this scoring matrix the sum of all frames in time where a known (*k*) or predicted (*p*) homology group occurs is first summed into Σ_*k*_{1…*i*}__ and Σ_*p*_{1…*j*}__ respectively. In an ideal setting for each discrete feature there will only be one predicted homology group set so that Σ_*k*_{1…}__ = Σ_*p*_{1…*j*}__ and the number of valid linkage events between these homology groups could be defined as:

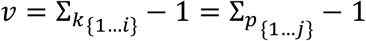

**Figure 6.**
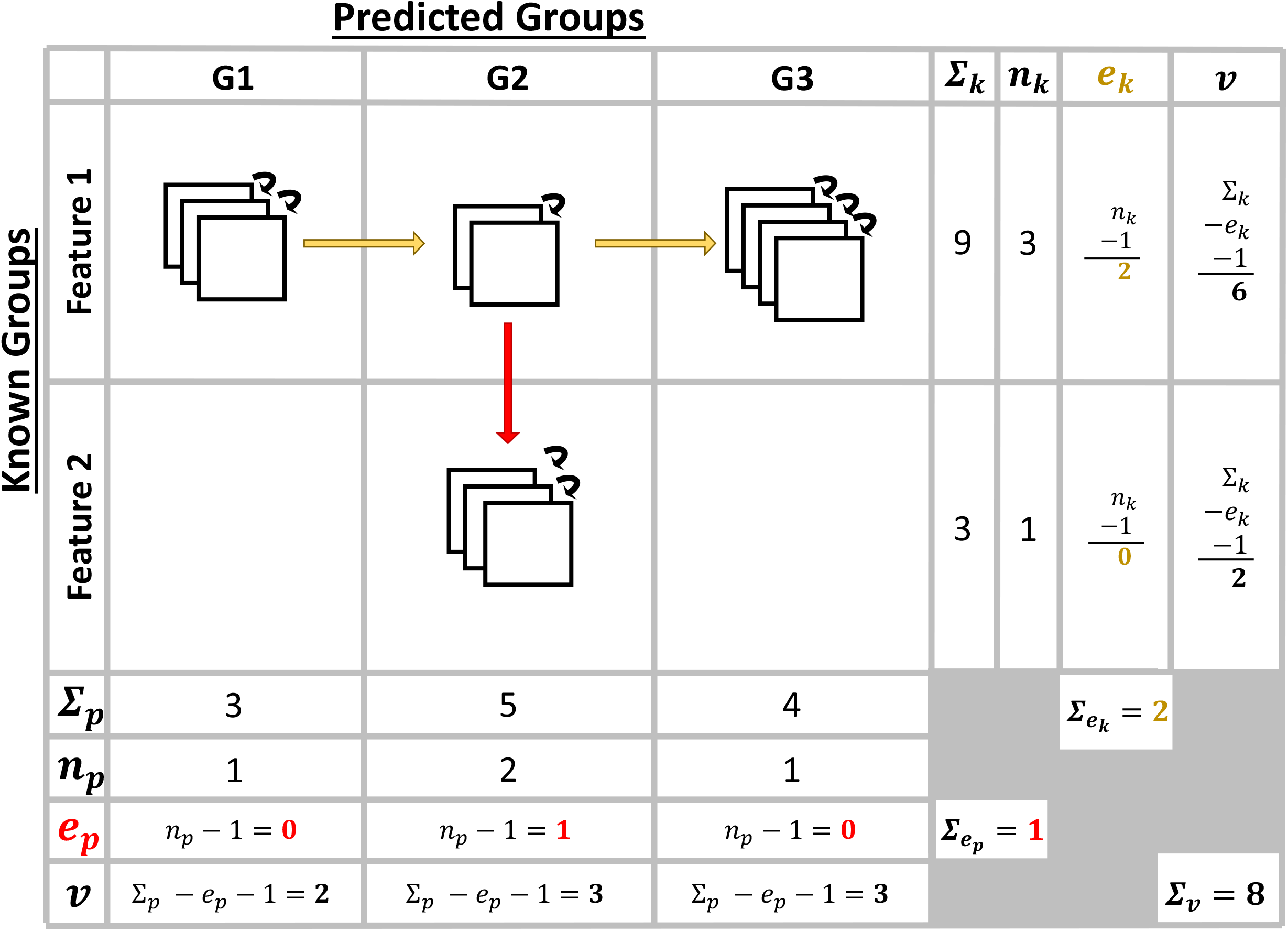
Defining the idealized method for scoring landmark linkage accuracy through use of a pairwise matrix to estimate valid calls (*v*) with respect to known features and predicted homology groups while also defining splitting error (*e_k_*) and clumping error (*e_p_*). Squares represent images through time with black arrows being valid linkage calls, yellow arrows being splitting errors, and red arrows being clumping errors.

However, in cases where an error occurs, the valid calls between these known and predicted groups may decay as erroneous linkage assignments are made. The overall number of erroneous calls can be defined as the disparity in numbers of known to predicted groups (i.e. *n*_*k*_{1…*j*}__ ≠ *n*_*p*_{1…*j*}__). In this way, an event where splitting errors occurred, and 3 groups were needed to describe the ontogeny of one known feature, the erroneous calling events would be defined as:

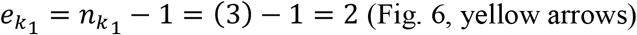

Reciprocally, a single clumping error in which two known homology groups are misidentified as the same predicted feature would be defined as:

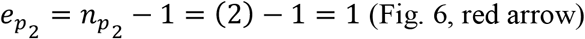

The valid known or predicted calls can then be tabulated once these sources of error are known as:

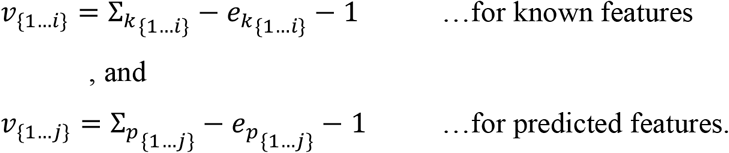

Finally, the summation of the known or predicted valid calls should be equal to one another as (Σ_*v*_).

Following these calculations for each ontogeny, overall landmark linkage accuracy can be estimated by three percentages, the valid call rate, the splitting call rate, and the clumping call rate. These three measures can be defined respectively in the context of Figure 6 as:

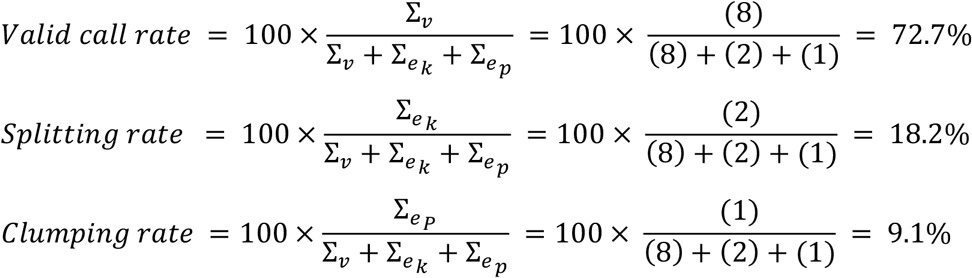

Of these measures, priority is usually given to the first and third values because valid calls represent the true accuracy of the landmark linkage method while clumping errors in conflating two distinct structures is the more serious error. As such efforts we aimed to maximize valid calls while minimizing clumping errors.

## Results

### *Acute* measurement accuracy compared to manual measures

To assess the reliability of phenotypic measures derived from *acute* p-lms, a side-by-side comparison with companion ground truth measures was conducted. Plant height based on the distance from the base of each plant to its highest ligule was used because ligules (which correspond to leaf axils in our images) are readily defined by *acute* and because this type of measure is commonly used in genetic and physiological research. We would expect a 1:1 ratio between manually curated (standard) heights plotted against p-lm estimates if *acute* correlates perfectly with the manual measurement. This 1:1 relationship would then result in a slope equal to 1 if a linear regression was fitted to these estimates and thus deviations in this slope could then be expected to reflect a tendency of *acute* to either over-estimate or under-estimate phenotypic measures based on the placement of these p-lms. Moreover, the R-squared value of this linear regression could provide a means to assess the degree to which these measures correlate with one another.

When these results are plotted out for each genotype (Fig. 7A,C,E,G,I) there is a strong correlation between manual and p-lm estimated heights (R-squared values > 0.9). Within genotypes B100, RIL110 and RIL159, there was a slight tendency of *acute* to over-estimate height, with the shallower slope of this regression likely due to the leaf axil being pushed slightly higher than the true ligule because the wide blades of these genotypes are inserted at an acute angle to the culm. The A10 and RIL39 genotypes more closely approximate a 1:1 relationship between the height estimates and their manual counterparts, which may be due to the larger insertion angle and narrower leaves of these genotypes.

**Figure 7.**
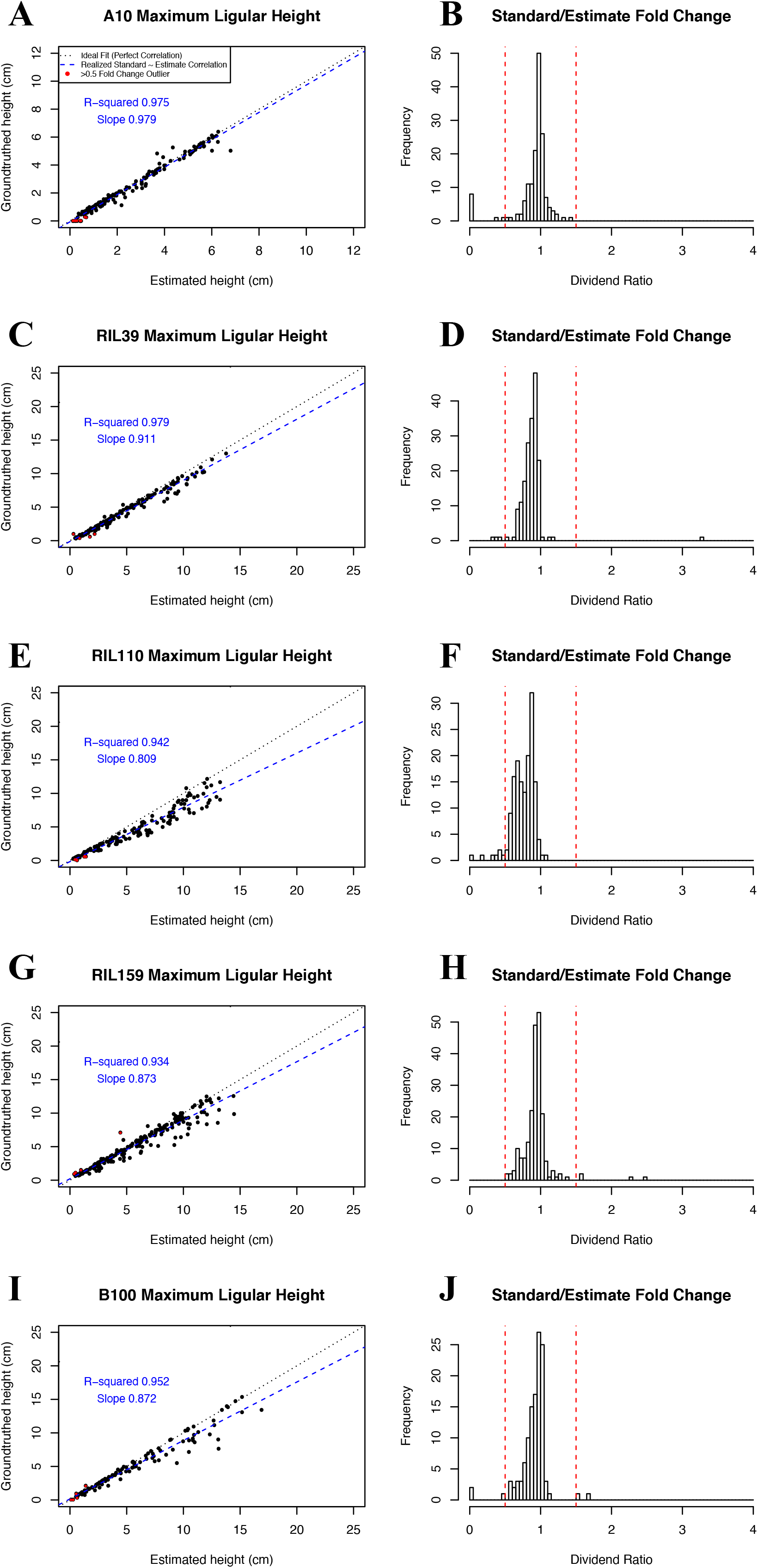
Validating the accuracy of the *acute* p-lm workflow for generating measures which approximate features of biological interest such as culm height. Regressions of *acute* derived culm heights between the highest ligule position and the base of the plant for genotypes A10 (A), RIL39 (C), RIL110 (E), RIL159 (G), and B100 (I) where the black dotted line represents an ideal 1:1 relationship and the blue dashed line represents the realized correlation between the manual and automated measures. The corresponding dividend ratio generated by dividing the manually generated measurements by the *acute* estimates of A10 (B), RIL39 (D), RIL110 (F), RIL159 (H), and B100 (J) reveals that the majority of these measures are centered around 1.

Another means to approximate the accuracy of this fit, since multiple points could potentially overlap in the same X-Y coordinate position, is to assay the fold change by dividing the ground truth measures by the p-lm estimates. The central tendency of such a ratio should center around 1 if these measures correspond to one another (Fig. 7B,D,F,H,J). A clear unimodal relationship can be seen in all cases with a slight left skew being visible in B100 and RIL110 because the insertion position of their leaf axils are estimated slightly higher than their ground truth counterparts (Fig. 7F, J). Overall, these results confirm that *acute* derived measurements are accurately approximating manually generated phenotypic measures.

### Factor Loadings from *starscape* following dimensionality reduction

Dimensionality reduction and reorientation was performed through *starscape* on data sets that included only the Euclidean distances between p-lms, but those created in *space* that also included distances from the bounding box and centroid coordinates, and those that included the orientation of the p-lms. *Starscape* works by dynamically extracting the minimum number of informative components (eigenvalues > 1) which in all cases was found to be the first three principle component (PC) axes. Although the informative PCs can be used directly for homology assignments via *Constella* it is worth first considering the variation of each via their loadings prior to downstream analyses.

When the factor loadings across all genotypes are visualized as a single graph it is apparent that there is underlying structured variation within the PC space, which resembles a torus with a cylinder running through its core (Fig. 8A). The torus which is predominantly represented by PC1 and PC2 is comprised of positional information based on the X-Y coordinates for p-lms (Fig. 8B) or their relative distances from bounding box corners (Fig. 8C). It is unsurprising that there should be great overlap between the assigned tip of the acute region (plm-x, plm-y), and the starting and ending points used with the tip to measure orientation (SS-x, SS-y, TS-x, TS-y), since these define virtually the same coordinates. The cylinder varies predominantly in the third PC axis space and describes distance of each p-lm from the centroid of each plant (Fig. 8C pink dots) and the concavity-convexity scores for individual p-lms (Fig. 8C grey dots). The orientation and centroid orientation vary mostly in the third PC axis but appear as somewhat diffuse clouds of points (Fig. 8D).

**Figure 8.**
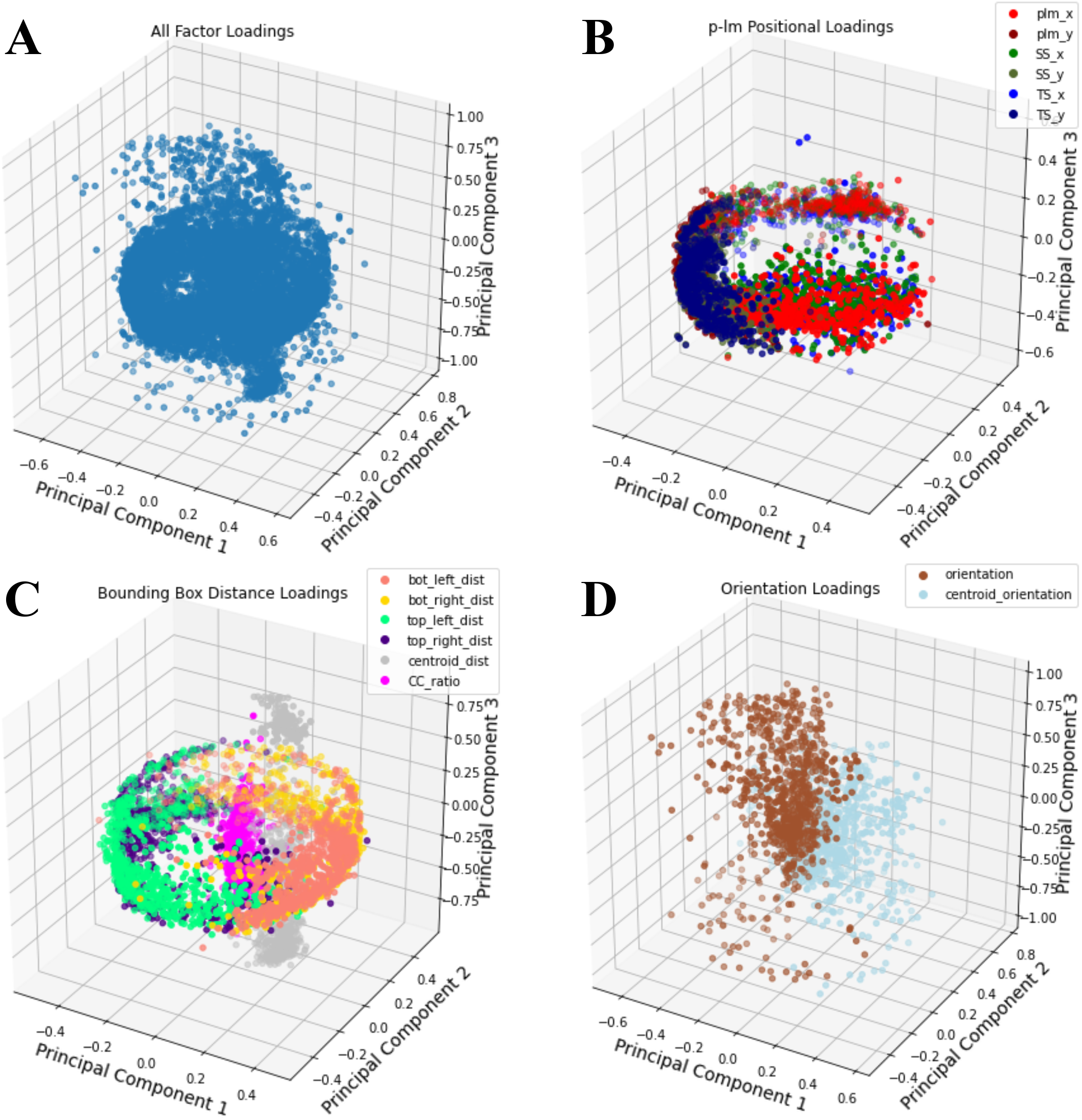
Factor loadings for various p-lm attributes on the informative PC axes across genotypes. (A) Loadings for all factor for all genotypes. This shows a general conserved symmetry of a torus in the PC1 and PC2 dimensions and a cylinder running through its center that preferentially loads onto the PC3 axis. (B) The colorized p-lm (red), starting site (green), and termination site (blue) x and y values which show that the coordinate positions preferentially load into the torus. (C) The colorized distances from each bounding box corner, in addition to the distance from the centroid (gray), and the CC ratio (pink). The bounding box distances again appear to preferentially load into the torus whereas the centroid distance and CC ratio appear to compose the cylinder which loads preferentially onto PC3. (D) The individual p-lm orientation (brown) and orientation with respect to the centroid (silver) loadings appear to have no overlap with one another and are distributed diffusely in the 3D PCA space.

### Accuracy of *constella* in nearest neighbor homology grouping assignments

To assess the accuracy of *constella*, a subordinate function, *constellaQC*, was used, which calculated homology grouping accuracy based both on the rate of valid homology calls made as well as rates of the types of spurious calls, splitting errors (creating more than one group for the same structure) and clumping errors (clumping more than one structure into a single group). *ConstellaQC* was first used to assess the accuracy of the pipeline with raw *acute* outputs (i.e. prior to the use of *space* or the dimensionality reduction of *starscape*). While approximately 50% of neighboring points in consecutive images were correctly called, approximately 25% each of splitting (creating more than one group for the same structure) and clumping (clumping more than one structure into a single group) errors was found (Table 1). When *starscape* was used alone, valid call rates increased to 80-90%, showing that the refinement afforded by the principal components analysis of distances was of great value in linking homologous p-lms together between consecutive images (Table 1). We then used the *space* function to modify the PCA outputs and assess the usefulness of adding remeasured distances from the bounding box corners and from the centroid for each image, as well as the addition of data on orientation of each p-lm and whether the p-lm described a convex (leaf tip) or concave (leaf blade insertion point) structure.

**Table 1.** Homology accuracy and error rates estimated by *constellaQC* when run under various scenarios in which either *space* and *starscape*, *space*, or certain metadata dimensions of *space* are omitted prior to running *constella*.

| Valid Call Rates (%) | A10 | B100 | RIL39 | RIL110 | RIL159 | Mean |
| --- | --- | --- | --- | --- | --- | --- |
| Grouping without *space* or *starscape* | 43.65 | 63.99 | 46.54 | 50.5 | 35.01 | 47.94 |
| Grouping without *space* | **82.79** | **93.59** | 86.34 | 87.33 | 82 | 86.41 |
| Grouping with *space* (all attributes) | 82.07 | 93.42 | 86.77 | 87.8 | 82.35 | 86.482 |
| Excluding bounding box distances | 82.36 | 93.18 | 86.07 | 87.48 | 82.27 | 86.272 |
| Excluding bounding box and centroid distances | 81.26 | 93.35 | 85.71 | 87.98 | 82.33 | 86.126 |
| Excluding all p-lm orientations | 82.04 | 93.52 | 86.45 | **88.08** | 82.51 | 86.52 |
| Excluding centroid distance and p-lm orientations | 81.67 | **93.59** | **86.97** | 87.93 | **82.59** | **86.55** |
| **Maximum Percentage** | **82.79** | **93.59** | **86.97** | **88.08** | **82.59** | **86.55** |
| Splitting Call Rates (%) | A10 | B100 | RIL39 | RIL110 | RIL159 | Mean |
| Grouping without *space* or *starscape* | 28.77 | 18.31 | 27.03 | 25.3 | 33.5 | 26.58 |
| Grouping without *space* | **7.83** | **2.89** | 5.87 | 5.74 | 8.55 | 6.176 |
| Grouping with *space* | 8.26 | 3 | 5.67 | 5.51 | 8.33 | 6.154 |
| Excluding bounding box distances | 8.09 | 3.13 | 6.04 | 5.64 | 8.37 | 6.254 |
| Excluding bounding box and centroid distances | 8.63 | 3.03 | 6.2 | 5.41 | 8.34 | 6.322 |
| Excluding all p-lm orientations | 8.21 | 2.93 | 5.8 | **5.37** | 8.25 | 6.112 |
| Excluding centroid distance and p-lm orientations | 8.42 | **2.89** | **5.57** | 5.46 | **8.22** | **6.112** |
| **Minimum Percentage** | **7.83** | **2.89** | **5.57** | **5.37** | **8.22** | **6.112** |
| Clumping Call Rates (%) | A10 | B100 | RIL39 | RIL110 | RIL159 | Mean |
| Grouping without *space* or *starscape* | 27.58 | 17.7 | 26.43 | 24.2 | 31.5 | 25.48 |
| Grouping without *space* | **9.38** | **3.52** | 7.79 | 6.93 | 9.45 | 7.414 |
| Grouping with *space* | 9.67 | 3.59 | 7.56 | 6.7 | 9.33 | 7.37 |
| Excluding bounding box distances | 9.55 | 3.69 | 7.89 | 6.88 | 9.36 | 7.474 |
| Excluding bounding box and centroid distances | 10.12 | 3.62 | 8.08 | 6.6 | 9.33 | 7.55 |
| Excluding all p-lm orientations | 9.75 | 3.55 | 7.76 | **6.56** | 9.24 | 7.372 |
| Excluding centroid distance and p-lm orientations | 9.91 | **3.52** | **7.46** | 6.6 | **9.19** | **7.336** |
| **Minimum Percentage** | **9.38** | **3.52** | **7.46** | **6.56** | **9.19** | **7.336** |

The usefulness of the various additional data-types provided by *space* was assessed, and to our surprise we found that, when all data types were used, the accuracy as measured by *constellaQC* was never better than without the use of *space*. When individual data types were individually assessed, the addition of only the distances from the bounding box corners equaled or marginally increased valid call accuracy in three of the five genotypes. the addition of centroid distances improved the call rate for one genotype, whilst the remaining genotype had the highest valid call rate without the use of the *space* function. In all cases, differences were less than 1%, making the need for the *space* function debatable.

## Discussion

### *Acute* pseudo-landmarking

The quantification of morphology has long been a practice to explore biological systems and characterize the mechanisms underlying them. Although numerous manual and automated tools have been developed to address these phenotyping efforts *acute* has the advantage of being able to readily generate pseudo landmarks based on geometry alone, to identify structures of interest such as leaf tips or ligules. *Acute* appears to be precise, with a high degree of correlation between de novo p-lm measures and manual measurements. Accuracy is also high, although occasionally slightly biased by morphological differences. However, it appears from our results that such biases are systematic and relate to differences in which point on the plant is actually being measured. As such, they are easily corrected if need be. Although culm height is largely used in this context as a test case, we observed differences in internode expansion rates between genotypes and between internode positions within genotypes that offer intriguing insights into growth patterns of the whole plant. These will be the subject of future investigations.

### The *homology* workflow

The homology groupings generated by the *homology* workflow are based on the idea of object permanence where the absolute positions of points in space between consecutive images matters little compared to their location in space with respect to one another. Simply put, this method assumes that the majority of structures maintain more or less the same location in space and are unlikely to migrate, allowing most components of these constellations of p-lms to effectively remain stationary through time while younger homology groups are still actively migrating to their stationary positions. The iterative developmental strategy lends itself well to this method of homology assignment via conserved constellations, given that the majority of structures in a plant’s body plan will be terminally differentiated and static over time. The use of principal component analysis using *starscape* is crucial for accurately linking p-lms that represent the same plant structure through time, but we did not see much gain in the valid call rate by incorporating additional data dimensions through the *space* function. It is possible that different data types, such as hyperspectral data (e.g. Miao et al. 2020) could be incorporated into the *space* function to improve the rate of valid calls. This general workflow is a special case of clustering following principal component analysis, although for our application hierarchical clustering is used in place of the more common K-means generally used in these workflows (Ding and He, 2004).

By clustering on component axes, information for successful homology grouping may be maximized either through removing correlated variation between factors or scale differences between dimensions. Moreover, although the homology assignments described here were of p-lms derived from *acute*, we suggest that both *starscape* and *constella* should be useful for other datasets that provide morphological information. Those datasets with more dimensionality, such as 3D morphology datasets (e.g. Zhang et al. 2017) could in fact be expected to have higher valid call rates because of the increased ability to characterize structures through time. The need to assess the factor loadings generated by *starscape* will likely serve as a best practice when utilizing this workflow on a new type of morphological data. Ultimately, the *homology* workflow is designed to be a robust tool for assigning identity to nearest neighbors through time and should be able to extend to broader applications than illustrated in this study.

## Conclusions

This workflow represents an effective means for studying plant growth through time, by providing a solution to the problem of delineating equivalent positions between consecutive images in time-series. We have shown that the pseudo-landmarks generated by *acute* can effectively capture plant morphological variation and provide a means to analyze differences in growth between genotypes. As demonstrated by comparisons of manual measures such as culm height the pseudo-landmarks precisely and accurately measure traits of interest, with minimal, and easily correctable, bias. The *homology* workflow, particularly *starscape* and *constella*, serves as a relatively simple and effective manner of clustering segmented plant data across timeseries images. Moreover, the demonstration of different strategies of utilizing and exploring the outputs of these functions reveal that attempts to mine additional metadata from existing variation such as the implementation of *space* in our analysis can offer improvements but often at too modest of an amount to warrant their usage. *Constella* through its usage of inferring shared identity based on a duet/quartet grouping scheme to find nearest neighbors has proven to be robust even within the somewhat complex morphologies generated in this study. In addition, *constellaQC* provides a means of assessing different types of error such as clumping, which represents confounding two distinct structures, and splitting, which represents mistaking the same structure over time for discrete objects. The two pipelines represent fast and accurate methods to automate and easily curate time-series data sets of plant growth and development.

## Acknowledgements

We would like to thank Samantha M. Cady for her feedback during the development and implementation of the *starscape* and *constella* homology grouping procedures.

