## Appendix 1 for "De novo homology assessment from landmark data: A workflow to identify and track segmented structures in plant time series images"

### Exercise 1: Overview of the Acute pseudo-landmarking

This is an introduction to the pseudo-landmarking method based around the acute function found within PlantCV. This tool is designed for morphometric analysis which due to it's relative simplicity can easily scale between different datasets in order to capture informative shape data in the form of de novo landmarks. This notebook serves as demonstration of the image data curration required by acute and also documents the initial outputs of acute which can be used either to optimzie this workflow. Later exercises will build off of what is covered within this document in order to show the potential of this method to end users. To begin, let's start by loading the modules we'll need and then take stock of the acute function and how it operates by running help to see what inputs it requires...

In [1]:

```python
import cv2
import sys
import pandas as pd
import numpy as np
import matplotlib.pyplot as plt
from plantcv.plantcv.homology.acute import acute

debug = True

help(acute)
```

```
Help on function acute in module plantcv.plantcv.homology.acute:

acute(obj, mask, win, threshold, debug)
    acute: identify landmark positions within a contour for morphometri
c analysis

    Inputs:
    obj          = An opencv contour array of interest to be scanned for
landmarks
    mask         = binary mask used to generate contour array (necessary
for ptvals)
    win          = maximum cumulative pixel distance window for calculat
ing angle
                   score; 1 cm in pixels often works well
    thresh       = angle score threshold to be applied for mapping out l
andmark
                   coordinate clusters within each contour
    debug        = Debugging mode enabled/disabled for use in troublesho
oting

    Outputs:
    homolog_pts = pseudo-landmarks selected from each landmark cluster
    start_pts   = pseudo-landmark island starting position; useful in p
arsing homolog_pts in downstream analyses
    stop_pts    = pseudo-landmark island end position ; useful in parsi
ng homolog_pts in downstream analyses
    ptvals      = average values of pixel intensity from the mask used
to generate cont;
                   useful in parsing homolog_pts in downstream analyses
    chain       = raw angle scores for entire contour, used to visualiz
e landmark
                   clusters
    verbose_out = supplemental file which stores coordinates, distance
from
                   landmark cluster edges, and angle score for entire co
ntour.  Used
                   in troubleshooting.

    :param obj: ndarray
    :param mask: ndarray
    :param win: int
    :param thresh: int
    :return homolog_pts:
```

### Required image input variables

We'll need 4 variables for each image we run. Two are derived from the image itself, a contour array representing the outline of a plants image mask (obj) and the image mask itself which is used for output purposes (mask).

Let's first start by creating these first two objects. To begin, let's load our first image from a time series sequence.

In [2]:
```
day=10

path='/Path/To/Images/plm_tutorial/'
name='B100_rep1_d'+str(day)

img = cv2.imread(path+name+'.jpg')

#Plot results
fig1=plt.figure(figsize=(6, 8))
fig1=plt.imshow(img)
fig1=plt.xscale('linear')
fig1=plt.axis('off')
fig1=plt.title('B100 day '+str(day))
plt.show(fig1)
```

B100 day 10

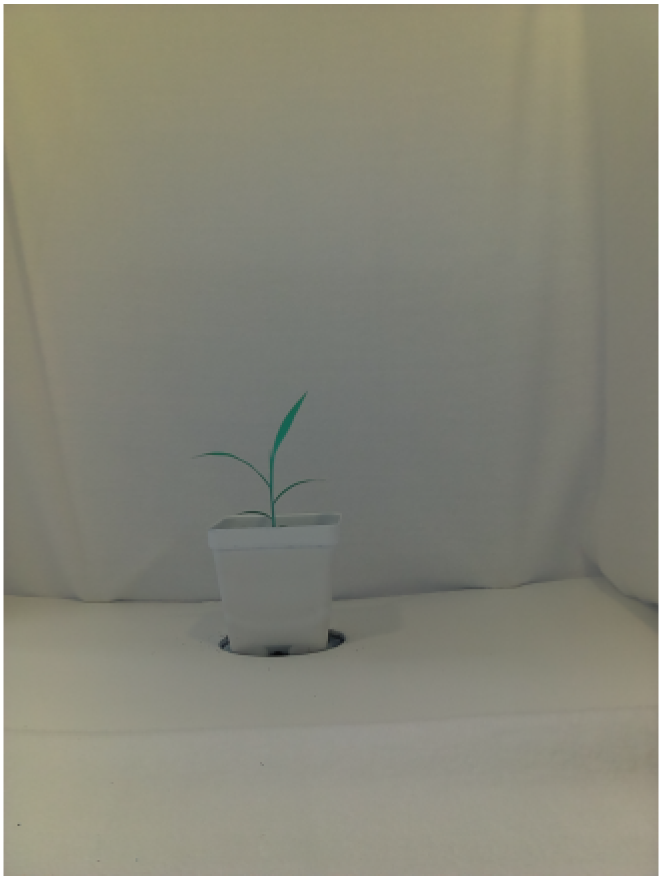

### Reviewing our loaded image

From what we can see, the plant is on a mostly homogeneous white background. It should be relatively easy to use the color channel differences to threshold the pixels representing our plant from the rest of the image to create our mask. To begin let's take a look at the color channels to see which will be the most useful.

In [3]:
```
lab_img=cv2.cvtColor(img, cv2.COLOR_BGR2LAB)
hsv_img=cv2.cvtColor(img, cv2.COLOR_BGR2HSV)

img_hsv_lab_colorspaces = cv2.hconcat((lab_img, hsv_img))

#Plot results
fig1=plt.figure(figsize=(12, 8))
fig1=plt.imshow(img_hsv_lab_colorspaces)
fig1=plt.xscale('linear')
fig1=plt.axis('off')
fig1=plt.title('Images of \'Lab\' and \'HSV\' color spaces respectively'
)
plt.show(fig1)
```

Images of 'Lab' and 'HSV' color spaces respectively

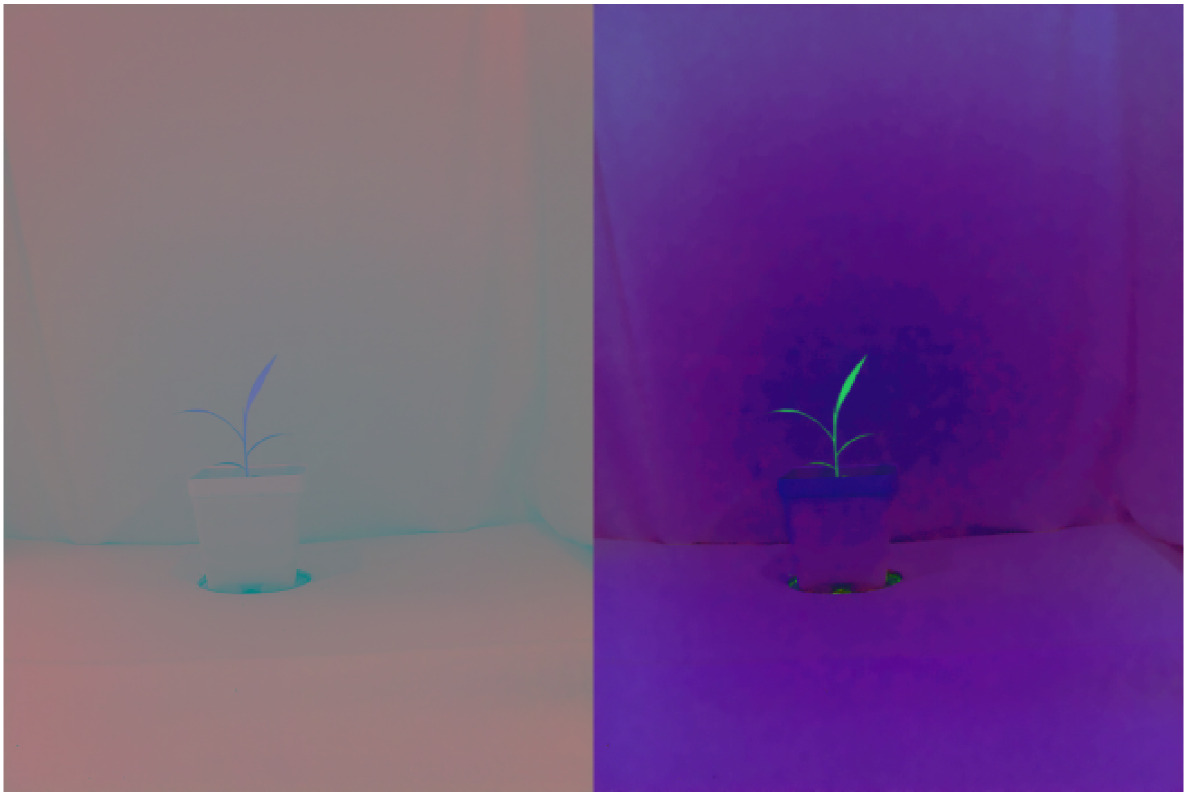

### Selecting color channels for thresholding

In comparing the 'Lab' and 'HSV' color spaces it appears there's a bit more contrast to work within the HSV space, but that soil near the stage for the pot could be a problem. With that in mind the 'Lab' color space is our best bet. Let's take a look at which individual channels are the most informative...

In [4]:
```
img_l, img_a, img_b = cv2.split(lab_img)

img_lab_channels = cv2.hconcat((img_l, img_a, img_b))

#Plot results
fig1=plt.figure(figsize=(21, 10))
fig1=plt.imshow(img_lab_channels, 'gray')
fig1=plt.xscale('linear')
fig1=plt.axis('off')
fig1=plt.title('Grayscale images of \'L\', \'a\', and \'b\' channels res
pectively')
plt.show(fig1)
```

Grayscale images of 'L', 'a', and 'b' channels respectively

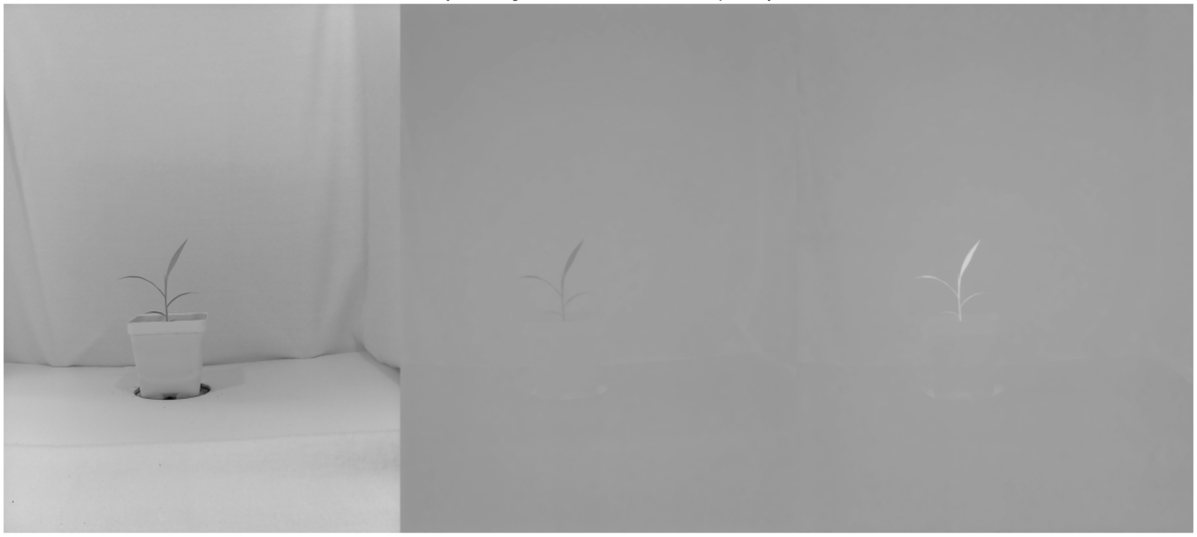

### Binary thresholding

The 'L' channel unfortunately doesn't appear to help very much in denoting the plant pixels from the background. However, there's good signal in the 'a' channel (darker pixels) and the 'b' channel (brighter pixels) so using a conjunction of these two grayscale images should give us a reasonable mask to work with!

In [5]:
```
#These threshold bounds will provide the best signal but feel free to ex
periment!
a_bound = np.array([123, 255])
b_bound = np.array([133, 255])

#Note that we're inverting the binary threshold of color channel 'a' so
 that the areas
#with the darkest pixels will be flagged as a white mask.  This will be
 important when
#compared against the mask generated from color channel 'b'.
mask_a = cv2.threshold(img_a, a_bound[0], a_bound[1], cv2.THRESH_BINARY_
INV)
a_thresh = cv2.cvtColor(mask_a[1], cv2.COLOR_GRAY2RGB)

mask_b = cv2.threshold(img_b, b_bound[0], b_bound[1], cv2.THRESH_BINARY)
b_thresh = cv2.cvtColor(mask_b[1], cv2.COLOR_GRAY2RGB)

img_ab_thresholds = cv2.hconcat((a_thresh, b_thresh))

#Plot results
fig1=plt.figure(figsize=(12, 8))
fig1=plt.imshow(img_ab_thresholds, 'gray')
fig1=plt.xscale('linear')
fig1=plt.axis('off')
fig1=plt.title('Binary images of \'a\' and \'b\' thresholds respectivel
y')
plt.show(fig1)
```

Binary images of 'a' and 'b' thresholds respectively

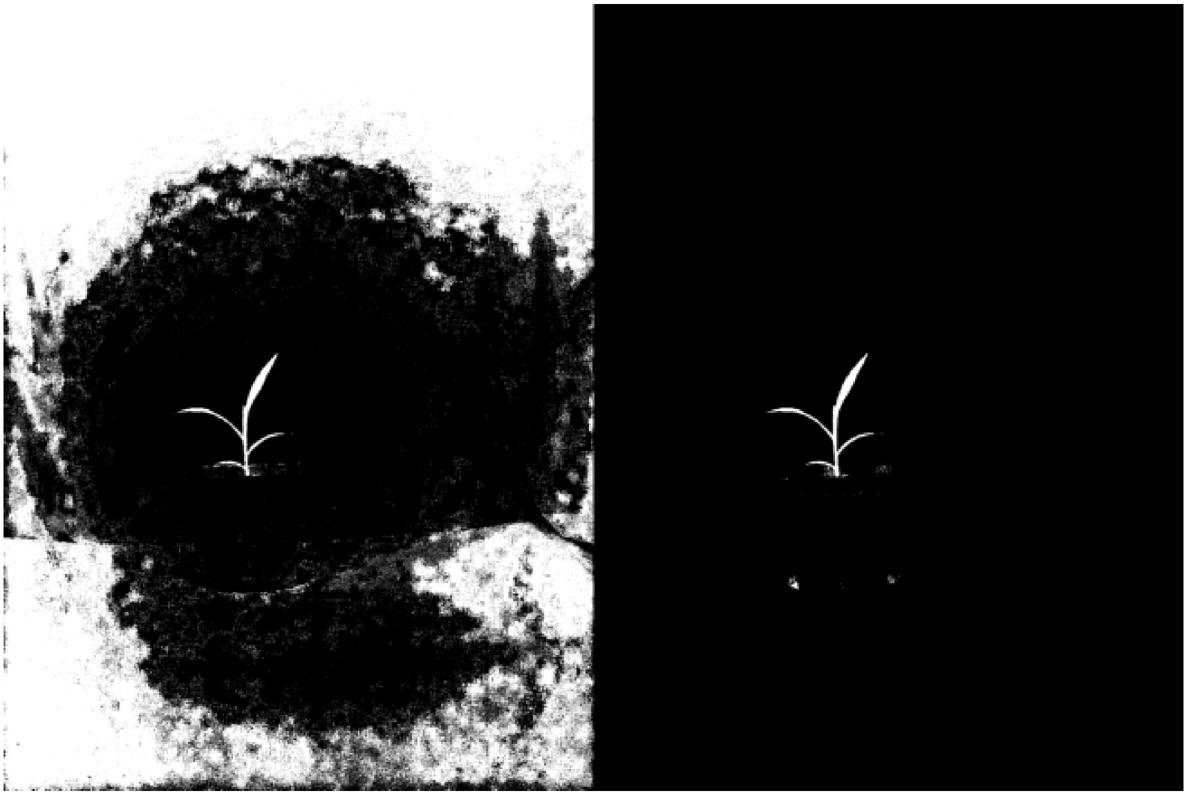

### Merging binary thresholds into our mask

In both cases we have a few stray pixels (more so in mask a), but we're certainly on the right track! Now lets go ahead and identify which pixels these thresholds can agree on keeping.

In [6]:
```
mask=cv2.bitwise_and(mask_a[1], mask_b[1])

#Plot results
fig1=plt.figure(figsize=(6, 8))
fig1=plt.imshow(mask, 'gray')
fig1=plt.xscale('linear')
fig1=plt.axis('off')
fig1=plt.title('Merged mask')
plt.show(fig1)
```

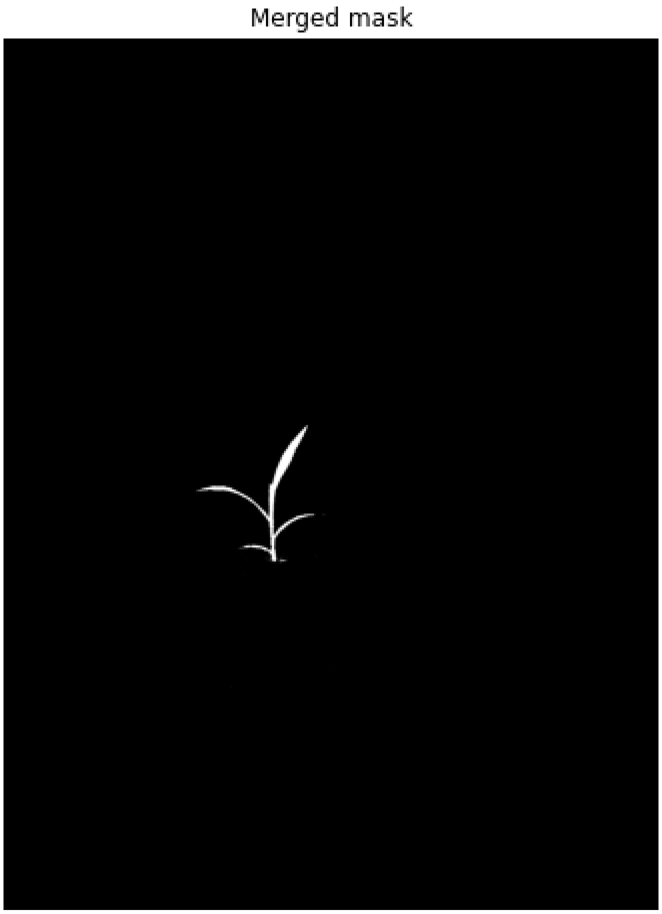

### Extracting OpenCV image contour arrays

Now with our merged mask defining the shape our our plant we can extract our contour to use for pseudo-landmark identification. This will be done through the use of the findContours function latent to openCV where we will use a simple approximation (to make this step less computationally intensive) and a tree hierarchy will be extracted as well (important for images in which internal volumes resulting from crossovers between structures occurs).

Although not necessary yet in this demonstration the steps below demonstration how the plants outer contour is defined based on which bears the largest volume. Contours contained within this 'parent' contour are then stored as well for downstream analysis within a contour list. This will effectively exclude other components of the mask unrelated to our plant.

In [7]:

```
cont, hierarchy = cv2.findContours(mask, cv2.RETR_TREE, cv2.CHAIN_APPROX
_SIMPLE)

mask_contour = cv2.cvtColor(mask, cv2.COLOR_GRAY2RGB)

#Find largest contour of subject (outer boundary of subject)
cont_list = []
hull = [0, 0]
for c in range(len(cont)):
    a = cv2.contourArea(cont[c])
    if a > hull[0]:
        hull = [a, c]

cont_list.append(hull[1])
#Capture children of parent contour
for e in range(len(hierarchy[0])):
    if (hierarchy[0][e][3] == hull[1]) & (len(cont[e]) > 10):
        cont_list.append(e)

#Draw the individual contour outlines onto the duplicate mask in a purpl
e hue
for c in cont_list:
    cv2.drawContours(mask_contour, cont[c], -1, (180, 0, 180), 8)

#Plot results
fig1=plt.figure(figsize=(6, 8))
fig1=plt.imshow(mask_contour)
fig1=plt.xscale('linear')
fig1=plt.axis('off')
fig1=plt.title('Plant contour (purple)')
plt.show(fig1)
```

Plant contour (purple)

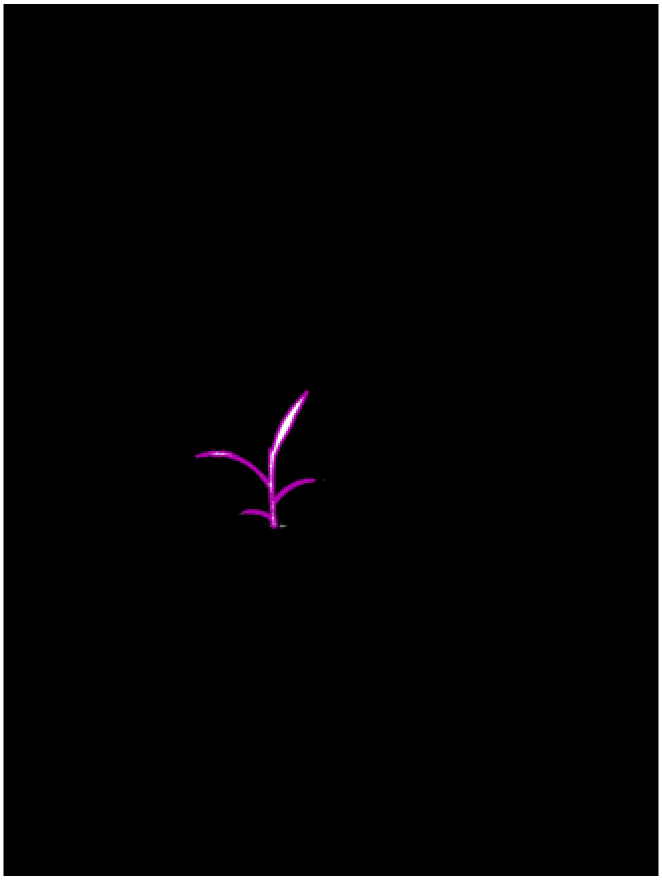

### Acute pseudo-landmark identification

Now that we have our contours we can come back to the previous two input parameters of acute which we have thus far ignored but are key to it's functionality. Acute operates using a modified form of chain-coding akin to a navigators compass taking steps along a contour and within a local window two bounding points on either side of this window are defined from which an angle score can be calculated for the vertex of the 3 points. The size of this local window is defined as a pixel distance using the 'win' variable. Following the calculation of this angle score it is then weighed against a threshold that is stored in our last variable 'thresh' allowing for features of interest to be defined de novo. Given that acute regions are often areas of interest for morphometric analysis setting this threshold to maximize the 'acuteness' of the contour serves to provide a relatively simple way to identify pseudo-landmarks.

When specifying 'win' it is often best to select a value which is at least half the distance of the smallest feature in the plant that is deemed relevant. In the case of Setaria which we are using in this demonstration the first leaf is usually 2 cm long so selecting a window size <=1 cm is optimal to prevent conflict between adjacent landmarks along the contour.

When specifying 'thresh' the best practice is to leave this value at 90 given in order to identify acute regions. However, to provide downstream flexibility this parameter has the capacity to use other user defined values in case more stringent or lax thresholds are required.

In [8]:
```
win=25
thresh=90
```

### Running acute in debugging mode

As is standard with other PlantCV packages acute is built with debugging features that produces verbose outputs for the sake of troubleshooting. Given this is our first attempt at running this function lets go ahead and run it with debugging enabled to see what these outputs are...

*Note: while iterating through the contour list isn't necessary for a single outline as we have here this step is invaluable in later stages where volumes internal to our plant outline are present.

In [9]:
```
landmark_output=[]

for l in cont_list:
    if cv2.arcLength(cont[l],True) > 2*win:
        print('Contour volume: '+str(cv2.arcLength(cont[l],True)))

        cv2.drawContours(mask, cont[l], -1, (128,0,0), 3)
        homolog_pts, homolog_start, homolog_stop, homolog_cc, chain, ver
bose = acute(cont[l], mask, win, thresh, debug)
        homolog_hier = l*len(homolog_pts)
        cv2.drawContours(mask, homolog_pts, -1, (0,0,255), 3)
        print('     ' + 'landmark number: ' + str(len(homolog_pts)))

        for h in range(0,len(homolog_pts)):
            landmark_output.append([name, homolog_pts[h][0][0], homolog_
pts[h][0][1], homolog_start[h][0][0], homolog_start[h][0][1], homolog_st
op[h][0][0], homolog_stop[h][0][1], homolog_cc[h],])
```

```
Contour volume: 2134.4002673625946
Fusing contour edges
route C
Landmark site:  1125  , Start site:  1113  , Term. site:  19
Landmark point indices:  [1125]
Starting site indices:  [1113]
Termination site indices:  [19]
route C
Landmark site:  203  , Start site:  188  , Term. site:  221
Landmark point indices:  [1125, 203]
Starting site indices:  [1113, 188]
Termination site indices:  [19, 221]
route C
Landmark site:  353  , Start site:  337  , Term. site:  363
Landmark point indices:  [1125, 203, 353]
Starting site indices:  [1113, 188, 337]
Termination site indices:  [19, 221, 363]
route C
Landmark site:  538  , Start site:  531  , Term. site:  547
Landmark point indices:  [1125, 203, 353, 538]
Starting site indices:  [1113, 188, 337, 531]
Termination site indices:  [19, 221, 363, 547]
route C
Landmark site:  594  , Start site:  571  , Term. site:  602
Landmark point indices:  [1125, 203, 353, 538, 594]
Starting site indices:  [1113, 188, 337, 531, 571]
Termination site indices:  [19, 221, 363, 547, 602]
route C
Landmark site:  672  , Start site:  655  , Term. site:  689
Landmark point indices:  [1125, 203, 353, 538, 594, 672]
Starting site indices:  [1113, 188, 337, 531, 571, 655]
Termination site indices:  [19, 221, 363, 547, 602, 689]
route C
Landmark site:  809  , Start site:  795  , Term. site:  824
Landmark point indices:  [1125, 203, 353, 538, 594, 672, 809]
Starting site indices:  [1113, 188, 337, 531, 571, 655, 795]
Termination site indices:  [19, 221, 363, 547, 602, 689, 824]
route C
Landmark site:  895  , Start site:  877  , Term. site:  908
Landmark point indices:  [1125, 203, 353, 538, 594, 672, 809, 895]
Starting site indices:  [1113, 188, 337, 531, 571, 655, 795, 877]
Termination site indices:  [19, 221, 363, 547, 602, 689, 824, 908]
    landmark number: 8
```

### Acute angle score chain-code output

We did quite a bit of work just above so let's go ahead and try to break it down item by item. To start we successfully ran acute which used it's chain coding based scoring method to identify landmarks (8 in this example). However we started with several hundred vertices comprising our outline (the purple dots we saw earlier) so how did we reduce those down to 8 pseudo-landmarks!?

Acute actually undergoes an extra step beyond simply calculating angle scores in that it attempts to identify 'islands' of acute points and upon finding these regions it approximates the best mid point of each location within the contour. Let's have a look at our contour described by it's acute angle scores...

In [10]:
```
chain_pos=range(0, len(chain))

fig, (fig1, fig2) = plt.subplots(1, 2, figsize=(12, 6))

#Plot results
fig1.plot(chain_pos, chain, color='black')
fig1.axhline(y=thresh, color='r', linestyle='-')
fig1.set_title('Angle scores by position')

fig2.hist(chain, color='black')
fig2.axvline(x=thresh, color='r', linestyle='-')
fig2.set_title('Angle score histogram')

plt.show(fig)
```

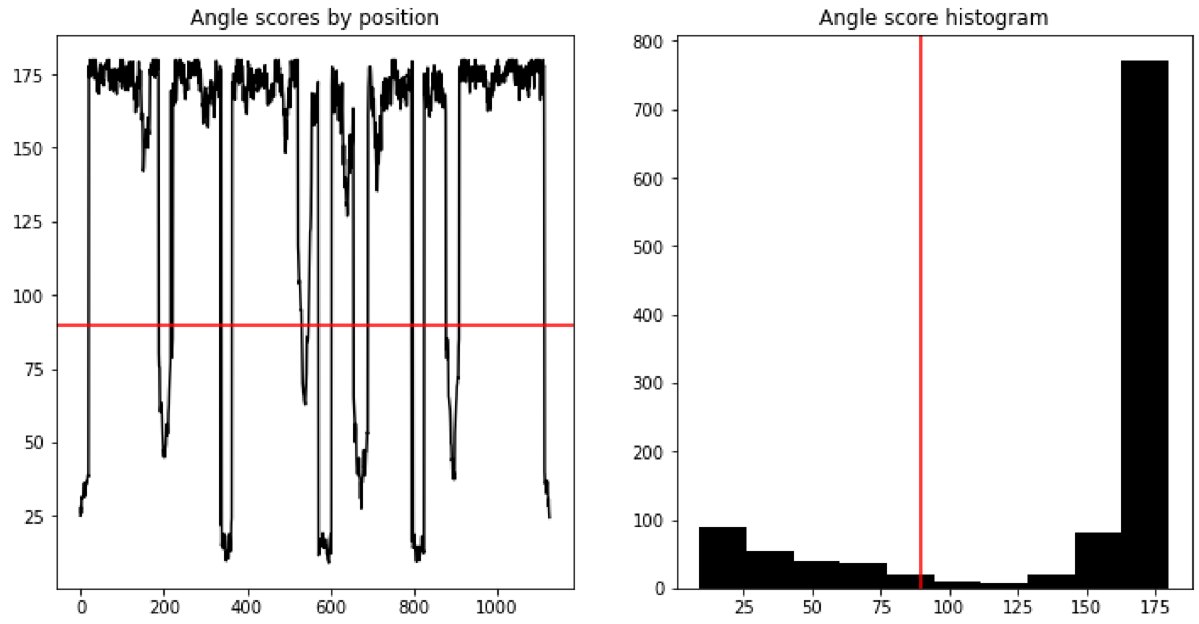

If we focus on our first graph on the left the black line representing the individual acute angle scores of each vertex along our contour outline we can see there definitely seems to be a waveform that quickly decays to zero as we hit each acute island. It's also probably apparent we're actually splitting on of these islands in half at either end of our chain in linearizing this output. The 'Fusing contour edges' step acute does automatically (appeared in the output panel) is performed to remedy mistaking these two segments as different regions.

When we compare these regions to our threshold (the red line) it becomes clear these 'waves' correspond to our landmarks. If we felt a particular need to optimize this threshold since we should optimally have a bimodal output a histogram of the angle scores can be generated as shown on the right to train this threshold in order to better optimize signal.

Now that we have a general idea of how acute is determining it's primary outputs let's see how they compare to our original image...

In [11]:
```
img_plms = img.copy()

for c in cont_list:
    cv2.drawContours(img_plms, homolog_pts, -1, (255, 255, 255), 14)

#Plot results
plm_fig=plt.figure(figsize=(7, 10))
plm_fig=plt.imshow(img_plms)
plm_fig=plt.xscale('linear')
plm_fig=plt.axis('off')
plm_fig=plt.title('B100 day '+str(day))
plt.show(plm_fig)
```

B100 day 10

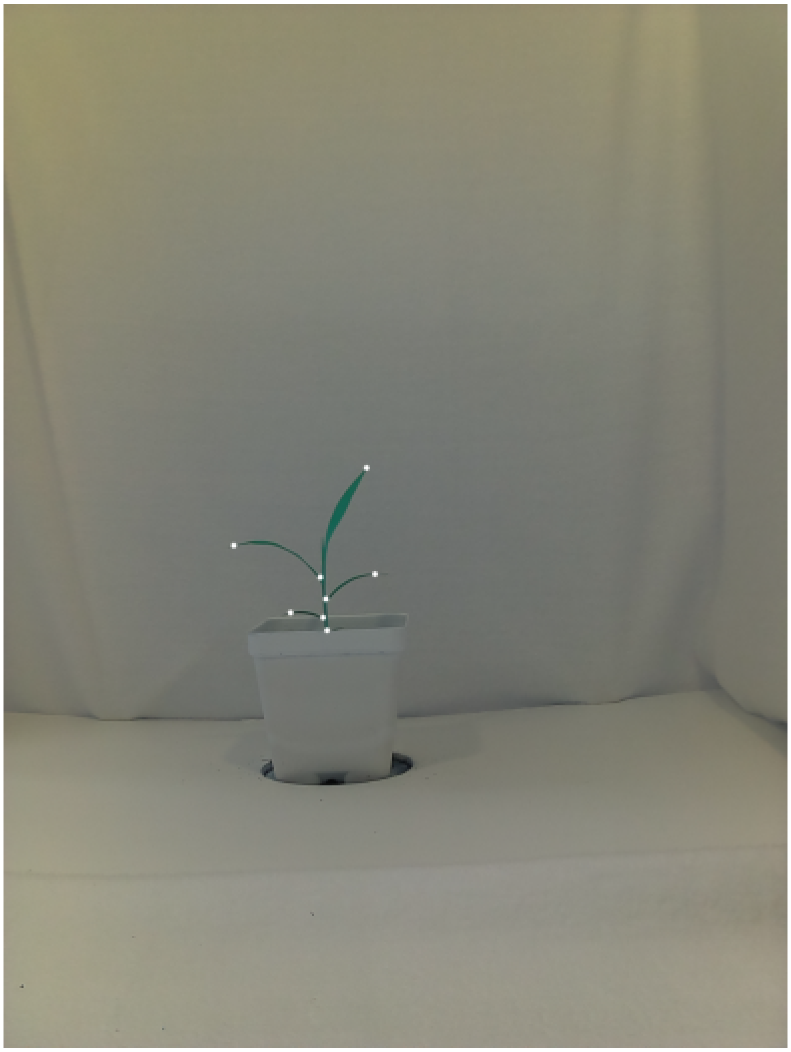

In viewing our plms plotted in white while looking back on our original image we used for this exercise we can see that not only were we able to clearly define regions of interest using a threshold on our angle score waveforms but these do in fact appear to correlate with regions of interest for morphometric analysis such as the tips of leaves as well as the ligules where the base of each leaf attaches to the culm (the grass equivalent to a stem). This is the basis by which acute functions and serves as the basic mode of operation for this workflow. In the next exercise 2 we will expand on this knowledge by learning to interact with and store time series data which in exercise 3 can subsequently be fused together into homology groups.

```
In [ ]:
```

### Exercise 2: Applying acute on batch datasets

In the previous exercise we learned how to use some basic tools latent to PlantCV to explore color spaces and extract plant shapes from an image then use this shape data for generating pseudo-landmarks (plms) with acute while thinking a bit about how acute is making its calls for de novo landmarks. While this is useful and necessary we also need to think about how best to scale up our code in order to be used on batch datasets which can consist of multiple images either of the same genotype or of comparable stages of different genotypes. For the sake of this demonstration we'll design a loop to run acute on a time series from the dataset we began to explore in the previous exercise. With that all being said lets get started by taking a look at the files we have on hand...

In [1]:
```
import cv2
import os
import sys
import pandas as pd
import numpy as np
import matplotlib.pyplot as plt

from plantcv.plantcv.homology.acute import acute

win=25
thresh=90
debug = True

path='/Path/To/Images/plm_tutorial/'

os.listdir(path)
```

Out[1]:
```
['.DS_Store',
 'B100_rep1_d12.jpg',
 'B100_rep1_d13.jpg',
 'B100_rep1_d11.jpg',
 'B100_rep1_d10.jpg']
```

So already we can see that there seems to be a general theme with how these images are named following a 'Genotype' + 'Timepoint' + '.jpg' format. Although this is just one form of serial naming strategy it is always advised with projects that will consist of large scale datasets to perform some initial step of data carpentry and decide a consistent manner of naming early on. The reason this is important we'll see below when we design a for loop. Before we run the loop itself let's go ahead and specify the variables we'll need for this serial naming scheme and discuss what each one represents...

In [2]:
```
days=range(10,14)
name_prefix='B100_rep1_d'
```

So we have 3 different variables we'll need we've created above days which is a list of integers that ranges from 10-13 in a pythonic fashion, a name prefix we can attach to each days integer to complete our serial names, and we'll also need a path to our files in the directory space which we have already specified earlier. Before we go ahead and scale up our code from the previous exercise let's first build a dummy loop which can perform a simple task on these files to be sure that we can read them in properly...

In [3]:
```
for day in days:
    img = cv2.imread(path+name_prefix+str(day)+'.jpg')
    fig1=plt.figure(figsize=(6, 8))
    fig1=plt.imshow(img)
    fig1=plt.xscale('linear')
    fig1=plt.axis('off')
    fig1=plt.title(name_prefix+str(day))
    plt.show(fig1)
```

B100_rep1_d10

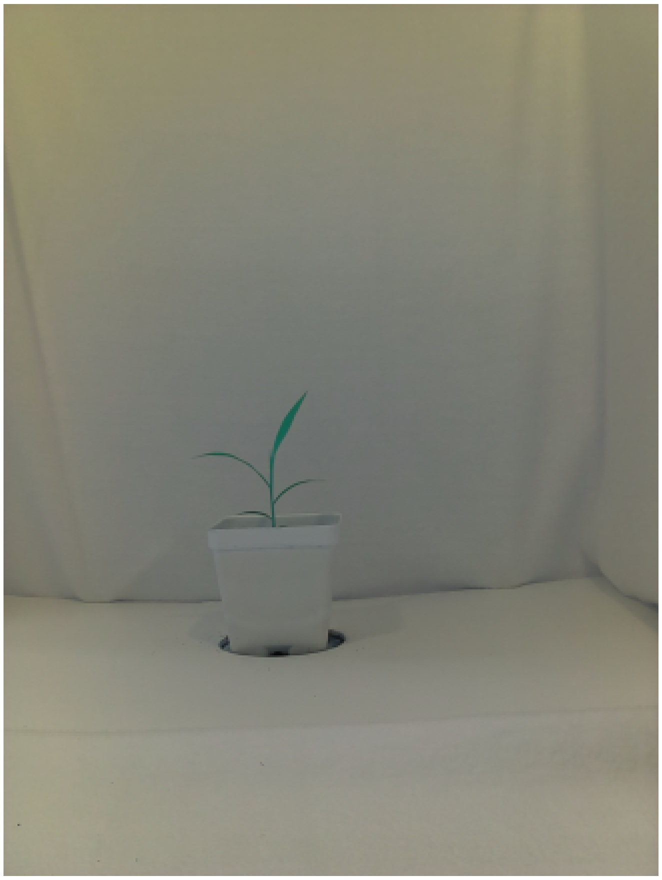

B100_rep1_d11

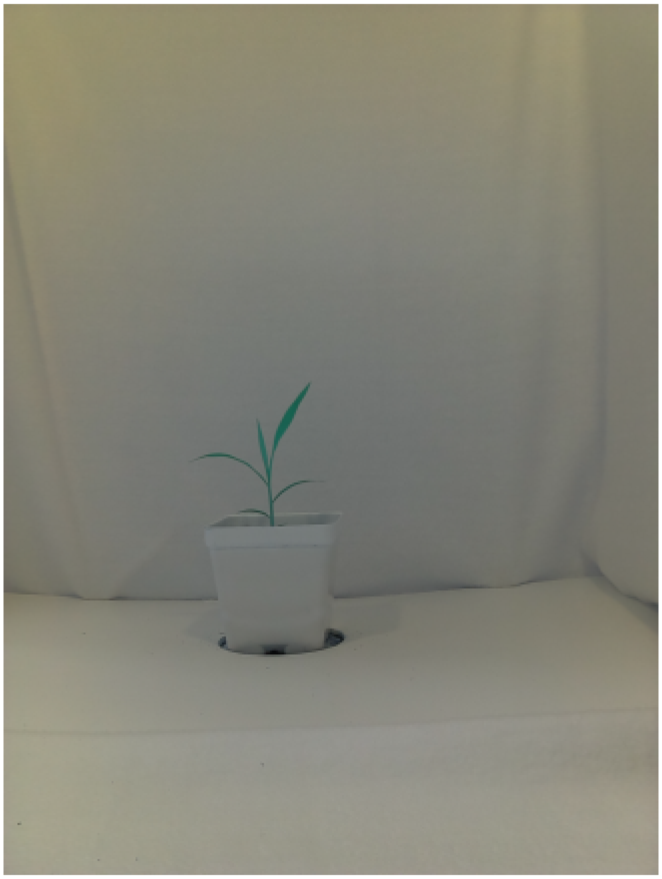

B100_rep1_d12

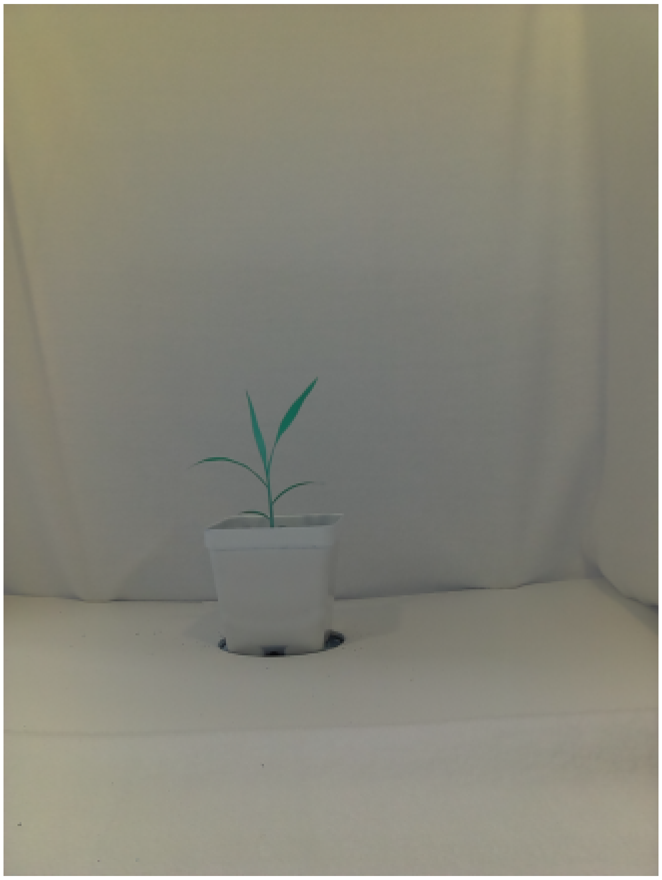

B100_rep1_d13

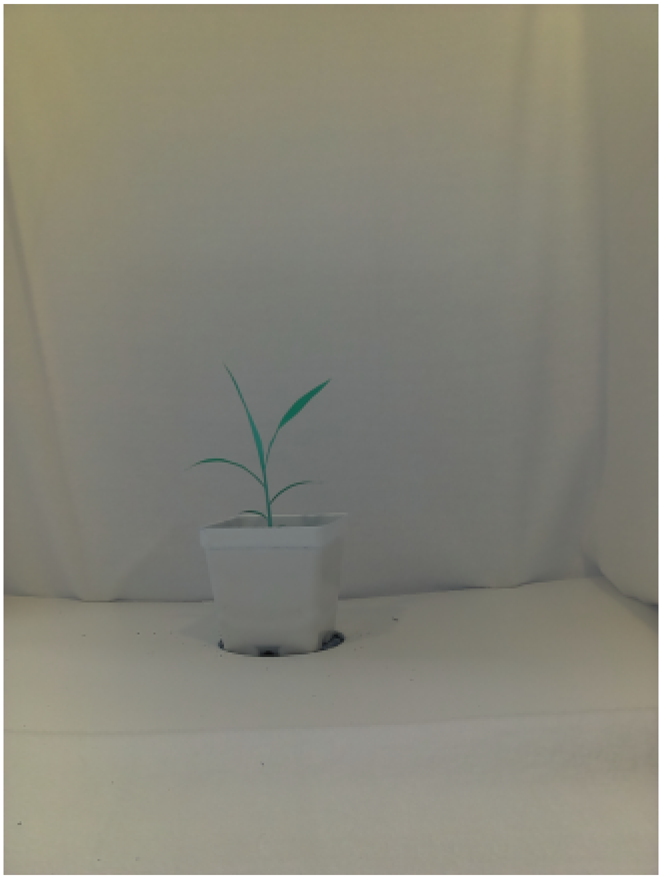

Notice in the script above that we have largely reused some code from our previous exercise, however, just to call attention to a concept we didn't discuss before notice how we can stitch together strings in python by concatenating them with the '+'. One caveat is that numeric/integer variables such as day need to be converted into a string so that python isn't confused by the operation of the '+' symbol that is desired, hence why we wrapped it in the str() function! We're almost ready to iteratively run the acute workflow but before we do that we'll need to create an empty list (but we'll at least fill it with a header at least to start).

In [4]:
```
landmark_output=[['name', 'plm_x', 'plm_y', 'SS_x', 'SS_y', 'TS_x', 'TS_y', 'CC_ratio']]
```

As we iteratively run acute on each frame we'll end up storing the landmarks generated from that day within landmark output. Note that if we didn't have this on hand (or another comparable list) before we started our variables wouldn't have anywhere to go for us to successfully save them! Now let's repeat what we learned from the previous exercise at scale...

In [5]:
```
for day in days:

    #1. Reading our image into the environment

    img = cv2.imread(path+name_prefix+str(day)+'.jpg')

    #2. Converting our RGB image into an Lab color space

    lab_img=cv2.cvtColor(img, cv2.COLOR_BGR2LAB)

    #3. Splitting our Lab image into separate color spaces

    img_l, img_a, img_b = cv2.split(lab_img)

    img_lab_channels = cv2.hconcat((img_l, img_a, img_b))

    #4. Thresholding our a and b color channels to create two masks

    #These threshold bounds will provide the best signal but feel free t
o experiment!
    a_bound = np.array([123, 255])
    b_bound = np.array([133, 255])

    #Note that we're inverting the binary threshold of color channel 'a'
so that the areas
    #with the darkest pixels will be flagged as a white mask.  This will
be important when
    #compared against the mask generated from color channel 'b'.
    mask_a = cv2.threshold(img_a, a_bound[0], a_bound[1], cv2.THRESH_BIN
ARY_INV)
    a_thresh = cv2.cvtColor(mask_a[1], cv2.COLOR_GRAY2RGB)

    mask_b = cv2.threshold(img_b, b_bound[0], b_bound[1], cv2.THRESH_BIN
ARY)
    b_thresh = cv2.cvtColor(mask_b[1], cv2.COLOR_GRAY2RGB)

    img_ab_thresholds = cv2.hconcat((a_thresh, b_thresh))

    #5. Merging our individual a and b thresholded masks

    mask=cv2.bitwise_and(mask_a[1], mask_b[1])

    #6. Extracting our contours from the final mask

    cont, hierarchy = cv2.findContours(mask, cv2.RETR_TREE, cv2.CHAIN_AP
PROX_SIMPLE)

    mask_contour = cv2.cvtColor(mask, cv2.COLOR_GRAY2RGB)

    #Find largest contour of subject (outer boundary of subject)
    cont_list = []
    hull = [0, 0]
    for c in range(len(cont)):
        a = cv2.contourArea(cont[c])
        if a > hull[0]:
            hull = [a, c]
```

```
    cont_list.append(hull[1])
    #Capture children of parent contour
    for e in range(len(hierarchy[0])):
        if (hierarchy[0][e][3] == hull[1]) & (len(cont[e]) > 10):
            cont_list.append(e)

    #7. Extracting pseudo-landmarks from the plant contours
    for l in cont_list:
        if cv2.arcLength(cont[l],True) > 2*win:
            print('Contour volume: '+str(cv2.arcLength(cont[l],True)))

            cv2.drawContours(mask, cont[l], -1, (128,0,0), 3)
            homolog_pts, homolog_start, homolog_stop, homolog_cc, chain,
verbose = acute(cont[l], mask, win, thresh, debug)
            homolog_hier = l*len(homolog_pts)
            cv2.drawContours(mask, homolog_pts, -1, (0,0,255), 3)
            print('    ' + 'landmark number: ' + str(len(homolog_pts)))

            for h in range(0, len(homolog_pts)):
                landmark_output.append([name_prefix+str(day), homolog_pt
s[h][0][0], homolog_pts[h][0][1], homolog_start[h][0][0], homolog_start[
h][0][1], homolog_stop[h][0][0], homolog_stop[h][0][1], homolog_cc[h],])

    #8. Plotting pseudo-landmarks on current image
    img_plms = img.copy()

    for c in cont_list:
        cv2.drawContours(img_plms, homolog_pts, -1, (255, 255, 255), 14)

    #Plot results
    plm_fig=plt.figure(figsize=(7, 10))
    plm_fig=plt.imshow(img_plms)
    plm_fig=plt.xscale('linear')
    plm_fig=plt.axis('off')
    plm_fig=plt.title('B100 day '+str(day))
    plt.show(plm_fig)
```

```
Contour volume: 2134.4002673625946
Fusing contour edges
route C
Landmark site:  1125  , Start site:  1113  , Term. site:  19
Landmark point indices:  [1125]
Starting site indices:  [1113]
Termination site indices:  [19]
route C
Landmark site:  203  , Start site:  188  , Term. site:  221
Landmark point indices:  [1125, 203]
Starting site indices:  [1113, 188]
Termination site indices:  [19, 221]
route C
Landmark site:  353  , Start site:  337  , Term. site:  363
Landmark point indices:  [1125, 203, 353]
Starting site indices:  [1113, 188, 337]
Termination site indices:  [19, 221, 363]
route C
Landmark site:  538  , Start site:  531  , Term. site:  547
Landmark point indices:  [1125, 203, 353, 538]
Starting site indices:  [1113, 188, 337, 531]
Termination site indices:  [19, 221, 363, 547]
route C
Landmark site:  594  , Start site:  571  , Term. site:  602
Landmark point indices:  [1125, 203, 353, 538, 594]
Starting site indices:  [1113, 188, 337, 531, 571]
Termination site indices:  [19, 221, 363, 547, 602]
route C
Landmark site:  672  , Start site:  655  , Term. site:  689
Landmark point indices:  [1125, 203, 353, 538, 594, 672]
Starting site indices:  [1113, 188, 337, 531, 571, 655]
Termination site indices:  [19, 221, 363, 547, 602, 689]
route C
Landmark site:  809  , Start site:  795  , Term. site:  824
Landmark point indices:  [1125, 203, 353, 538, 594, 672, 809]
Starting site indices:  [1113, 188, 337, 531, 571, 655, 795]
Termination site indices:  [19, 221, 363, 547, 602, 689, 824]
route C
Landmark site:  895  , Start site:  877  , Term. site:  908
Landmark point indices:  [1125, 203, 353, 538, 594, 672, 809, 895]
Starting site indices:  [1113, 188, 337, 531, 571, 655, 795, 877]
Termination site indices:  [19, 221, 363, 547, 602, 689, 824, 908]
    landmark number: 8
```

B100 day 10

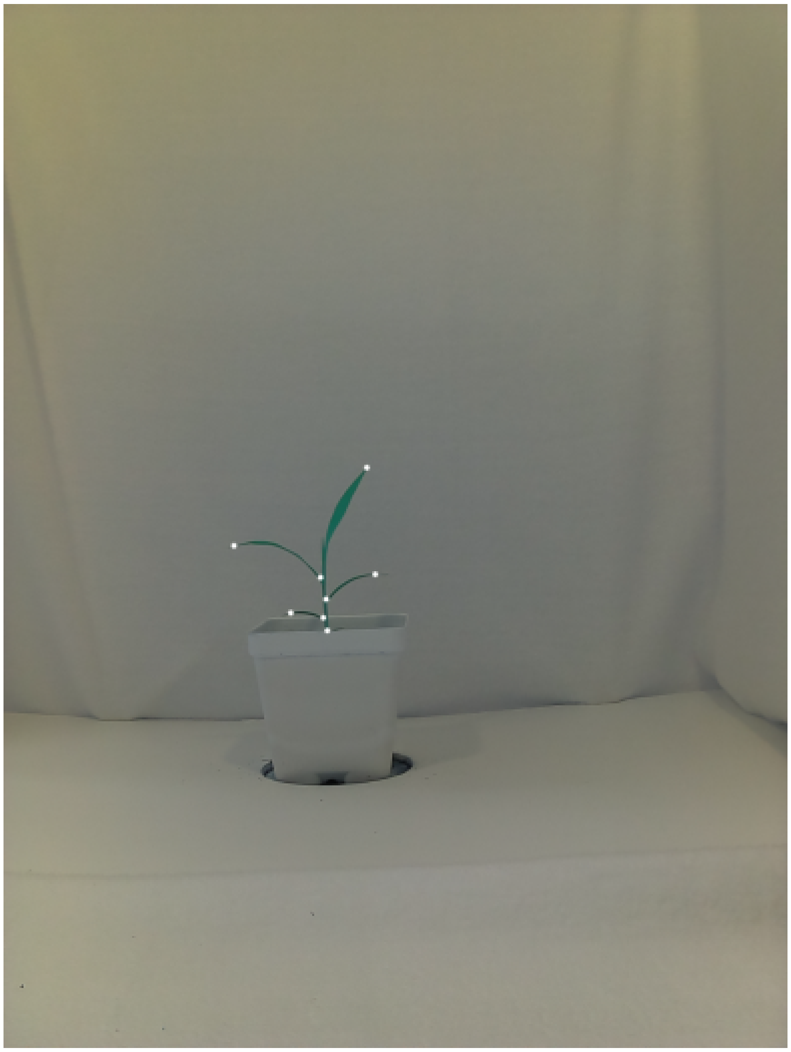

```
Contour volume: 2406.708920955658
Fusing contour edges
route C
Landmark site:  1180  , Start site:  1166  , Term. site:  7
Landmark point indices:  [1180]
Starting site indices:  [1166]
Termination site indices:  [7]
route C
Landmark site:  167  , Start site:  153  , Term. site:  180
Landmark point indices:  [1180, 167]
Starting site indices:  [1166, 153]
Termination site indices:  [7, 180]
route C
Landmark site:  237  , Start site:  224  , Term. site:  249
Landmark point indices:  [1180, 167, 237]
Starting site indices:  [1166, 153, 224]
Termination site indices:  [7, 180, 249]
route C
Landmark site:  331  , Start site:  316  , Term. site:  346
Landmark point indices:  [1180, 167, 237, 331]
Starting site indices:  [1166, 153, 224, 316]
Termination site indices:  [7, 180, 249, 346]
route C
Landmark site:  474  , Start site:  462  , Term. site:  484
Landmark point indices:  [1180, 167, 237, 331, 474]
Starting site indices:  [1166, 153, 224, 316, 462]
Termination site indices:  [7, 180, 249, 346, 484]
route C
Landmark site:  652  , Start site:  642  , Term. site:  656
Landmark point indices:  [1180, 167, 237, 331, 474, 652]
Starting site indices:  [1166, 153, 224, 316, 462, 642]
Termination site indices:  [7, 180, 249, 346, 484, 656]
route C
Landmark site:  709  , Start site:  689  , Term. site:  722
Landmark point indices:  [1180, 167, 237, 331, 474, 652, 709]
Starting site indices:  [1166, 153, 224, 316, 462, 642, 689]
Termination site indices:  [7, 180, 249, 346, 484, 656, 722]
route C
Landmark site:  782  , Start site:  759  , Term. site:  791
Landmark point indices:  [1180, 167, 237, 331, 474, 652, 709, 782]
Starting site indices:  [1166, 153, 224, 316, 462, 642, 689, 759]
Termination site indices:  [7, 180, 249, 346, 484, 656, 722, 791]
route C
Landmark site:  893  , Start site:  881  , Term. site:  906
Landmark point indices:  [1180, 167, 237, 331, 474, 652, 709, 782, 893]
Starting site indices:  [1166, 153, 224, 316, 462, 642, 689, 759, 881]
Termination site indices:  [7, 180, 249, 346, 484, 656, 722, 791, 906]
route C
Landmark site:  974  , Start site:  965  , Term. site:  982
Landmark point indices:  [1180, 167, 237, 331, 474, 652, 709, 782, 893,
974]
Starting site indices:  [1166, 153, 224, 316, 462, 642, 689, 759, 881,
965]
Termination site indices:  [7, 180, 249, 346, 484, 656, 722, 791, 906,
982]
    landmark number: 10
```

B100 day 11

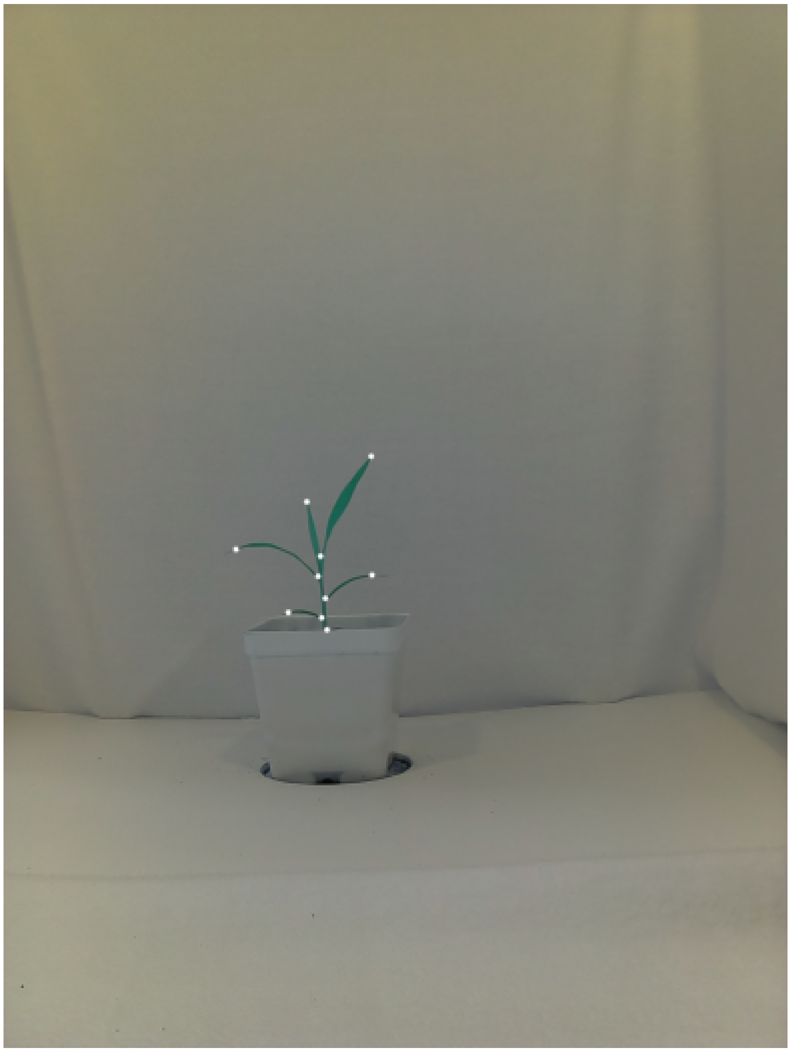

```
Contour volume: 2661.887634396553
Fusing contour edges
route C
Landmark site:  0  , Start site:  1352  , Term. site:  9
Landmark point indices:  [0]
Starting site indices:  [1352]
Termination site indices:  [9]
route C
Landmark site:  180  , Start site:  165  , Term. site:  186
Landmark point indices:  [0, 180]
Starting site indices:  [1352, 165]
Termination site indices:  [9, 186]
route C
Landmark site:  301  , Start site:  286  , Term. site:  318
Landmark point indices:  [0, 180, 301]
Starting site indices:  [1352, 165, 286]
Termination site indices:  [9, 186, 318]
route C
Landmark site:  458  , Start site:  448  , Term. site:  481
Landmark point indices:  [0, 180, 301, 458]
Starting site indices:  [1352, 165, 286, 448]
Termination site indices:  [9, 186, 318, 481]
route C
Landmark site:  610  , Start site:  601  , Term. site:  627
Landmark point indices:  [0, 180, 301, 458, 610]
Starting site indices:  [1352, 165, 286, 448, 601]
Termination site indices:  [9, 186, 318, 481, 627]
route C
Landmark site:  787  , Start site:  778  , Term. site:  800
Landmark point indices:  [0, 180, 301, 458, 610, 787]
Starting site indices:  [1352, 165, 286, 448, 601, 778]
Termination site indices:  [9, 186, 318, 481, 627, 800]
route C
Landmark site:  846  , Start site:  830  , Term. site:  851
Landmark point indices:  [0, 180, 301, 458, 610, 787, 846]
Starting site indices:  [1352, 165, 286, 448, 601, 778, 830]
Termination site indices:  [9, 186, 318, 481, 627, 800, 851]
route C
Landmark site:  902  , Start site:  889  , Term. site:  919
Landmark point indices:  [0, 180, 301, 458, 610, 787, 846, 902]
Starting site indices:  [1352, 165, 286, 448, 601, 778, 830, 889]
Termination site indices:  [9, 186, 318, 481, 627, 800, 851, 919]
route C
Landmark site:  1033  , Start site:  1015  , Term. site:  1046
Landmark point indices:  [0, 180, 301, 458, 610, 787, 846, 902, 1033]
Starting site indices:  [1352, 165, 286, 448, 601, 778, 830, 889, 1015]
Termination site indices:  [9, 186, 318, 481, 627, 800, 851, 919, 1046]
route C
Landmark site:  1127  , Start site:  1113  , Term. site:  1134
Landmark point indices:  [0, 180, 301, 458, 610, 787, 846, 902, 1033, 1
127]
Starting site indices:  [1352, 165, 286, 448, 601, 778, 830, 889, 1015,
1113]
Termination site indices:  [9, 186, 318, 481, 627, 800, 851, 919, 1046,
1134]
    landmark number: 10
```

B100 day 12

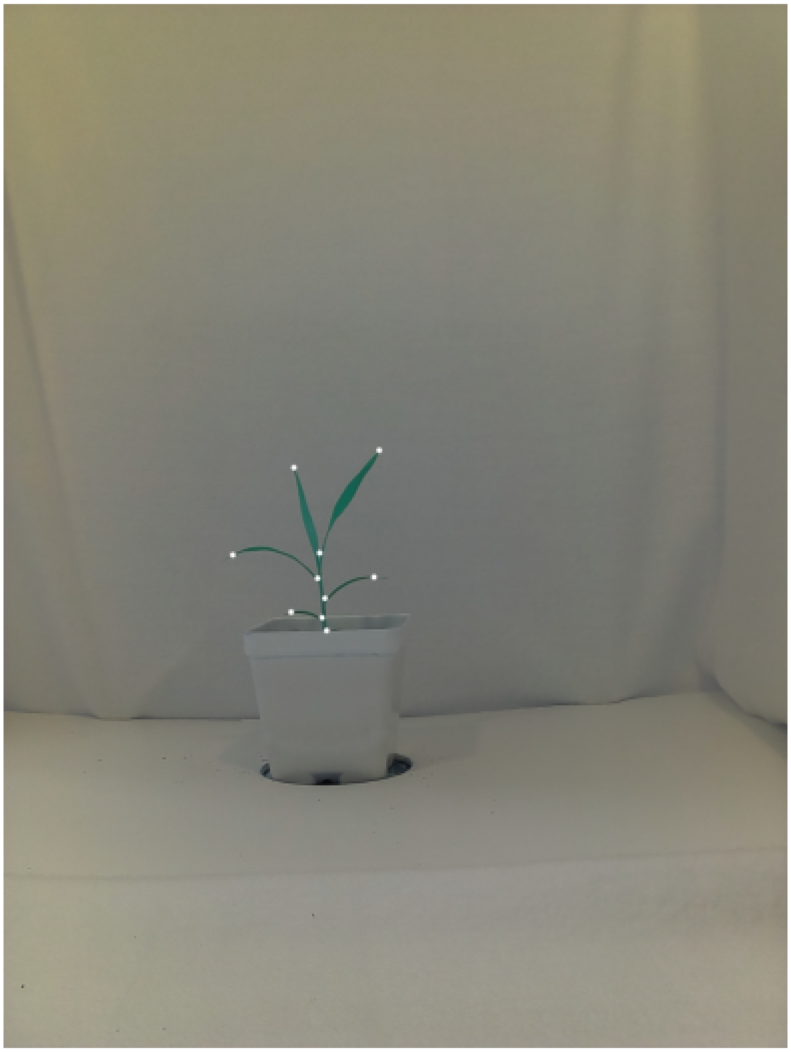

```
Contour volume: 2923.1912364959717
Fusing contour edges
route C
Landmark site:  0  , Start site:  1540  , Term. site:  13
Landmark point indices:  [0]
Starting site indices:  [1540]
Termination site indices:  [13]
route C
Landmark site:  222  , Start site:  205  , Term. site:  245
Landmark point indices:  [0, 222]
Starting site indices:  [1540, 205]
Termination site indices:  [13, 245]
route C
Landmark site:  370  , Start site:  355  , Term. site:  381
Landmark point indices:  [0, 222, 370]
Starting site indices:  [1540, 205, 355]
Termination site indices:  [13, 245, 381]
route C
Landmark site:  590  , Start site:  579  , Term. site:  598
Landmark point indices:  [0, 222, 370, 590]
Starting site indices:  [1540, 205, 355, 579]
Termination site indices:  [13, 245, 381, 598]
route C
Landmark site:  634  , Start site:  616  , Term. site:  653
Landmark point indices:  [0, 222, 370, 590, 634]
Starting site indices:  [1540, 205, 355, 579, 616]
Termination site indices:  [13, 245, 381, 598, 653]
route C
Landmark site:  708  , Start site:  690  , Term. site:  722
Landmark point indices:  [0, 222, 370, 590, 634, 708]
Starting site indices:  [1540, 205, 355, 579, 616, 690]
Termination site indices:  [13, 245, 381, 598, 653, 722]
route C
Landmark site:  833  , Start site:  832  , Term. site:  850
Landmark point indices:  [0, 222, 370, 590, 634, 708, 833]
Starting site indices:  [1540, 205, 355, 579, 616, 690, 832]
Termination site indices:  [13, 245, 381, 598, 653, 722, 850]
route C
Landmark site:  916  , Start site:  908  , Term. site:  924
Landmark point indices:  [0, 222, 370, 590, 634, 708, 833, 916]
Starting site indices:  [1540, 205, 355, 579, 616, 690, 832, 908]
Termination site indices:  [13, 245, 381, 598, 653, 722, 850, 924]
route C
Landmark site:  1173  , Start site:  1162  , Term. site:  1192
Landmark point indices:  [0, 222, 370, 590, 634, 708, 833, 916, 1173]
Starting site indices:  [1540, 205, 355, 579, 616, 690, 832, 908, 1162]
Termination site indices:  [13, 245, 381, 598, 653, 722, 850, 924, 119
2]
route C
Landmark site:  1357  , Start site:  1342  , Term. site:  1375
Landmark point indices:  [0, 222, 370, 590, 634, 708, 833, 916, 1173, 1
357]
Starting site indices:  [1540, 205, 355, 579, 616, 690, 832, 908, 1162,
1342]
Termination site indices:  [13, 245, 381, 598, 653, 722, 850, 924, 119
2, 1375]
    landmark number: 10
```

B100 day 13

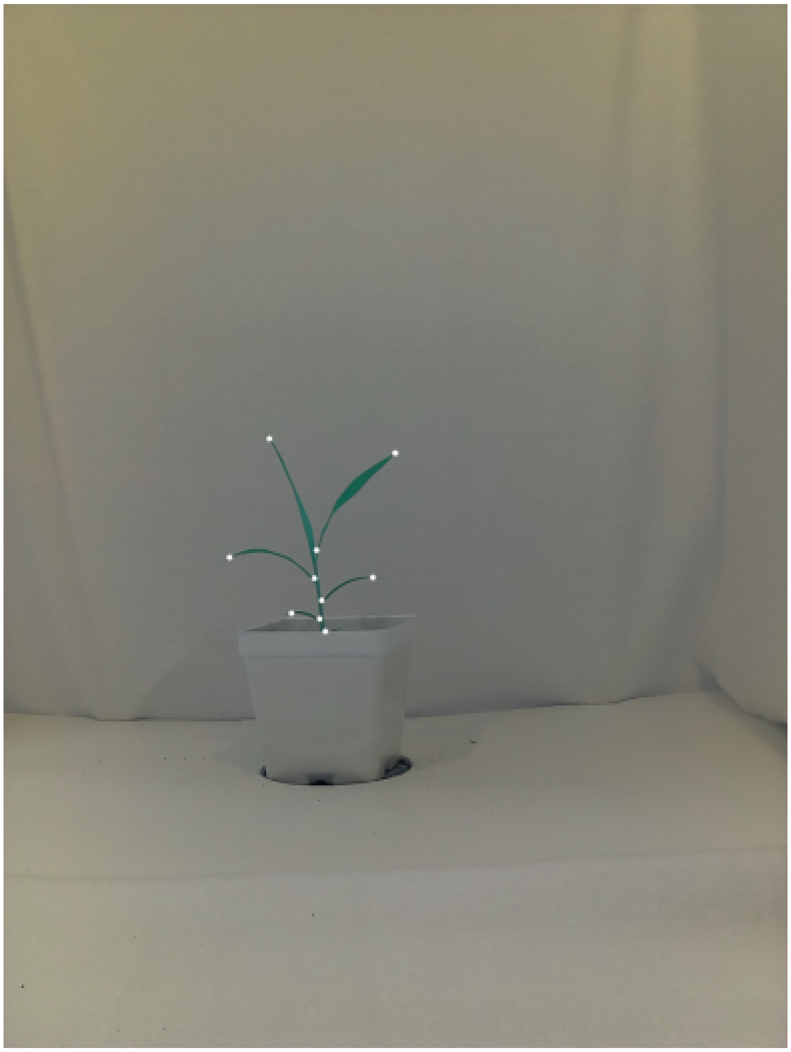

In running the code block we should have recovered a comparable series of images with the pseudo-landmarks overlaid on top of them. So how do our pseudo-landmarks look that we stored? Let's have a look!

In [6]:
```
landmark_output
```

```
Out[6]: [['name', 'plm_x', 'plm_y', 'SS_x', 'SS_y', 'TS_x', 'TS_y', 'CC_rati
o'],
 ['B100_rep1_d10', 901, 1151, 894, 1171, 885, 1167, 179.7864077669903],
 ['B100_rep1_d10', 786, 1423, 789, 1402, 772, 1404, 51.35802469135802
5],
 ['B100_rep1_d10', 571, 1344, 592, 1336, 593, 1342, 136.7586206896551
7],
 ['B100_rep1_d10', 793, 1523, 796, 1512, 783, 1519, 77.3953488372093],
 ['B100_rep1_d10', 712, 1511, 735, 1508, 734, 1512, 165.0416666666666
6],
 ['B100_rep1_d10', 803, 1555, 796, 1538, 807, 1535, 205.4915254237288],
 ['B100_rep1_d10', 922, 1415, 898, 1420, 900, 1416, 133.1836734693877
4],
 ['B100_rep1_d10', 800, 1477, 816, 1459, 801, 1454, 48.89325842696629],
 ['B100_rep1_d11', 912, 1123, 905, 1142, 896, 1139, 201.4607843137255],
 ['B100_rep1_d11', 786, 1370, 792, 1350, 783, 1347, 91.42857142857143],
 ['B100_rep1_d11', 752, 1236, 761, 1252, 754, 1260, 181.6222222222222
3],
 ['B100_rep1_d11', 780, 1420, 780, 1398, 767, 1404, 57.92700729927007],
 ['B100_rep1_d11', 576, 1353, 595, 1343, 595, 1350, 153.0151515151515],
 ['B100_rep1_d11', 789, 1523, 791, 1511, 780, 1517, 61.13432835820895
4],
 ['B100_rep1_d11', 706, 1510, 728, 1506, 728, 1510, 136.6590909090909],
 ['B100_rep1_d11', 802, 1553, 791, 1532, 801, 1533, 206.3617021276595
8],
 ['B100_rep1_d11', 914, 1417, 893, 1423, 891, 1417, 145.4313725490196],
 ['B100_rep1_d11', 798, 1475, 807, 1466, 795, 1459, 57.52808988764045],
 ['B100_rep1_d12', 932, 1108, 925, 1131, 919, 1121, 200.4018691588785],
 ['B100_rep1_d12', 784, 1362, 793, 1343, 780, 1343, 51.89189189189189
5],
 ['B100_rep1_d12', 720, 1151, 730, 1170, 724, 1175, 176.0540540540540
6],
 ['B100_rep1_d12', 779, 1425, 779, 1402, 763, 1407, 43.62011173184357],
 ['B100_rep1_d12', 568, 1367, 585, 1358, 591, 1361, 140.9152542372881
4],
 ['B100_rep1_d12', 790, 1523, 789, 1511, 778, 1518, 67.28205128205128],
 ['B100_rep1_d12', 713, 1509, 736, 1506, 719, 1511, 162.1923076923076
8],
 ['B100_rep1_d12', 801, 1555, 790, 1534, 801, 1535, 221.1728971962617],
 ['B100_rep1_d12', 919, 1422, 897, 1426, 895, 1422, 130.5918367346938
9],
 ['B100_rep1_d12', 797, 1475, 807, 1467, 793, 1461, 53.07317073170731
4],
 ['B100_rep1_d13', 659, 1079, 673, 1097, 670, 1101, 140.7],
 ['B100_rep1_d13', 771, 1425, 770, 1405, 753, 1408, 60.04938271604938],
 ['B100_rep1_d13', 560, 1373, 581, 1362, 581, 1368, 164.2857142857142
8],
 ['B100_rep1_d13', 784, 1526, 784, 1506, 771, 1518, 54.06034482758620
6],
 ['B100_rep1_d13', 714, 1512, 736, 1507, 736, 1513, 137.7692307692307
7],
 ['B100_rep1_d13', 798, 1557, 784, 1543, 796, 1539, 210.7640449438202
4],
 ['B100_rep1_d13', 916, 1423, 900, 1423, 892, 1421, 128.0],
 ['B100_rep1_d13', 789, 1480, 799, 1473, 787, 1464, 54.4367816091954],
 ['B100_rep1_d13', 971, 1115, 958, 1132, 950, 1125, 193.1858407079646
```

```
 2],
 ['B100_rep1_d13', 776, 1355, 784, 1337, 771, 1332, 64.0]]
```

This looks like it saved quite a bit more than an X-Y coordinates and a name for the file it came from... We glossed over a few of the acute outputs in the previous exercise but it's probably worth sitting down and thinking about what they are now.

If we look at the first 3 'columns' of this output we can see that there are names that correspond to our original files alongside a X-Y coordinate list which represents the plms we've been plotting. So what these other SS and TS coordinates that we seem to be storing as well? Let's review a graph we've seen before in the previous exercise and then discuss...

In [7]:
```
chain_pos=range(0, len(chain))

#Plot results
fig1=plt.plot(chain_pos, chain, color='black')
fig1=plt.axhline(y=thresh, color='r', linestyle='-')
fig1=plt.title('Angle scores by position')
plt.show(fig1)
```

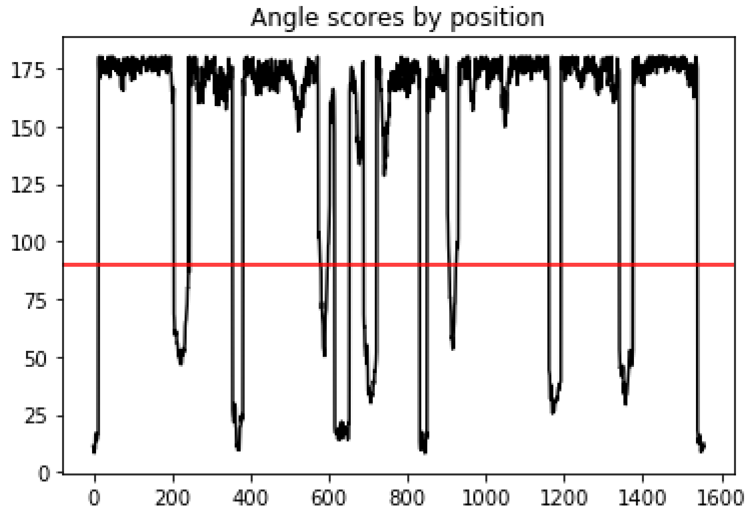


This is that waveform we've been introduced to before that defines our landmarks acute is generating. Notice that as we walk along this contour there are clearly a consecutive span of points defining an acute 'region' rather than and clear point in space (hence why we have a little valley of low angles rather than an abrupt dip)? When a pseudo-landmark is defined the midpoint of each of these valleys is taken as THE 'pseudo-landmark' and the ends of either side of these acute regions is stored as well and defined as the 'acute region start site (SS)' and the 'acute region termination site (TS)' (yes, this terminology may have been appropriated from molecular biology). So in fact we're actually storing a bit of extra spatial information than what is needed to plot as we generate our pseudo-landmarks.

Now that we solved that mystery let's run 'landmark_output' and pick up where we left off...

In [8]:

```
landmark_output
```

```
Out[8]: [['name', 'plm_x', 'plm_y', 'SS_x', 'SS_y', 'TS_x', 'TS_y', 'CC_rati
o'],
 ['B100_rep1_d10', 901, 1151, 894, 1171, 885, 1167, 179.7864077669903],
 ['B100_rep1_d10', 786, 1423, 789, 1402, 772, 1404, 51.35802469135802
5],
 ['B100_rep1_d10', 571, 1344, 592, 1336, 593, 1342, 136.7586206896551
7],
 ['B100_rep1_d10', 793, 1523, 796, 1512, 783, 1519, 77.3953488372093],
 ['B100_rep1_d10', 712, 1511, 735, 1508, 734, 1512, 165.0416666666666
6],
 ['B100_rep1_d10', 803, 1555, 796, 1538, 807, 1535, 205.4915254237288],
 ['B100_rep1_d10', 922, 1415, 898, 1420, 900, 1416, 133.1836734693877
4],
 ['B100_rep1_d10', 800, 1477, 816, 1459, 801, 1454, 48.89325842696629],
 ['B100_rep1_d11', 912, 1123, 905, 1142, 896, 1139, 201.4607843137255],
 ['B100_rep1_d11', 786, 1370, 792, 1350, 783, 1347, 91.42857142857143],
 ['B100_rep1_d11', 752, 1236, 761, 1252, 754, 1260, 181.6222222222222
3],
 ['B100_rep1_d11', 780, 1420, 780, 1398, 767, 1404, 57.92700729927007],
 ['B100_rep1_d11', 576, 1353, 595, 1343, 595, 1350, 153.0151515151515],
 ['B100_rep1_d11', 789, 1523, 791, 1511, 780, 1517, 61.13432835820895
4],
 ['B100_rep1_d11', 706, 1510, 728, 1506, 728, 1510, 136.6590909090909],
 ['B100_rep1_d11', 802, 1553, 791, 1532, 801, 1533, 206.3617021276595
8],
 ['B100_rep1_d11', 914, 1417, 893, 1423, 891, 1417, 145.4313725490196],
 ['B100_rep1_d11', 798, 1475, 807, 1466, 795, 1459, 57.52808988764045],
 ['B100_rep1_d12', 932, 1108, 925, 1131, 919, 1121, 200.4018691588785],
 ['B100_rep1_d12', 784, 1362, 793, 1343, 780, 1343, 51.89189189189189
5],
 ['B100_rep1_d12', 720, 1151, 730, 1170, 724, 1175, 176.0540540540540
6],
 ['B100_rep1_d12', 779, 1425, 779, 1402, 763, 1407, 43.62011173184357],
 ['B100_rep1_d12', 568, 1367, 585, 1358, 591, 1361, 140.9152542372881
4],
 ['B100_rep1_d12', 790, 1523, 789, 1511, 778, 1518, 67.28205128205128],
 ['B100_rep1_d12', 713, 1509, 736, 1506, 719, 1511, 162.1923076923076
8],
 ['B100_rep1_d12', 801, 1555, 790, 1534, 801, 1535, 221.1728971962617],
 ['B100_rep1_d12', 919, 1422, 897, 1426, 895, 1422, 130.5918367346938
9],
 ['B100_rep1_d12', 797, 1475, 807, 1467, 793, 1461, 53.07317073170731
4],
 ['B100_rep1_d13', 659, 1079, 673, 1097, 670, 1101, 140.7],
 ['B100_rep1_d13', 771, 1425, 770, 1405, 753, 1408, 60.04938271604938],
 ['B100_rep1_d13', 560, 1373, 581, 1362, 581, 1368, 164.2857142857142
8],
 ['B100_rep1_d13', 784, 1526, 784, 1506, 771, 1518, 54.06034482758620
6],
 ['B100_rep1_d13', 714, 1512, 736, 1507, 736, 1513, 137.7692307692307
7],
 ['B100_rep1_d13', 798, 1557, 784, 1543, 796, 1539, 210.7640449438202
4],
 ['B100_rep1_d13', 916, 1423, 900, 1423, 892, 1421, 128.0],
 ['B100_rep1_d13', 789, 1480, 799, 1473, 787, 1464, 54.4367816091954],
 ['B100_rep1_d13', 971, 1115, 958, 1132, 950, 1125, 193.1858407079646
```

```
          2],
           ['B100_rep1_d13', 776, 1355, 784, 1337, 771, 1332, 64.0]]
```

At this point the first 7 'columns' we're looking at here should make some intuitive sense for what they are representing. However, we still have one last variable left we're storing which is unclear in it's role. This is a special variable we've created within acute using the volume generated between the plm, the SS, and the TS called the 'convexity-concavity ratio' or 'CC-ratio' for short. What could this be used for? Let's take a look at our last mask again just to get a bit of context here.

In [9]:
```
#Plot results
colorized_mask = cv2.cvtColor(mask, cv2.COLOR_GRAY2RGB)
img_mask_plm = cv2.hconcat((colorized_mask, img_plms))

#Plot results
fig1=plt.figure(figsize=(21, 10))
fig1=plt.imshow(img_mask_plm, 'gray')
fig1=plt.xscale('linear')
fig1=plt.axis('off')
fig1=plt.title('Binary mask of plant and plms generated from this contou
r respectively')
plt.show(fig1)
```

Binary mask of plant and plms generated from this contour respectively

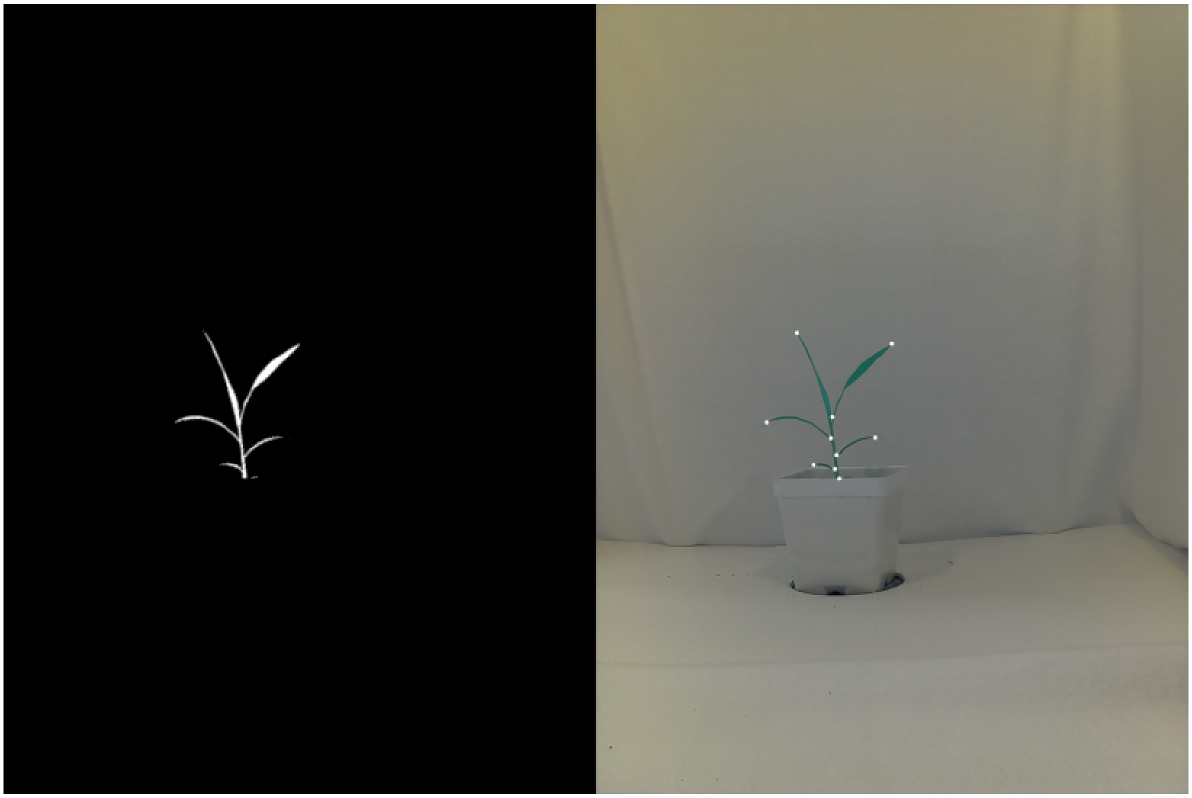

As we look at our landmarks (white points) against our mask we used to generate them notice that we're retrieving both our leaf tips and our ligules (i.e. the joints where the leaves meet the 'stem')? Now remember that our 'CC-ratio' was computed using the volume of the space between our plm, SS, and TS? If we were to draw a triangle around a leaf tip and a ligule it seems like the average pixel color could differ pretty drastically right? If we go back up and look at our CC-ratio's we'll notice that they range between 0 and 255 (the range of standard pixel intensities) with each one representing an average pixel intensity of all pixels internal to this volume we've specified. Thus, we could expect values closer to black (0) to be more common in our ligules and values closer to white (255) to be more common in our leaf tips. Thus we can use this range as a score to specify convex regions of the contour as being closer to 255 and concave regions of the contour to be closer to 0. Although it seems like we just did a bunch of extra work for no reason in generating this meta-data we'll get to see why this strategy was so important within our third and final exercise to learn the general operations of acute.

In [ ]:

```

```

In [ ]:

```

```

### Exercise 3: Generating homology groups from pseudo-landmark data

In this third exercise we will review one of the downstream applications of our batch plm data we generated in our previous exercise by generating de novo homology groups. It should be noted that this method, while incredibly powerful, has some prior assumptions in it's usage. To drive home the point, THIS METHOD IS DESIGNED TO ESTIMATE GROUPS BY MAKING ASSUMPTIONS ABOUT BIOLOGICAL HOMOLOGY (i.e. not persistent homology which is an completely different analytical method!).

Ideally, when image data of sufficient quality is presented to this workflow homology groups could even be inferred to be orthologous to one another although, similar to (phylo)genetic clustering methods, you get out what you put in and so this may be subjective based on your dataset. That being said, there are concievably two datasets where this homology grouping set is applicable:

1) Linking landmarks through time series image data to survey growth and development of independent structures through time.

2) Linking landmarks between comparable static materials either between individuals or genotypes for comparing variability of these landmarks in analogous organismal datasets (i.e. leaves with readily apparent lobes, awns, or sinuses, as one example).

Given our homology grouping workflow was designed for the former dataset we will work through a demonstration of how this works and how best to go about performing this analysis in an idealized dataset. Let's get started by importing what we need...

In [1]:
```
import cv2
import os
import sys
import pandas as pd
import numpy as np
import matplotlib.pyplot as plt

from plantcv.plantcv.homology.acute import acute
from plantcv.plantcv.homology.space import space
from plantcv.plantcv.homology.starscape import starscape
from plantcv.plantcv.homology.constella import constella

win=25
thresh=90
debug = True

path='/Path/To/Images/plm_tutorial/'
days=range(10,14)
name_prefix='B100_rep1_d'

group_iter = 1
```

**Before we get rolling though we'll have you enter in a output file path to save some graphs this workflow will generate which will be appended to our output prefix.**

In [2]:
```
outpath='/Path/To/Output/Directory/'
outfile_prefix = outpath+'B100_d10_d11_test'
```

Now that we have what we need to rerun the script we walked through in the previous exercise let's run through the code block we covered last time and then think about how best to move forward with our landmark outputs.

In [3]:
```
landmark_output=[['group', 'plmname', 'filename', 'plm_x', 'plm_y', 'SS_
x', 'SS_y', 'TS_x', 'TS_y', 'CC_ratio']]

for day in days:

    #1. Reading our image into the environment

    img = cv2.imread(path+name_prefix+str(day)+'.jpg')

    #2. Converting our RGB image into an Lab color space

    lab_img=cv2.cvtColor(img, cv2.COLOR_BGR2LAB)

    #3. Splitting our Lab image into separate color spaces

    img_l, img_a, img_b = cv2.split(lab_img)

    img_lab_channels = cv2.hconcat((img_l, img_a, img_b))

    #4. Thresholding our a and b color channels to create two masks

    #These threshold bounds will provide the best signal but feel free t
o experiment!
    a_bound = np.array([123, 255])
    b_bound = np.array([133, 255])

    #Note that we're inverting the binary threshold of color channel 'a'
so that the areas
    #with the darkest pixels will be flagged as a white mask.  This will
be important when
    #compared against the mask generated from color channel 'b'.
    mask_a = cv2.threshold(img_a, a_bound[0], a_bound[1], cv2.THRESH_BIN
ARY_INV)
    a_thresh = cv2.cvtColor(mask_a[1], cv2.COLOR_GRAY2RGB)

    mask_b = cv2.threshold(img_b, b_bound[0], b_bound[1], cv2.THRESH_BIN
ARY)
    b_thresh = cv2.cvtColor(mask_b[1], cv2.COLOR_GRAY2RGB)

    img_ab_thresholds = cv2.hconcat((a_thresh, b_thresh))

    #5. Merging our individual a and b thresholded masks

    mask=cv2.bitwise_and(mask_a[1], mask_b[1])

    #6. Extracting our contours from the final mask

    cont, hierarchy = cv2.findContours(mask, cv2.RETR_TREE, cv2.CHAIN_AP
PROX_SIMPLE)

    mask_contour = cv2.cvtColor(mask, cv2.COLOR_GRAY2RGB)

    #Find largest contour of subject (outer boundary of subject)
    cont_list = []
    hull = [0, 0]
    for c in range(len(cont)):
```

```
        a = cv2.contourArea(cont[c])
        if a > hull[0]:
            hull = [a, c]

    cont_list.append(hull[1])
    #Capture children of parent contour
    for e in range(len(hierarchy[0])):
        if (hierarchy[0][e][3] == hull[1]) & (len(cont[e]) > 10):
            cont_list.append(e)

    #7. Extracting pseudo-landmarks from the plant contours
    for l in cont_list:
        if cv2.arcLength(cont[l],True) > 2*win:
            print('Contour volume: '+str(cv2.arcLength(cont[l],True)))

            cv2.drawContours(mask, cont[l], -1, (128,0,0), 3)
            homolog_pts, homolog_start, homolog_stop, homolog_cc, chain,
verbose = acute(cont[l], mask, win, thresh, debug)
            homolog_hier = l*len(homolog_pts)
            cv2.drawContours(mask, homolog_pts, -1, (0,0,255), 3)
            print('    ' + 'landmark number: ' + str(len(homolog_pts)))

            for h in range(0, len(homolog_pts)):
                landmark_output.append([None, name_prefix+str(day)+'_pl
m'+str(h+1), name_prefix+str(day), homolog_pts[h][0][0], homolog_pts[h][
0][1], homolog_start[h][0][0], homolog_start[h][0][1], homolog_stop[h][0
][0], homolog_stop[h][0][1], homolog_cc[h],])

#Convert out output to a pandas dataframe for ease of use hereafter...
landmark_pandas=pd.DataFrame(landmark_output[1:len(landmark_output)], co
lumns=landmark_output[0][0:11])
```

```
Contour volume: 2134.4002673625946
Fusing contour edges
route C
Landmark site:  1125  , Start site:  1113  , Term. site:  19
Landmark point indices:  [1125]
Starting site indices:  [1113]
Termination site indices:  [19]
route C
Landmark site:  203  , Start site:  188  , Term. site:  221
Landmark point indices:  [1125, 203]
Starting site indices:  [1113, 188]
Termination site indices:  [19, 221]
route C
Landmark site:  353  , Start site:  337  , Term. site:  363
Landmark point indices:  [1125, 203, 353]
Starting site indices:  [1113, 188, 337]
Termination site indices:  [19, 221, 363]
route C
Landmark site:  538  , Start site:  531  , Term. site:  547
Landmark point indices:  [1125, 203, 353, 538]
Starting site indices:  [1113, 188, 337, 531]
Termination site indices:  [19, 221, 363, 547]
route C
Landmark site:  594  , Start site:  571  , Term. site:  602
Landmark point indices:  [1125, 203, 353, 538, 594]
Starting site indices:  [1113, 188, 337, 531, 571]
Termination site indices:  [19, 221, 363, 547, 602]
route C
Landmark site:  672  , Start site:  655  , Term. site:  689
Landmark point indices:  [1125, 203, 353, 538, 594, 672]
Starting site indices:  [1113, 188, 337, 531, 571, 655]
Termination site indices:  [19, 221, 363, 547, 602, 689]
route C
Landmark site:  809  , Start site:  795  , Term. site:  824
Landmark point indices:  [1125, 203, 353, 538, 594, 672, 809]
Starting site indices:  [1113, 188, 337, 531, 571, 655, 795]
Termination site indices:  [19, 221, 363, 547, 602, 689, 824]
route C
Landmark site:  895  , Start site:  877  , Term. site:  908
Landmark point indices:  [1125, 203, 353, 538, 594, 672, 809, 895]
Starting site indices:  [1113, 188, 337, 531, 571, 655, 795, 877]
Termination site indices:  [19, 221, 363, 547, 602, 689, 824, 908]
    landmark number: 8
Contour volume: 2406.708920955658
Fusing contour edges
route C
Landmark site:  1180  , Start site:  1166  , Term. site:  7
Landmark point indices:  [1180]
Starting site indices:  [1166]
Termination site indices:  [7]
route C
Landmark site:  167  , Start site:  153  , Term. site:  180
Landmark point indices:  [1180, 167]
Starting site indices:  [1166, 153]
Termination site indices:  [7, 180]
route C
Landmark site:  237  , Start site:  224  , Term. site:  249
```

```
Landmark point indices:  [1180, 167, 237]
Starting site indices:  [1166, 153, 224]
Termination site indices:  [7, 180, 249]
route C
Landmark site:  331  , Start site:  316  , Term. site:  346
Landmark point indices:  [1180, 167, 237, 331]
Starting site indices:  [1166, 153, 224, 316]
Termination site indices:  [7, 180, 249, 346]
route C
Landmark site:  474  , Start site:  462  , Term. site:  484
Landmark point indices:  [1180, 167, 237, 331, 474]
Starting site indices:  [1166, 153, 224, 316, 462]
Termination site indices:  [7, 180, 249, 346, 484]
route C
Landmark site:  652  , Start site:  642  , Term. site:  656
Landmark point indices:  [1180, 167, 237, 331, 474, 652]
Starting site indices:  [1166, 153, 224, 316, 462, 642]
Termination site indices:  [7, 180, 249, 346, 484, 656]
route C
Landmark site:  709  , Start site:  689  , Term. site:  722
Landmark point indices:  [1180, 167, 237, 331, 474, 652, 709]
Starting site indices:  [1166, 153, 224, 316, 462, 642, 689]
Termination site indices:  [7, 180, 249, 346, 484, 656, 722]
route C
Landmark site:  782  , Start site:  759  , Term. site:  791
Landmark point indices:  [1180, 167, 237, 331, 474, 652, 709, 782]
Starting site indices:  [1166, 153, 224, 316, 462, 642, 689, 759]
Termination site indices:  [7, 180, 249, 346, 484, 656, 722, 791]
route C
Landmark site:  893  , Start site:  881  , Term. site:  906
Landmark point indices:  [1180, 167, 237, 331, 474, 652, 709, 782, 893]
Starting site indices:  [1166, 153, 224, 316, 462, 642, 689, 759, 881]
Termination site indices:  [7, 180, 249, 346, 484, 656, 722, 791, 906]
route C
Landmark site:  974  , Start site:  965  , Term. site:  982
Landmark point indices:  [1180, 167, 237, 331, 474, 652, 709, 782, 893,
974]
Starting site indices:  [1166, 153, 224, 316, 462, 642, 689, 759, 881,
965]
Termination site indices:  [7, 180, 249, 346, 484, 656, 722, 791, 906,
982]
    landmark number: 10
Contour volume: 2661.887634396553
Fusing contour edges
route C
Landmark site:  0  , Start site:  1352  , Term. site:  9
Landmark point indices:  [0]
Starting site indices:  [1352]
Termination site indices:  [9]
route C
Landmark site:  180  , Start site:  165  , Term. site:  186
Landmark point indices:  [0, 180]
Starting site indices:  [1352, 165]
Termination site indices:  [9, 186]
route C
Landmark site:  301  , Start site:  286  , Term. site:  318
Landmark point indices:  [0, 180, 301]
```

```
Starting site indices:  [1352, 165, 286]
Termination site indices:  [9, 186, 318]
route C
Landmark site:  458  , Start site:  448  , Term. site:  481
Landmark point indices:  [0, 180, 301, 458]
Starting site indices:  [1352, 165, 286, 448]
Termination site indices:  [9, 186, 318, 481]
route C
Landmark site:  610  , Start site:  601  , Term. site:  627
Landmark point indices:  [0, 180, 301, 458, 610]
Starting site indices:  [1352, 165, 286, 448, 601]
Termination site indices:  [9, 186, 318, 481, 627]
route C
Landmark site:  787  , Start site:  778  , Term. site:  800
Landmark point indices:  [0, 180, 301, 458, 610, 787]
Starting site indices:  [1352, 165, 286, 448, 601, 778]
Termination site indices:  [9, 186, 318, 481, 627, 800]
route C
Landmark site:  846  , Start site:  830  , Term. site:  851
Landmark point indices:  [0, 180, 301, 458, 610, 787, 846]
Starting site indices:  [1352, 165, 286, 448, 601, 778, 830]
Termination site indices:  [9, 186, 318, 481, 627, 800, 851]
route C
Landmark site:  902  , Start site:  889  , Term. site:  919
Landmark point indices:  [0, 180, 301, 458, 610, 787, 846, 902]
Starting site indices:  [1352, 165, 286, 448, 601, 778, 830, 889]
Termination site indices:  [9, 186, 318, 481, 627, 800, 851, 919]
route C
Landmark site:  1033  , Start site:  1015  , Term. site:  1046
Landmark point indices:  [0, 180, 301, 458, 610, 787, 846, 902, 1033]
Starting site indices:  [1352, 165, 286, 448, 601, 778, 830, 889, 1015]
Termination site indices:  [9, 186, 318, 481, 627, 800, 851, 919, 1046]
route C
Landmark site:  1127  , Start site:  1113  , Term. site:  1134
Landmark point indices:  [0, 180, 301, 458, 610, 787, 846, 902, 1033, 1
127]
Starting site indices:  [1352, 165, 286, 448, 601, 778, 830, 889, 1015,
1113]
Termination site indices:  [9, 186, 318, 481, 627, 800, 851, 919, 1046,
1134]
    landmark number: 10
Contour volume: 2923.1912364959717
Fusing contour edges
route C
Landmark site:  0  , Start site:  1540  , Term. site:  13
Landmark point indices:  [0]
Starting site indices:  [1540]
Termination site indices:  [13]
route C
Landmark site:  222  , Start site:  205  , Term. site:  245
Landmark point indices:  [0, 222]
Starting site indices:  [1540, 205]
Termination site indices:  [13, 245]
route C
Landmark site:  370  , Start site:  355  , Term. site:  381
Landmark point indices:  [0, 222, 370]
Starting site indices:  [1540, 205, 355]
```

```
Termination site indices:  [13, 245, 381]
route C
Landmark site:  590  , Start site:  579  , Term. site:  598
Landmark point indices:  [0, 222, 370, 590]
Starting site indices:  [1540, 205, 355, 579]
Termination site indices:  [13, 245, 381, 598]
route C
Landmark site:  634  , Start site:  616  , Term. site:  653
Landmark point indices:  [0, 222, 370, 590, 634]
Starting site indices:  [1540, 205, 355, 579, 616]
Termination site indices:  [13, 245, 381, 598, 653]
route C
Landmark site:  708  , Start site:  690  , Term. site:  722
Landmark point indices:  [0, 222, 370, 590, 634, 708]
Starting site indices:  [1540, 205, 355, 579, 616, 690]
Termination site indices:  [13, 245, 381, 598, 653, 722]
route C
Landmark site:  833  , Start site:  832  , Term. site:  850
Landmark point indices:  [0, 222, 370, 590, 634, 708, 833]
Starting site indices:  [1540, 205, 355, 579, 616, 690, 832]
Termination site indices:  [13, 245, 381, 598, 653, 722, 850]
route C
Landmark site:  916  , Start site:  908  , Term. site:  924
Landmark point indices:  [0, 222, 370, 590, 634, 708, 833, 916]
Starting site indices:  [1540, 205, 355, 579, 616, 690, 832, 908]
Termination site indices:  [13, 245, 381, 598, 653, 722, 850, 924]
route C
Landmark site:  1173  , Start site:  1162  , Term. site:  1192
Landmark point indices:  [0, 222, 370, 590, 634, 708, 833, 916, 1173]
Starting site indices:  [1540, 205, 355, 579, 616, 690, 832, 908, 1162]
Termination site indices:  [13, 245, 381, 598, 653, 722, 850, 924, 119
2]
route C
Landmark site:  1357  , Start site:  1342  , Term. site:  1375
Landmark point indices:  [0, 222, 370, 590, 634, 708, 833, 916, 1173, 1
357]
Starting site indices:  [1540, 205, 355, 579, 616, 690, 832, 908, 1162,
1342]
Termination site indices:  [13, 245, 381, 598, 653, 722, 850, 924, 119
2, 1375]
    landmark number: 10
```

Now that we have our analyses run again let's have another look at data to think about how we'll proceed...

In [4]:
```
landmark_pandas.head()
```

Out[4]:

| | group | plmname | filename | plm_x | plm_y | SS_x | SS_y | TS_x | TS_y | CC_rati |
|---|---|---|---|---|---|---|---|---|---|---|
| **0** | None | B100_rep1_d10_plm1 | B100_rep1_d10 | 901 | 1151 | 894 | 1171 | 885 | 1167 | 179.78640 |
| **1** | None | B100_rep1_d10_plm2 | B100_rep1_d10 | 786 | 1423 | 789 | 1402 | 772 | 1404 | 51.35802 |
| **2** | None | B100_rep1_d10_plm3 | B100_rep1_d10 | 571 | 1344 | 592 | 1336 | 593 | 1342 | 136.75862 |
| **3** | None | B100_rep1_d10_plm4 | B100_rep1_d10 | 793 | 1523 | 796 | 1512 | 783 | 1519 | 77.39534 |
| **4** | None | B100_rep1_d10_plm5 | B100_rep1_d10 | 712 | 1511 | 735 | 1508 | 734 | 1512 | 165.04166 |

Thus far, we've largely been considering this data as a table where we really only cared about our X-Y coordinates that describe our plms. However, when we think about this matrix beyond the the filename and plm x/y columns we can see that we really have quite a few extra dimensions which add some context to our data. These added dimensions were originally deemed to be potentially useful for generating a rich multivariate dataset to to pull these plms together into homology groups. Space no longer is seen as a required component of this pipeline, however, given that analyses seem to only produce negligibly better results with it's inclusion. That being said, this approach does produce some novel types of metadata which could have alternative applications so we'll at least discuss what Space is doing here in it's original context, even if we gloss over it in tutorial 4. You may have also noticed we now have a new empty column we've added that didn't exist before called 'group' but for now we'll just ignore it.

To begin, let's take our initial outputs from Acute and expand them into our expanded multivariate space to use for homology grouping.

In [5]:
```
day=10

filenames=landmark_pandas.loc[:,['filename']].values
cur_plms=landmark_pandas[filenames==name_prefix+str(day)]
cur_plms=cur_plms.append(landmark_pandas[filenames==name_prefix+str(day+
1)])

cur_plms = space(cur_plms, debug=True, include_bound_dist=True, include_
centroid_dist=True, include_orient_angles=True)
```

```
  group             plmname       filename  plm_x  plm_y  SS_x  SS_y  T
S_x  \
0  None  B100_rep1_d10_plm1  B100_rep1_d10    901   1151   894  1171
885
1  None  B100_rep1_d10_plm2  B100_rep1_d10    786   1423   789  1402
772
2  None  B100_rep1_d10_plm3  B100_rep1_d10    571   1344   592  1336
593
3  None  B100_rep1_d10_plm4  B100_rep1_d10    793   1523   796  1512
783
4  None  B100_rep1_d10_plm5  B100_rep1_d10    712   1511   735  1508
734

   TS_y    CC_ratio  bot_left_dist  bot_right_dist  top_left_dist  \
0  1167  179.786408     521.647390      404.545424     331.185748
1  1404   51.358025     252.287534      189.525724     369.086711
2  1342  136.758621     211.000000      409.538765     221.000000
3  1519   77.395349     224.294449      132.909744     457.475682
4  1512  165.041667     147.705789      214.560015     412.825629

   top_right_dist  centroid_dist  orientation  centroid_orientation
0       35.000000     283.704071  -147.425943            155.422333
1      329.387310      14.317821   -15.376251             12.094757
2      414.779459     221.740840   103.091893            -72.954263
3      420.286807     114.437756   -25.016893              5.013114
4      441.184769     124.277914    92.544804           -145.159056
```

Now as we look at our outputs from the space function we can see that there is clearly quite a bit of extra information we've just added. Let's breakdown what each of these new elements are item by item just to understand what new information we've generated.

To begin we can consider five distance elements, 'bot_left_dist', 'bot_right_dist', 'top_left_dist', 'top_right_dist', and 'centroid_dist'. These new values are distances between the plms representing each row and the bounding box corners capturing our current image pairs plms. In addition, we also calculate a centroid point for our current image pair to generate a distance from the 'center of gravity' for these paired plms. Given we've largely focused on spatial positions alone distance measures, while analogous in terms of being pixel measures, help by giving us some added indication as to where in space our landmarks fall compared to one another.

Beyond these distance measures we have two other elements, 'orientation' and 'centroid_orientation'. As could be anticipated from these names these elements are both providing some additional information about the direction of the plms in space as opposed to raw distance measures, however, they are accomplishing this in very different ways. The 'orientation' measures are based purely on the plm, SS, TS coordinates in which the midpoint between SS and TS are calculated and this midpoint is then used to drive a line towards the plm to generate a slope. Following the generation of a slope an angle can be generated using the formula:

```
angle = arctan(slope)*(180/pi)
```

By contrast, the 'centroid_orientation' begins at the centroid and drives a line towards the plm to generate a slope then uses a similar formula to what was described above in order to calculate an angle of orientation.

Now that we have a multivariate dataset that is rich in context for comparisons to be made we can begin to determine how similar or distant they are to one another through time. For the initial steps we will use two approaches, PCA which is extremely useful in maximizing the amount of variation while reducing dimensionality (key in a dataset such as ours) followed by clustering approaches used to link nearest neighbors (which will help us stitch our plms together through time).

Let's begin with our PCA approach which will be found within our StarScape function...

In [6]:
```
groupA = name_prefix+str(day)
groupB = name_prefix+str(day+1)
finalDf, eigvals, loadings = starscape(cur_plms, groupA, groupB, outfile
_prefix, True)
plt.show()

finalDf.head()
```

```
Eigenvalues:  [7.64245625 4.69779072 1.43669426 0.55238408]


Var. Explained:  [0.51556252 0.31691445 0.09691985 0.03726401]


Cumul. Var. Explained:  [0.51556252 0.83247698 0.92939683 0.96666083]


3  components sufficiently informative
```

Out[6]:

| | plmname | filename | PC1 | PC2 | PC3 |
|---|---|---|---|---|---|
| **0** | B100_rep1_d10_plm1 | B100_rep1_d10 | 678.087482 | -66.393699 | -41.954341 |
| **1** | B100_rep1_d10_plm2 | B100_rep1_d10 | -42.672012 | -29.263175 | 87.104739 |
| **2** | B100_rep1_d10_plm3 | B100_rep1_d10 | -5.424657 | 464.544858 | -45.729843 |
| **3** | B100_rep1_d10_plm4 | B100_rep1_d10 | -219.879998 | -120.743091 | -36.344972 |
| **4** | B100_rep1_d10_plm5 | B100_rep1_d10 | -288.475024 | 91.257940 | -66.560096 |

Using the StarScape function above a principal component analysis is undertaken to reduce the dimensionality of our multivariate space to a minimal number of maximally informative dimensions (3 in this example) while also providing some helpful outputs for consideration as we perform our later homology grouping with Constella. When running StarScape in debugging mode as we have it should be noted that various attributes of the PCA which was performed such as the eigenvalues and eigenvectors will be printed as outputs.

The first of the graphical outputs that StarScape produces is a scree plot. The eigenvalues plotted in this graph are used to dynamically define the number of components required for explaining the relationship of our plms groupings within multivariate space. As we can observe in this scree plot, as can be expected with most PCA analyses, that the vast majority of our variance can be explained with the first few dimensions which are then stored as an output dataframe. The number of output components can be defined by the user although it is recommended to have a strong reasoning from deviating from the default setting built within this script.

Following the identification of our number of informative components we can then observe our 'starscape' as two overlaid scatter plots reflecting the first three PC dimensions. In this graph we can also observe that our two perspectives in time between this image neighbor pair are color coded allowing us to see that in fact several of these plms appear to be almost perfectly overlapping through time suggesting they likely represent the same structure. This neighbor pair was purposefully chosen for this demonstration as we can see day 11 has 2 points which appear to lack partners. This is due to the fact that a new leaf was exerted in this frame resulting in two new plms representing a leaf tip and ligule.

This PC space will provide a perfect test case for our demonstrating the methodology of our homology grouping script Constella...

In [7]:
```
cur_plms, group_iter = constella(cur_plms, finalDf, group_iter, outfile_prefix, True)
plt.show()
```

```
18 plms to group
```

Although we initially only see the hierarchical cluster used by Constella shown as a dendrogram graphic quite a bit has actually happened when we ran this function in order to generate our homology groupings!

Let's start by thinking about what our hierarchical cluster of our neighboring frames looks like in this graphic. We can see that for the vast majority of our plms there appear to be paired points which correspond to a plm from each frame (given the 3D plot from our starscape plot this probably isn't much of a surprise!). Given this initial finding it would almost seem at first glance that focusing on groups consisting of two plms would be sufficient, however, there is some nuance to plm datasets given they are dynamically describing growth as it occurs. For example, we can see at least one case in which clusters of three plms form within this dendrogram, and another more complex situation in which day 11 plm 2 becomes a rogue point in the proximity of a pair of homology groups. In each of these cases one of the emergent plms that just appeared in the day 11 frame is clustering around its nearest cluster pair in the starscape output. Even when they are no longer emergent it is often common for these new points to rapidly migrate for several days before reaching stationarity as the structure they represent grows and eventually arrests its development. As such we need a fairly robust means of describing structures which are more or less non-moving while also being able to dynamically characterize noisier subcomponents of the dataset which may be undertaking fairly rapid change for a transient period of time. Ultimately Constella is designed around the concept for describing groups as duets which are adjacent to one another in time. Let's use a series of examples to grasp this concept:

#### Constella homology grouping example (i.e. identifying duets, quartets, and rogues)

1)

```
                 --- Day 11 Group 1   |   As we look at this initial illustr
ation of a dendrogram it is clear
-----------------|                    |   that there is a clear group which
 we refer to as a 'duet' which will
                 --- Day 12 Group 1   |   share a group ID serial number dur
ing Constella de novo assignment.
```

2)

```
                 --- Day 11 Group 1   |  As development continues things oft
en become more complicated with
            ----|                     |  novel structures begin to appear an
d lacking partners due to their
           |     --- Day 12 Group 1   |  recent appearance they often cluste
r around a known duet. These
-----------|                          |  points which appear to lack any not
able partner to pair with are
            -------- Day 12 Group 2   |  referred to as rogues and are often
given their own group ID number.
```

3)

```
                 --- Day 12 Group 1   |  Development continues and further e
vidence begins to accumuluate for
            ----|                     |  group 2 with a partner now appearin
g in day 13.  However, when growth
           |     --- Day 13 Group 1   |  is rapidly occuring duets sometimes
have difficulty manifesting due
         --|                          |  to rapid changes between day 12 and
13 for group 2. This leads to a
        |  |                          |  grade luck structure as shown here
 we refer to as a quartet which is
--------|   -------- Day 12 Group 2   |  merely an artifact of a similar pro
blem known as 'long branch attract
        |                             |  -ion' in phylyogenetics. So long as
a grade of 2 plms exactly can be
         ----------- Day 13 Group 2   |  resolved a quartet can be used to a
ssign the identity of group 2.
```

4)

```
                 --- Day 13 Group 1   |  As development continues and the ra
pid growth that gave rise to the
     -----------|                     |  quartet structure abates we can beg
in to clearly resolve duets for
    |            --- Day 14 Group 1   |  groups 1 and 2.  These structured d
uets often make up the bulk of
----|                                 |  our dendrogram results as shown abo
ve which, like figure (1) can
    |        ------- Day 13 Group 2   |  readily be used to assign group ide
ntities to duets.
     -------|
             ------- Day 14 Group 2
```

In the manner described above, Constella operates through iteratively assigning points identities through an expanding nearest neighbor homology grouping scheme which is superfically similar to neighbor joining. Although these steps are critical to defining how Constella weighs homology, as important is how Constella chooses to define new serial number identities vs. perserving existing ones:

#### Constella groups: seeding vs. linking

Now that we have covered the basics of how Constella detects groups it is worth taking a moment to discuss how Constella assigns names. There are generally two strategies which largely are based on if prior encounters with plms that are being grouped through image series/time series data has occurred. When we first began this notebook we assigned the variable 'group_iter' to 1 which serves as our counter variable for assigning serial numbers to each homology group as Constella detects them. When a novel group is detected, be it a duet, graded pair in a quartet, or rogue plms Constella 'seeds' these groups by assigning them the current group_iter number and iterating the counter by one. By contrast, some groups should be expected to appear for several images in a row, especially in time series data, and in these cases an identity is already established for one of the current pair. In these cases 'linking' occurs in which the known identity for one of the pair is passed on to the yet to be defined member so that the identity of this group is allowed to be carried through time or across an image series of analogous data.

Given this naming strategy of assigning numbers as identities it is probably worth noting that although Constella is designed for use in homology-based approaches it operates in an analogous sphere to de novo genome assemblers in that although both can identify probable relationships (either as genomic scaffolds or plm linkage groups) it makes no attempt to assign known identity to these groups akin to changing scaffold identities to that of known chromosomes for a given genome. This step of defining plm groups as a specific leaf tip, a leaf axil/ligule, or a floral structure such as an inflorescence apex is a post analysis step to be undertaken by an end user.

###### Where we left off...

Now that we have a thorough understanding of exactly what we did by running running Constella it would probably be good to see how well it did wouldn't it? Let's have a look!

In [8]:
```
cur_plms
```

Out[8]:

| | group | plmname | filename | plm_x | plm_y | SS_x | SS_y | TS_x | TS_y | CC_r |
|---|---|---|---|---|---|---|---|---|---|---|
| **0** | 7 | B100_rep1_d10_plm1 | B100_rep1_d10 | 901 | 1151 | 894 | 1171 | 885 | 1167 | 179.786 |
| **1** | 8 | B100_rep1_d10_plm2 | B100_rep1_d10 | 786 | 1423 | 789 | 1402 | 772 | 1404 | 51.358 |
| **2** | 6 | B100_rep1_d10_plm3 | B100_rep1_d10 | 571 | 1344 | 592 | 1336 | 593 | 1342 | 136.758 |
| **3** | 3 | B100_rep1_d10_plm4 | B100_rep1_d10 | 793 | 1523 | 796 | 1512 | 783 | 1519 | 77.395 |
| **4** | 5 | B100_rep1_d10_plm5 | B100_rep1_d10 | 712 | 1511 | 735 | 1508 | 734 | 1512 | 165.041 |
| **5** | 1 | B100_rep1_d10_plm6 | B100_rep1_d10 | 803 | 1555 | 796 | 1538 | 807 | 1535 | 205.491 |
| **6** | 4 | B100_rep1_d10_plm7 | B100_rep1_d10 | 922 | 1415 | 898 | 1420 | 900 | 1416 | 133.183 |
| **7** | 2 | B100_rep1_d10_plm8 | B100_rep1_d10 | 800 | 1477 | 816 | 1459 | 801 | 1454 | 48.893 |
| **8** | 7 | B100_rep1_d11_plm1 | B100_rep1_d11 | 912 | 1123 | 905 | 1142 | 896 | 1139 | 201.460 |
| **9** | 9 | B100_rep1_d11_plm2 | B100_rep1_d11 | 786 | 1370 | 792 | 1350 | 783 | 1347 | 91.428 |
| **10** | 10 | B100_rep1_d11_plm3 | B100_rep1_d11 | 752 | 1236 | 761 | 1252 | 754 | 1260 | 181.622 |
| **11** | 8 | B100_rep1_d11_plm4 | B100_rep1_d11 | 780 | 1420 | 780 | 1398 | 767 | 1404 | 57.927 |
| **12** | 6 | B100_rep1_d11_plm5 | B100_rep1_d11 | 576 | 1353 | 595 | 1343 | 595 | 1350 | 153.015 |
| **13** | 3 | B100_rep1_d11_plm6 | B100_rep1_d11 | 789 | 1523 | 791 | 1511 | 780 | 1517 | 61.134 |
| **14** | 5 | B100_rep1_d11_plm7 | B100_rep1_d11 | 706 | 1510 | 728 | 1506 | 728 | 1510 | 136.659 |
| **15** | 1 | B100_rep1_d11_plm8 | B100_rep1_d11 | 802 | 1553 | 791 | 1532 | 801 | 1533 | 206.361 |
| **16** | 4 | B100_rep1_d11_plm9 | B100_rep1_d11 | 914 | 1417 | 893 | 1423 | 891 | 1417 | 145.431 |
| **17** | 2 | B100_rep1_d11_plm10 | B100_rep1_d11 | 798 | 1475 | 807 | 1466 | 795 | 1459 | 57.528 |

It definitely appears as if we have paired groups between the majority of our plms across days 10 and 11! The only exceptions appear to be two plms specific to day 11 which are assigned to groups 9 and 10. It would probably be worth seeing how these stack up on our original data (i.e. the images) since these data tables are often aren't the easiest to process. With that being said, let's superimpose these groups onto the plms coordinates on each frame to see if they are in agreement.

In [9]:
```
img1 = cv2.imread(path+name_prefix+str(day)+'.jpg')

for p in range(0,cur_plms.shape[0]):
    if name_prefix+str(day) in cur_plms.at[p, 'plmname']:
        cv2.putText(img1, str(cur_plms.at[p, 'group']),
                    (int(cur_plms.at[p, 'plm_x'])-10, int(cur_plms.at[p,
'plm_y'])),
                    cv2.FONT_ITALIC, 1.5, (255,0,0), 6)

img2 = cv2.imread(path+name_prefix+str(day+1)+'.jpg')

for p in range(0,cur_plms.shape[0]):
    if name_prefix+str(day+1) in cur_plms.at[p, 'plmname']:
        cv2.putText(img2, str(cur_plms.at[p, 'group']),
                    (int(cur_plms.at[p, 'plm_x'])-10, int(cur_plms.at[p,
'plm_y'])),
                    cv2.FONT_ITALIC, 1.5, (0,0,255), 6)

img_neighbors = cv2.hconcat((img1, img2))

plm_groups_fig=plt.figure(figsize=(16, 12))
plm_groups_fig=plt.imshow(img_neighbors)
plm_groups_fig=plt.xscale('linear')
plm_groups_fig=plt.axis('off')
plm_groups_fig=plt.title('B100 day '+str(day)+'-'+str(day+1))
plt.show(plm_groups_fig)
```

B100 day 10-11

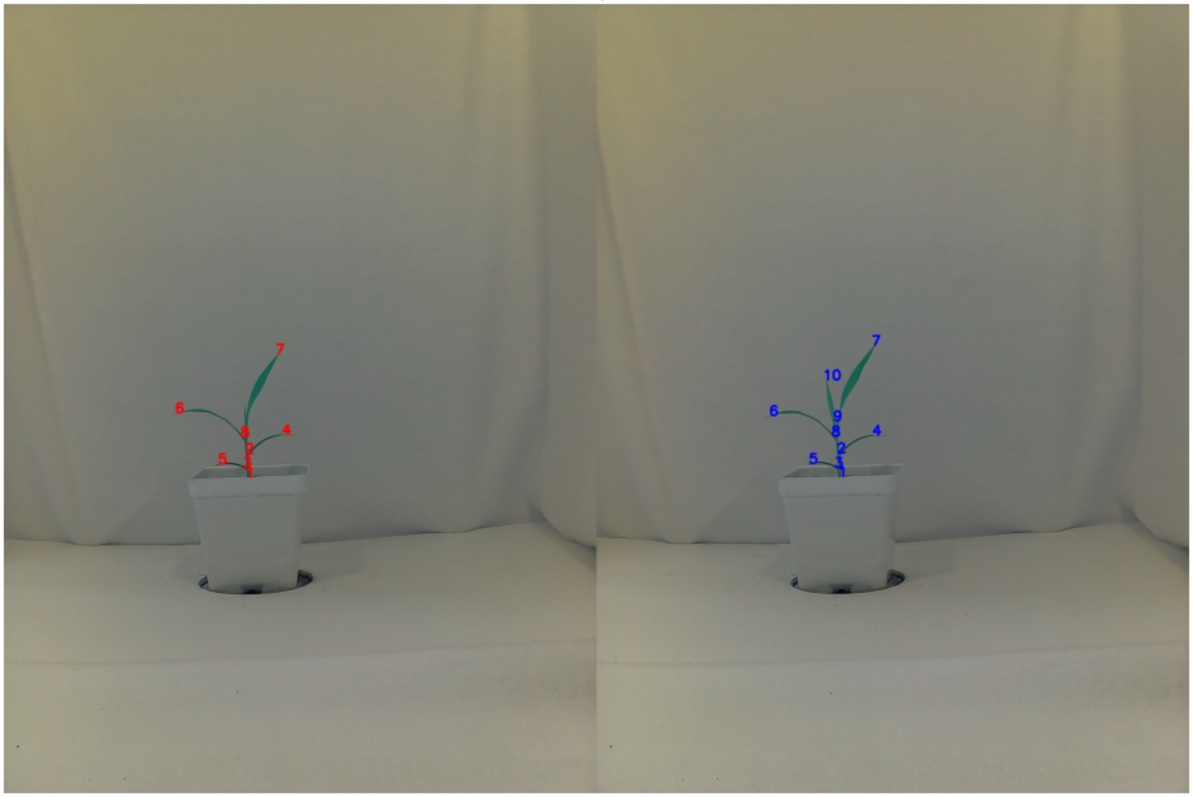

Looking at our groups overlaid against the leaf tips and ligules it seems like our attempts at forming homology groups through our workflow was a success! And note how our ligule and leaf tip plms corresponding to the emergent leaf in day 11 are represented by groups '9' and '10' which didn't appear in our first frame, seeding new groups as novel structures appear is clearly working as advertised as well!

Now that we understand how homology grouping works through the use of our Space >>> StarScape >>> Constella workflow we will use our final exercise to expand on what we've learned and apply it to store time series data and utilize groundtruthed plms of QC steps during pipeline development.

### Exercise 4: Batch homology grouping and downstream QC analyses

Following on the same strategy we employed in exercises 1 and 2 of first learning how to employ acute on a single image and then scaling up to batch image data we will now take what we have learned in exercise 3 for homology grouping with the StarScape and Constella workflow and scale this method up for use on batch image data. Following the generation of serial ID number homology groups and assigning them to our acute plms we will then assay the accuracy of these results through the use of a Quality Control (QC) test for Constella and discuss how these outputs should be interpreted.

To begin lets load the libraries and other input files we'll need to proceed...

In [1]:
```
import cv2
import os
import sys
import pandas as pd
import numpy as np
import matplotlib.pyplot as plt
import plantcv as pcv

from plantcv.plantcv.homology.acute import acute
from plantcv.plantcv.homology.space import space
from plantcv.plantcv.homology.starscape import starscape
from plantcv.plantcv.homology.constella import constella
from plantcv.plantcv.homology.constellaqc import constellaqc

win=25
thresh=90
debug = True

path='/Path/To/Images/plm_tutorial/'
days=range(10,14)
name_prefix='B100_rep1_d'
outpath='/Path/To/Output/Directory/'
outfile_prefix = outpath+'B100_test'

group_iter = 1
```

Now with our libraries loaded and initial parameters assigned let us begin by running a batch workflow with acute to generate our list of plms as a landmark dataframe.

In [2]:
```
landmark_output=[['group', 'plmname', 'filename', 'plm_x', 'plm_y', 'SS_
x', 'SS_y', 'TS_x', 'TS_y', 'CC_ratio']]

for day in days:

    #1. Reading our image into the environment

    img = cv2.imread(path+name_prefix+str(day)+'.jpg')

    #2. Converting our RGB image into an Lab color space

    lab_img=cv2.cvtColor(img, cv2.COLOR_BGR2LAB)

    #3. Splitting our Lab image into separate color spaces

    img_l, img_a, img_b = cv2.split(lab_img)

    img_lab_channels = cv2.hconcat((img_l, img_a, img_b))

    #4. Thresholding our a and b color channels to create two masks

    #These threshold bounds will provide the best signal but feel free t
o experiment!
    a_bound = np.array([123, 255])
    b_bound = np.array([133, 255])

    #Note that we're inverting the binary threshold of color channel 'a'
so that the areas
    #with the darkest pixels will be flagged as a white mask.  This will
be important when
    #compared against the mask generated from color channel 'b'.
    mask_a = cv2.threshold(img_a, a_bound[0], a_bound[1], cv2.THRESH_BIN
ARY_INV)
    a_thresh = cv2.cvtColor(mask_a[1], cv2.COLOR_GRAY2RGB)

    mask_b = cv2.threshold(img_b, b_bound[0], b_bound[1], cv2.THRESH_BIN
ARY)
    b_thresh = cv2.cvtColor(mask_b[1], cv2.COLOR_GRAY2RGB)

    img_ab_thresholds = cv2.hconcat((a_thresh, b_thresh))

    #5. Merging our individual a and b thresholded masks

    mask=cv2.bitwise_and(mask_a[1], mask_b[1])

    #6. Extracting our contours from the final mask

    cont, hierarchy = cv2.findContours(mask, cv2.RETR_TREE, cv2.CHAIN_AP
PROX_SIMPLE)

    mask_contour = cv2.cvtColor(mask, cv2.COLOR_GRAY2RGB)

    #Find largest contour of subject (outer boundary of subject)
    cont_list = []
    hull = [0, 0]
    for c in range(len(cont)):
```

```
        a = cv2.contourArea(cont[c])
        if a > hull[0]:
            hull = [a, c]

    cont_list.append(hull[1])
    #Capture children of parent contour
    for e in range(len(hierarchy[0])):
        if (hierarchy[0][e][3] == hull[1]) & (len(cont[e]) > 10):
            cont_list.append(e)

    #7. Extracting pseudo-landmarks from the plant contours
    for l in cont_list:
        if cv2.arcLength(cont[l],True) > 2*win:
            print('Contour volume: '+str(cv2.arcLength(cont[l],True)))

            cv2.drawContours(mask, cont[l], -1, (128,0,0), 3)
            homolog_pts, homolog_start, homolog_stop, homolog_cc, chain,
verbose = acute(cont[l], mask, win, thresh, debug)
            homolog_hier = l*len(homolog_pts)
            cv2.drawContours(mask, homolog_pts, -1, (0,0,255), 3)
            print('    ' + 'landmark number: ' + str(len(homolog_pts)))

            for h in range(0, len(homolog_pts)):
                landmark_output.append([None, name_prefix+str(day)+'_pl
m'+str(h+1), name_prefix+str(day), homolog_pts[h][0][0], homolog_pts[h][
0][1], homolog_start[h][0][0], homolog_start[h][0][1], homolog_stop[h][0
][0], homolog_stop[h][0][1], homolog_cc[h],])

#Convert out output to a pandas dataframe for ease of use hereafter...
landmark_pandas=pd.DataFrame(landmark_output[1:len(landmark_output)], co
lumns=landmark_output[0][0:11])
```

```
Contour volume: 2134.4002673625946
Fusing contour edges
route C
Landmark site:  1125  , Start site:  1113  , Term. site:  19
Landmark point indices:  [1125]
Starting site indices:  [1113]
Termination site indices:  [19]
route C
Landmark site:  203  , Start site:  188  , Term. site:  221
Landmark point indices:  [1125, 203]
Starting site indices:  [1113, 188]
Termination site indices:  [19, 221]
route C
Landmark site:  353  , Start site:  337  , Term. site:  363
Landmark point indices:  [1125, 203, 353]
Starting site indices:  [1113, 188, 337]
Termination site indices:  [19, 221, 363]
route C
Landmark site:  538  , Start site:  531  , Term. site:  547
Landmark point indices:  [1125, 203, 353, 538]
Starting site indices:  [1113, 188, 337, 531]
Termination site indices:  [19, 221, 363, 547]
route C
Landmark site:  594  , Start site:  571  , Term. site:  602
Landmark point indices:  [1125, 203, 353, 538, 594]
Starting site indices:  [1113, 188, 337, 531, 571]
Termination site indices:  [19, 221, 363, 547, 602]
route C
Landmark site:  672  , Start site:  655  , Term. site:  689
Landmark point indices:  [1125, 203, 353, 538, 594, 672]
Starting site indices:  [1113, 188, 337, 531, 571, 655]
Termination site indices:  [19, 221, 363, 547, 602, 689]
route C
Landmark site:  809  , Start site:  795  , Term. site:  824
Landmark point indices:  [1125, 203, 353, 538, 594, 672, 809]
Starting site indices:  [1113, 188, 337, 531, 571, 655, 795]
Termination site indices:  [19, 221, 363, 547, 602, 689, 824]
route C
Landmark site:  895  , Start site:  877  , Term. site:  908
Landmark point indices:  [1125, 203, 353, 538, 594, 672, 809, 895]
Starting site indices:  [1113, 188, 337, 531, 571, 655, 795, 877]
Termination site indices:  [19, 221, 363, 547, 602, 689, 824, 908]
    landmark number: 8
Contour volume: 2406.708920955658
Fusing contour edges
route C
Landmark site:  1180  , Start site:  1166  , Term. site:  7
Landmark point indices:  [1180]
Starting site indices:  [1166]
Termination site indices:  [7]
route C
Landmark site:  167  , Start site:  153  , Term. site:  180
Landmark point indices:  [1180, 167]
Starting site indices:  [1166, 153]
Termination site indices:  [7, 180]
route C
Landmark site:  237  , Start site:  224  , Term. site:  249
```

```
Landmark point indices:  [1180, 167, 237]
Starting site indices:  [1166, 153, 224]
Termination site indices:  [7, 180, 249]
route C
Landmark site:  331  , Start site:  316  , Term. site:  346
Landmark point indices:  [1180, 167, 237, 331]
Starting site indices:  [1166, 153, 224, 316]
Termination site indices:  [7, 180, 249, 346]
route C
Landmark site:  474  , Start site:  462  , Term. site:  484
Landmark point indices:  [1180, 167, 237, 331, 474]
Starting site indices:  [1166, 153, 224, 316, 462]
Termination site indices:  [7, 180, 249, 346, 484]
route C
Landmark site:  652  , Start site:  642  , Term. site:  656
Landmark point indices:  [1180, 167, 237, 331, 474, 652]
Starting site indices:  [1166, 153, 224, 316, 462, 642]
Termination site indices:  [7, 180, 249, 346, 484, 656]
route C
Landmark site:  709  , Start site:  689  , Term. site:  722
Landmark point indices:  [1180, 167, 237, 331, 474, 652, 709]
Starting site indices:  [1166, 153, 224, 316, 462, 642, 689]
Termination site indices:  [7, 180, 249, 346, 484, 656, 722]
route C
Landmark site:  782  , Start site:  759  , Term. site:  791
Landmark point indices:  [1180, 167, 237, 331, 474, 652, 709, 782]
Starting site indices:  [1166, 153, 224, 316, 462, 642, 689, 759]
Termination site indices:  [7, 180, 249, 346, 484, 656, 722, 791]
route C
Landmark site:  893  , Start site:  881  , Term. site:  906
Landmark point indices:  [1180, 167, 237, 331, 474, 652, 709, 782, 893]
Starting site indices:  [1166, 153, 224, 316, 462, 642, 689, 759, 881]
Termination site indices:  [7, 180, 249, 346, 484, 656, 722, 791, 906]
route C
Landmark site:  974  , Start site:  965  , Term. site:  982
Landmark point indices:  [1180, 167, 237, 331, 474, 652, 709, 782, 893,
974]
Starting site indices:  [1166, 153, 224, 316, 462, 642, 689, 759, 881,
965]
Termination site indices:  [7, 180, 249, 346, 484, 656, 722, 791, 906,
982]
    landmark number: 10
Contour volume: 2661.887634396553
Fusing contour edges
route C
Landmark site:  0  , Start site:  1352  , Term. site:  9
Landmark point indices:  [0]
Starting site indices:  [1352]
Termination site indices:  [9]
route C
Landmark site:  180  , Start site:  165  , Term. site:  186
Landmark point indices:  [0, 180]
Starting site indices:  [1352, 165]
Termination site indices:  [9, 186]
route C
Landmark site:  301  , Start site:  286  , Term. site:  318
Landmark point indices:  [0, 180, 301]
```

```
Starting site indices:  [1352, 165, 286]
Termination site indices:  [9, 186, 318]
route C
Landmark site:  458  , Start site:  448  , Term. site:  481
Landmark point indices:  [0, 180, 301, 458]
Starting site indices:  [1352, 165, 286, 448]
Termination site indices:  [9, 186, 318, 481]
route C
Landmark site:  610  , Start site:  601  , Term. site:  627
Landmark point indices:  [0, 180, 301, 458, 610]
Starting site indices:  [1352, 165, 286, 448, 601]
Termination site indices:  [9, 186, 318, 481, 627]
route C
Landmark site:  787  , Start site:  778  , Term. site:  800
Landmark point indices:  [0, 180, 301, 458, 610, 787]
Starting site indices:  [1352, 165, 286, 448, 601, 778]
Termination site indices:  [9, 186, 318, 481, 627, 800]
route C
Landmark site:  846  , Start site:  830  , Term. site:  851
Landmark point indices:  [0, 180, 301, 458, 610, 787, 846]
Starting site indices:  [1352, 165, 286, 448, 601, 778, 830]
Termination site indices:  [9, 186, 318, 481, 627, 800, 851]
route C
Landmark site:  902  , Start site:  889  , Term. site:  919
Landmark point indices:  [0, 180, 301, 458, 610, 787, 846, 902]
Starting site indices:  [1352, 165, 286, 448, 601, 778, 830, 889]
Termination site indices:  [9, 186, 318, 481, 627, 800, 851, 919]
route C
Landmark site:  1033  , Start site:  1015  , Term. site:  1046
Landmark point indices:  [0, 180, 301, 458, 610, 787, 846, 902, 1033]
Starting site indices:  [1352, 165, 286, 448, 601, 778, 830, 889, 1015]
Termination site indices:  [9, 186, 318, 481, 627, 800, 851, 919, 1046]
route C
Landmark site:  1127  , Start site:  1113  , Term. site:  1134
Landmark point indices:  [0, 180, 301, 458, 610, 787, 846, 902, 1033, 1
127]
Starting site indices:  [1352, 165, 286, 448, 601, 778, 830, 889, 1015,
1113]
Termination site indices:  [9, 186, 318, 481, 627, 800, 851, 919, 1046,
1134]
    landmark number: 10
Contour volume: 2923.1912364959717
Fusing contour edges
route C
Landmark site:  0  , Start site:  1540  , Term. site:  13
Landmark point indices:  [0]
Starting site indices:  [1540]
Termination site indices:  [13]
route C
Landmark site:  222  , Start site:  205  , Term. site:  245
Landmark point indices:  [0, 222]
Starting site indices:  [1540, 205]
Termination site indices:  [13, 245]
route C
Landmark site:  370  , Start site:  355  , Term. site:  381
Landmark point indices:  [0, 222, 370]
Starting site indices:  [1540, 205, 355]
```

```
Termination site indices:  [13, 245, 381]
route C
Landmark site:  590  , Start site:  579  , Term. site:  598
Landmark point indices:  [0, 222, 370, 590]
Starting site indices:  [1540, 205, 355, 579]
Termination site indices:  [13, 245, 381, 598]
route C
Landmark site:  634  , Start site:  616  , Term. site:  653
Landmark point indices:  [0, 222, 370, 590, 634]
Starting site indices:  [1540, 205, 355, 579, 616]
Termination site indices:  [13, 245, 381, 598, 653]
route C
Landmark site:  708  , Start site:  690  , Term. site:  722
Landmark point indices:  [0, 222, 370, 590, 634, 708]
Starting site indices:  [1540, 205, 355, 579, 616, 690]
Termination site indices:  [13, 245, 381, 598, 653, 722]
route C
Landmark site:  833  , Start site:  832  , Term. site:  850
Landmark point indices:  [0, 222, 370, 590, 634, 708, 833]
Starting site indices:  [1540, 205, 355, 579, 616, 690, 832]
Termination site indices:  [13, 245, 381, 598, 653, 722, 850]
route C
Landmark site:  916  , Start site:  908  , Term. site:  924
Landmark point indices:  [0, 222, 370, 590, 634, 708, 833, 916]
Starting site indices:  [1540, 205, 355, 579, 616, 690, 832, 908]
Termination site indices:  [13, 245, 381, 598, 653, 722, 850, 924]
route C
Landmark site:  1173  , Start site:  1162  , Term. site:  1192
Landmark point indices:  [0, 222, 370, 590, 634, 708, 833, 916, 1173]
Starting site indices:  [1540, 205, 355, 579, 616, 690, 832, 908, 1162]
Termination site indices:  [13, 245, 381, 598, 653, 722, 850, 924, 119
2]
route C
Landmark site:  1357  , Start site:  1342  , Term. site:  1375
Landmark point indices:  [0, 222, 370, 590, 634, 708, 833, 916, 1173, 1
357]
Starting site indices:  [1540, 205, 355, 579, 616, 690, 832, 908, 1162,
1342]
Termination site indices:  [13, 245, 381, 598, 653, 722, 850, 924, 119
2, 1375]
    landmark number: 10
```

Before we continue lets check our landmarks once more to ensure we'll have what we need to run our homology grouping workflow...

In [3]:
```
landmark_pandas.head()
```

Out[3]:

| | group | plmname | filename | plm_x | plm_y | SS_x | SS_y | TS_x | TS_y | CC_rati |
|---|---|---|---|---|---|---|---|---|---|---|
| 0 | None | B100_rep1_d10_plm1 | B100_rep1_d10 | 901 | 1151 | 894 | 1171 | 885 | 1167 | 179.78640 |
| 1 | None | B100_rep1_d10_plm2 | B100_rep1_d10 | 786 | 1423 | 789 | 1402 | 772 | 1404 | 51.35802 |
| 2 | None | B100_rep1_d10_plm3 | B100_rep1_d10 | 571 | 1344 | 592 | 1336 | 593 | 1342 | 136.75862 |
| 3 | None | B100_rep1_d10_plm4 | B100_rep1_d10 | 793 | 1523 | 796 | 1512 | 783 | 1519 | 77.39534 |
| 4 | None | B100_rep1_d10_plm5 | B100_rep1_d10 | 712 | 1511 | 735 | 1508 | 734 | 1512 | 165.04166 |

Provided the head of our table loaded in properly we're now ready to begin a batch run of our homology pipeline which follows the same structure as our previous exercise utilizing StarScape and Constella. Note that we have left out Space from this workflow, studies of this pipelines accuracy have suggested that this function isn't essential for generating tangible improvements in homology grouping. As such this workflow of parsing segmented morphological data directly into Starscape is considered the present best practice of this approach. Notice how our groups within landmark_pandas are universally assigned to 'None'? On the other side of this code block we should see the results of transfering group IDs from paired frames to this original dataframe resulting in forthcoming changes to this columns values. Let's get started!

In [4]:
```
for di in range(0,len(days)-1):

    print('\nBeginning next iteration for days '+str(days[di])+' and '+s
tr(days[di]+1)+'\n')
    filenames=landmark_pandas.loc[:,['filename']].values
    cur_plms=landmark_pandas[filenames==name_prefix+str(days[di])]
    cur_plms=cur_plms.append(landmark_pandas[filenames==name_prefix+str(
days[di]+1)])
    cur_plms

    groupA = name_prefix+str(days[di])
    groupB = name_prefix+str(days[di]+1)

    print('\nRunning StarScape...\n')
    finalDf, eigenvals, loadings = starscape(cur_plms, groupA, groupB,
'B100_rep1_d'+str(days[di])+'_test', True)
    plt.show()

    print('\nRunning Constella...\n')
    cur_plms, group_iter = constella(cur_plms, finalDf, group_iter, 'B10
0_rep1_d'+str(days[di])+'_test', True)
    plt.show()

    plmnames=landmark_pandas.loc[:,['plmname']].values
    cur_plmnames=cur_plms.loc[:,['plmname']].values

    for name in cur_plmnames:
        landmark_index=[i for i, x in enumerate(plmnames==name) if x]
        cur_plms_index=[i for i, x in enumerate(cur_plmnames==name) if x
]
        if landmark_pandas.iloc[landmark_index,0].values == None:
            landmark_pandas.iloc[landmark_index,0] = cur_plms.iloc[cur_p
lms_index,0]

    if 1==1:
        img1 = cv2.imread(path+name_prefix+str(days[di])+'.jpg')

        for p in range(0,cur_plms.shape[0]):
            if name_prefix+str(days[di]) in cur_plms.iloc[p, 2]:
                cv2.putText(img1, str(cur_plms.iloc[p, 0]),
                            (int(cur_plms.iloc[p, 3])-10, int(cur_plms.i
loc[p, 4])),
                            cv2.FONT_ITALIC, 1.5, (255,0,0), 6)

        img2 = cv2.imread(path+name_prefix+str(days[di]+1)+'.jpg')

        for p in range(0,cur_plms.shape[0]):
            if name_prefix+str(days[di]+1) in cur_plms.iloc[p, 2]:
                cv2.putText(img2, str(cur_plms.iloc[p, 0]),
                            (int(cur_plms.iloc[p, 3])-10, int(cur_plms.i
loc[p, 4])),
                            cv2.FONT_ITALIC, 1.5, (0,0,255), 6)

        img_neighbors = cv2.hconcat((img1, img2))

        plm_groups_fig=plt.figure(figsize=(16, 12))
```

```
        plm_groups_fig=plt.imshow(img_neighbors)
        plm_groups_fig=plt.xscale('linear')
        plm_groups_fig=plt.axis('off')
        plm_groups_fig=plt.title('B100 day '+str(days[di])+'-'+str(days[
di]+1))
        plt.show(plm_groups_fig)
```

Beginning next iteration for days 10 and 11

Running StarScape...

Eigenvalues:  [3.80272283 2.68008038 0.92036756 0.00479015]

Var. Explained:  [0.51306578 0.36159815 0.12417658 0.00064629]

Cumul. Var. Explained:  [0.51306578 0.87466393 0.9988405  0.99948679]

2  components sufficiently informative

Running Constella...

18 plms to group

B100 day 10-11

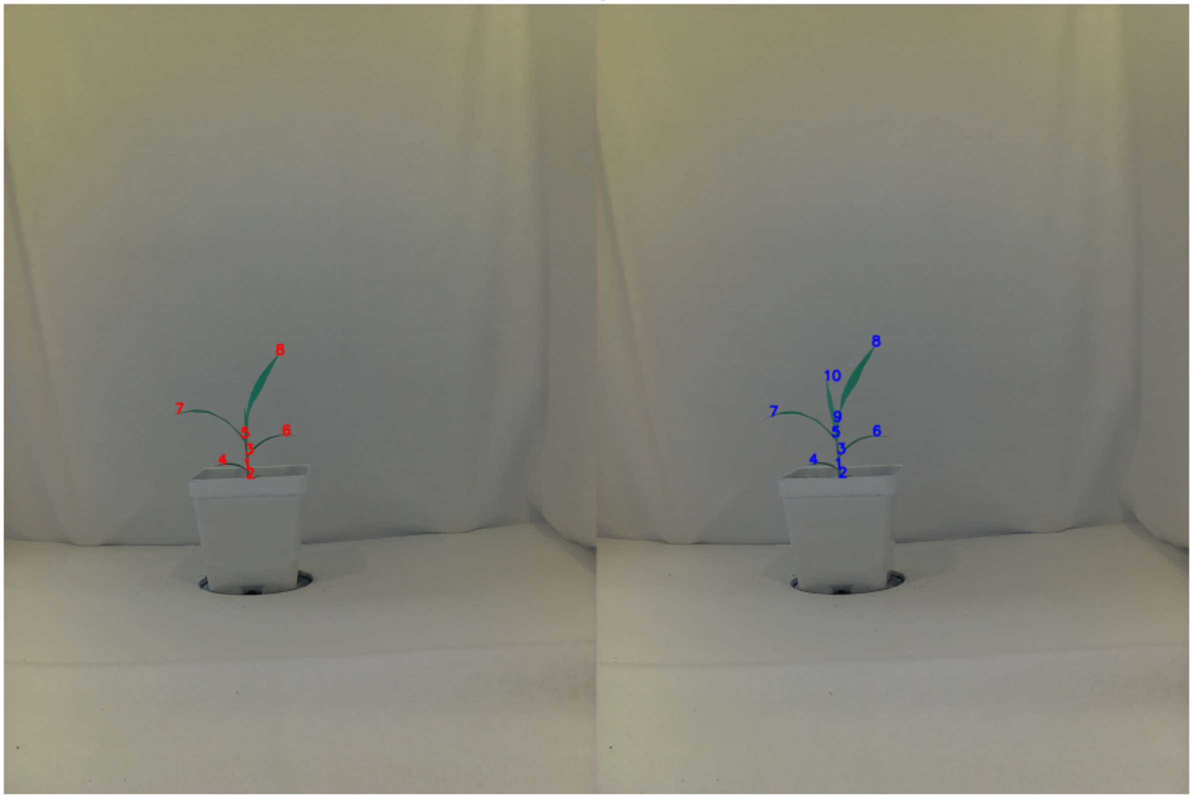

```
Beginning next iteration for days 11 and 12


Running StarScape...

Eigenvalues:  [3.79893075 2.6753918  0.88539829 0.0045825 ]


Var. Explained:  [0.51556917 0.36308889 0.1201612  0.00062191]


Cumul. Var. Explained:  [0.51556917 0.87865806 0.99881926 0.99944117]


2  components sufficiently informative

Running Constella...

20 plms to group
```

B100 day 11-12

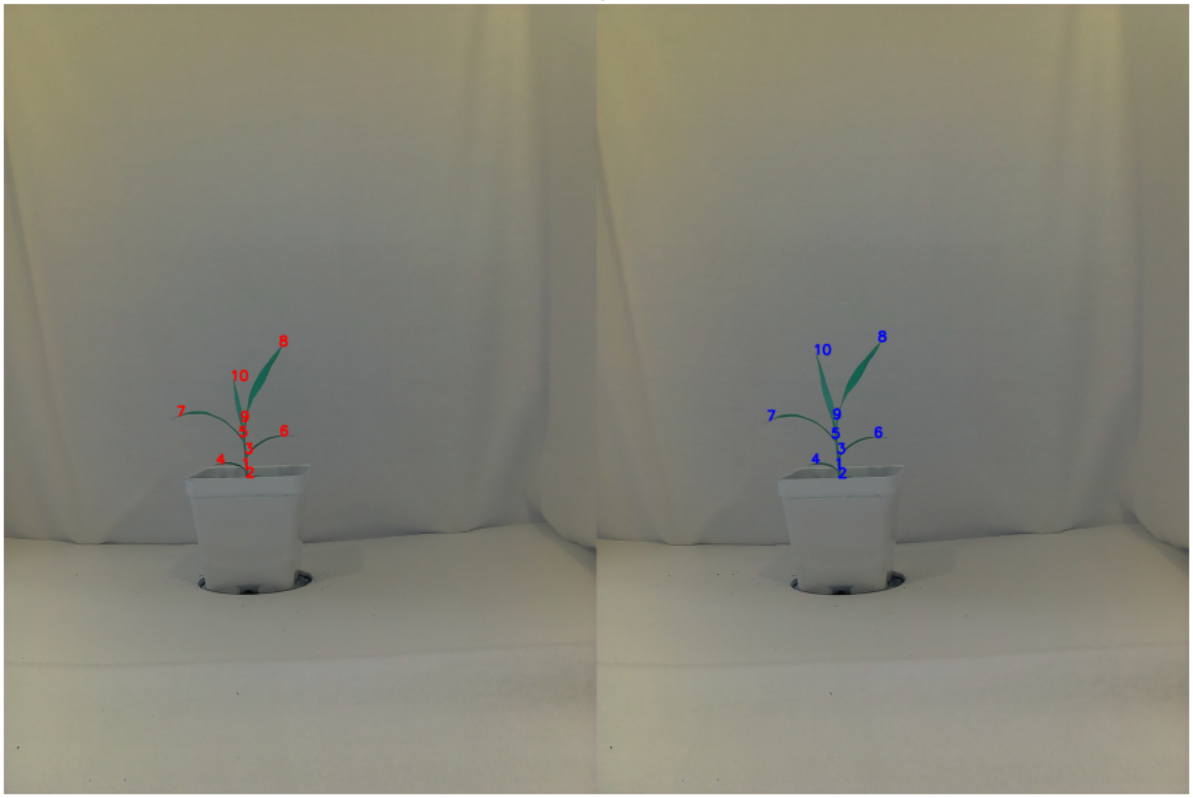

```
Beginning next iteration for days 12 and 13


Running StarScape...

Eigenvalues:  [3.55972863 2.8587077  0.9428435  0.00363656]


Var. Explained:  [0.48310603 0.38796747 0.12795733 0.00049353]


Cumul. Var. Explained:  [0.48310603 0.8710735  0.99903083 0.99952437]


2  components sufficiently informative

Running Constella...

20 plms to group
```

B100 day 12-13

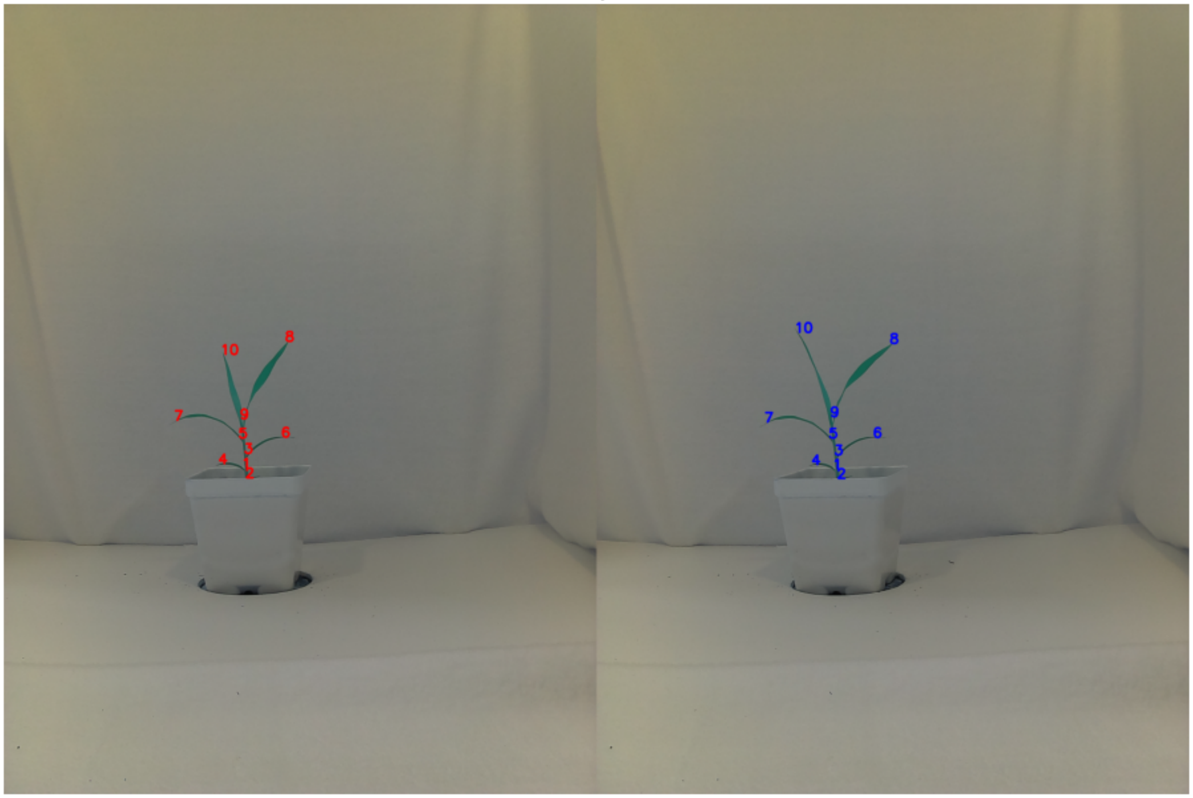


With the homology grouping workflow now completed a decent array of graphical outputs should be visible above displaying not only our PCA related graphs and the dendrogram used for our hierarchical clustering at each step, but also side by side images of our plants as well as their labeled homology groups to enable for easy point of reference for calling accuracy. Let's have a look at our de novo homology groups on our original landmark_pandas dataframe, this time we'll have a look at the full table though instead of just taking a quick glance at the head.

In [5]:

```
landmark_pandas
```

Out[5]:

| | group | plmname | filename | plm_x | plm_y | SS_x | SS_y | TS_x | TS_y | CC_r |
|---|---|---|---|---|---|---|---|---|---|---|
| **0** | 8 | B100_rep1_d10_plm1 | B100_rep1_d10 | 901 | 1151 | 894 | 1171 | 885 | 1167 | 179.786 |
| **1** | 5 | B100_rep1_d10_plm2 | B100_rep1_d10 | 786 | 1423 | 789 | 1402 | 772 | 1404 | 51.358 |
| **2** | 7 | B100_rep1_d10_plm3 | B100_rep1_d10 | 571 | 1344 | 592 | 1336 | 593 | 1342 | 136.758 |
| **3** | 1 | B100_rep1_d10_plm4 | B100_rep1_d10 | 793 | 1523 | 796 | 1512 | 783 | 1519 | 77.395 |
| **4** | 4 | B100_rep1_d10_plm5 | B100_rep1_d10 | 712 | 1511 | 735 | 1508 | 734 | 1512 | 165.041 |
| **5** | 2 | B100_rep1_d10_plm6 | B100_rep1_d10 | 803 | 1555 | 796 | 1538 | 807 | 1535 | 205.491 |
| **6** | 6 | B100_rep1_d10_plm7 | B100_rep1_d10 | 922 | 1415 | 898 | 1420 | 900 | 1416 | 133.183 |
| **7** | 3 | B100_rep1_d10_plm8 | B100_rep1_d10 | 800 | 1477 | 816 | 1459 | 801 | 1454 | 48.893 |
| **8** | 8 | B100_rep1_d11_plm1 | B100_rep1_d11 | 912 | 1123 | 905 | 1142 | 896 | 1139 | 201.460 |
| **9** | 9 | B100_rep1_d11_plm2 | B100_rep1_d11 | 786 | 1370 | 792 | 1350 | 783 | 1347 | 91.428 |
| **10** | 10 | B100_rep1_d11_plm3 | B100_rep1_d11 | 752 | 1236 | 761 | 1252 | 754 | 1260 | 181.622 |
| **11** | 5 | B100_rep1_d11_plm4 | B100_rep1_d11 | 780 | 1420 | 780 | 1398 | 767 | 1404 | 57.927 |
| **12** | 7 | B100_rep1_d11_plm5 | B100_rep1_d11 | 576 | 1353 | 595 | 1343 | 595 | 1350 | 153.015 |
| **13** | 1 | B100_rep1_d11_plm6 | B100_rep1_d11 | 789 | 1523 | 791 | 1511 | 780 | 1517 | 61.134 |
| **14** | 4 | B100_rep1_d11_plm7 | B100_rep1_d11 | 706 | 1510 | 728 | 1506 | 728 | 1510 | 136.659 |
| **15** | 2 | B100_rep1_d11_plm8 | B100_rep1_d11 | 802 | 1553 | 791 | 1532 | 801 | 1533 | 206.361 |
| **16** | 6 | B100_rep1_d11_plm9 | B100_rep1_d11 | 914 | 1417 | 893 | 1423 | 891 | 1417 | 145.431 |
| **17** | 3 | B100_rep1_d11_plm10 | B100_rep1_d11 | 798 | 1475 | 807 | 1466 | 795 | 1459 | 57.528 |
| **18** | 8 | B100_rep1_d12_plm1 | B100_rep1_d12 | 932 | 1108 | 925 | 1131 | 919 | 1121 | 200.401 |
| **19** | 9 | B100_rep1_d12_plm2 | B100_rep1_d12 | 784 | 1362 | 793 | 1343 | 780 | 1343 | 51.891 |
| **20** | 10 | B100_rep1_d12_plm3 | B100_rep1_d12 | 720 | 1151 | 730 | 1170 | 724 | 1175 | 176.054 |
| **21** | 5 | B100_rep1_d12_plm4 | B100_rep1_d12 | 779 | 1425 | 779 | 1402 | 763 | 1407 | 43.620 |
| **22** | 7 | B100_rep1_d12_plm5 | B100_rep1_d12 | 568 | 1367 | 585 | 1358 | 591 | 1361 | 140.915 |
| **23** | 1 | B100_rep1_d12_plm6 | B100_rep1_d12 | 790 | 1523 | 789 | 1511 | 778 | 1518 | 67.282 |
| **24** | 4 | B100_rep1_d12_plm7 | B100_rep1_d12 | 713 | 1509 | 736 | 1506 | 719 | 1511 | 162.192 |
| **25** | 2 | B100_rep1_d12_plm8 | B100_rep1_d12 | 801 | 1555 | 790 | 1534 | 801 | 1535 | 221.172 |
| **26** | 6 | B100_rep1_d12_plm9 | B100_rep1_d12 | 919 | 1422 | 897 | 1426 | 895 | 1422 | 130.591 |
| **27** | 3 | B100_rep1_d12_plm10 | B100_rep1_d12 | 797 | 1475 | 807 | 1467 | 793 | 1461 | 53.073 |
| **28** | 10 | B100_rep1_d13_plm1 | B100_rep1_d13 | 659 | 1079 | 673 | 1097 | 670 | 1101 | 140.700 |
| **29** | 5 | B100_rep1_d13_plm2 | B100_rep1_d13 | 771 | 1425 | 770 | 1405 | 753 | 1408 | 60.049 |
| **30** | 7 | B100_rep1_d13_plm3 | B100_rep1_d13 | 560 | 1373 | 581 | 1362 | 581 | 1368 | 164.285 |
| **31** | 1 | B100_rep1_d13_plm4 | B100_rep1_d13 | 784 | 1526 | 784 | 1506 | 771 | 1518 | 54.060 |
| **32** | 4 | B100_rep1_d13_plm5 | B100_rep1_d13 | 714 | 1512 | 736 | 1507 | 736 | 1513 | 137.769 |
| **33** | 2 | B100_rep1_d13_plm6 | B100_rep1_d13 | 798 | 1557 | 784 | 1543 | 796 | 1539 | 210.764 |

| | group | plmname | filename | plm_x | plm_y | SS_x | SS_y | TS_x | TS_y | CC_r |
|---|---|---|---|---|---|---|---|---|---|---|
| **34** | 6 | B100_rep1_d13_plm7 | B100_rep1_d13 | 916 | 1423 | 900 | 1423 | 892 | 1421 | 128.000 |
| **35** | 3 | B100_rep1_d13_plm8 | B100_rep1_d13 | 789 | 1480 | 799 | 1473 | 787 | 1464 | 54.436 |
| **36** | 8 | B100_rep1_d13_plm9 | B100_rep1_d13 | 971 | 1115 | 958 | 1132 | 950 | 1125 | 193.185 |
| **37** | 9 | B100_rep1_d13_plm10 | B100_rep1_d13 | 776 | 1355 | 784 | 1337 | 771 | 1332 | 64.000 |

We see group serial numbers 1-10 repeating once for each frame so it appears things ran pretty well! Moreover, when we glance at the side-by-side images with the serial numbers superimposed onto the original images things look like they're grouping as we'd expect.

However, as with all de novo methods there is the possibility for errors to be introduced which we might miss at a glance. This brings us to a key aspect of our plm workflow when scaling up to a full sized project which is Quality Control (QC) assessment of our de novo homologies. This is often done with a reduced subset of our full dataset in order to give us a general idea of the overall accuracy of our calls. Although there is currently one method of producing input StarScape files to feeding to Constella eventually as other ways to rescale our acute outputs are developed this method can provide a helpful means of comparing what method of metadata generation (plmSpace) and multivariate space transformation (StarScape) is the best for maximizing biologically informative signal. With this being said let's begin by loading in a table of our landmarks which have been annotated to represent the biological structures they represent*.

*Although not seen in this situation it is common practice to denote random plms which don't represent any meaningful features as '-'.

In [6]:
```
landmark_feat_standards = pd.read_csv('/Users/johnhodge/Documents/GitHu
b/Doust-lab-workflows/B100_timeseries_test_plms_annotated.csv')
landmark_feat_standards.head(10)
```

Out[6]:

| | group | plmname | filename | plm_x | plm_y | SS_x | SS_y | TS_x | TS_y | CC_rati |
|---|---|---|---|---|---|---|---|---|---|---|
| **0** | leaf5 | B100_rep1_d10_plm1 | B100_rep1_d10 | 901 | 1151 | 894 | 1171 | 885 | 1167 | 179.78640 |
| **1** | ligule4 | B100_rep1_d10_plm2 | B100_rep1_d10 | 786 | 1423 | 789 | 1402 | 772 | 1404 | 51.35802 |
| **2** | leaf4 | B100_rep1_d10_plm3 | B100_rep1_d10 | 571 | 1344 | 592 | 1336 | 593 | 1342 | 136.75862 |
| **3** | ligule2 | B100_rep1_d10_plm4 | B100_rep1_d10 | 793 | 1523 | 796 | 1512 | 783 | 1519 | 77.39534 |
| **4** | leaf2 | B100_rep1_d10_plm5 | B100_rep1_d10 | 712 | 1511 | 735 | 1508 | 734 | 1512 | 165.04166 |
| **5** | base | B100_rep1_d10_plm6 | B100_rep1_d10 | 803 | 1555 | 796 | 1538 | 807 | 1535 | 205.49152 |
| **6** | leaf3 | B100_rep1_d10_plm7 | B100_rep1_d10 | 922 | 1415 | 898 | 1420 | 900 | 1416 | 133.18367 |
| **7** | ligule3 | B100_rep1_d10_plm8 | B100_rep1_d10 | 800 | 1477 | 816 | 1459 | 801 | 1454 | 48.89325 |
| **8** | leaf5 | B100_rep1_d11_plm1 | B100_rep1_d11 | 912 | 1123 | 905 | 1142 | 896 | 1139 | 201.46078 |
| **9** | ligule5 | B100_rep1_d11_plm2 | B100_rep1_d11 | 786 | 1370 | 792 | 1350 | 783 | 1347 | 91.42857 |

After glancing at the table above we essentially have 3 types of features we're classifying, our leaf tips denoted as 'leaf', our axils where leaf blades attach to the stem as 'ligule' (common term for this feature in grasses), and 'base' which represents the bottom landmark at the base of our plant. Now we have a list of known features which we can compare to our corresponding list of our predicted homology groups.

In [7]:
```
constellaqc(landmark_pandas, landmark_feat_standards, debug)
```

```
Known Feature-Predicted Group Scoring Matrix:

          1   2   3   4   5   6   7   8   9   10
base      0   4   0   0   0   0   0   0   0   0
leaf2     0   0   0   4   0   0   0   0   0   0
leaf3     0   0   0   0   0   4   0   0   0   0
leaf4     0   0   0   0   0   0   4   0   0   0
leaf5     0   0   0   0   0   0   0   4   0   0
leaf6     0   0   0   0   0   0   0   0   0   3
ligule2   4   0   0   0   0   0   0   0   0   0
ligule3   0   0   4   0   0   0   0   0   0   0
ligule4   0   0   0   0   4   0   0   0   0   0
ligule5   0   0   0   0   0   0   0   0   3   0


Valid Call Rate:      100.0 %
Splitting Call Rate:  0.0 %
Clumping Call Rate:   0.0 %
```

And there we have it! As expected the valid calls were perfect within this tutorial although error, and importantly the type of error is important to keep track of when developing this workflow for your own research. To provide a bit of context let's discuss what our two sources of error represent.

#### Splitting Error

Splitting errors are essentially calls in which more than one de novo homology group was generated to represent a single, known, feature. Within this workflow these errors are often considered less egregious given that they can easily be reconciled together during manual curation of homology groups prior to using plm homology groups for morphometric analyses. A good analogy to this problem is that of scaffold generation during whole genome sequencing in which often only fragments of rather than complete chromosomes are reconstructed from the data. This issue is easily reconciled by a user specifying that these two scaffolds belong together and manually assigning linkage based on known attributes of this data which exist beyond the capacity of the de novo assembler. In a similar vein of logic, if a leaf tip is broken into two groups it can easily be tied together as these groups are given a biologically relevant name.

#### Clumping Error

Clumping errors by contrast are calls in which multiple known features are linked together under a single de novo homology group. Understandably this error is considered far more troubling and all efforts in designing this workflow have been to drive this error rate as low as possible (in most cases hovering in the 5% range for true experimental data). Often datasets which have a high degree of parallax (possessing perspective related artifacts of compressing 3-dimensional structures into a 2-dimensional frame) tend to drive up this error rate. It is often best to check this error rate under a reduced dataset of each genotype or environmental treatment that is anticipated to be used given that it can provide a user with an overall grasp of how well Constella's de novo assignments work within this pool of the data. Cases in which clumping error rates are higher may require a more stringent round of manual curration in order to ensure that morphometric analyses performed on this data afterwards are meaningful.

### In Conclusion

And there you have it! We've successfully started with a handful of time series images and learned how to prepare binary masks which can be used for acute in Tutorial 1. We then learned in Tutorial 2 how to scale up what we had learned in our first exercise to work on batch image datasets. Following the generation of this batch plm data we were then able to explore de novo homology grouping through the use of our StarScape & Constella pipeline in Tutorial 3. And finally in our last exercise we again scaled up what we learned for homology grouping on to batch datasets then were able to test what we generated using ConstellaQC in order to get a general idea of how much confidence we can have in our calls.

I hope this tutorial series have been informative and provides you with some quick-start code to get your own projects running. Cheers!

```
-JGH
```

In [ ]:
```

```
